# Single neurons detect spatiotemporal activity transitions through STP and EI imbalance

**DOI:** 10.1101/2024.10.30.621034

**Authors:** Aditya Asopa, Upinder Singh Bhalla

## Abstract

Sensory input and internal context converge onto the hippocampus as spatio-temporal activity patterns. Transitions in these input patterns are frequently salient. We demonstrate that short-term potentiation (STP) mediates escape from EI balance to implement mismatch detection in spatiotemporally patterned activity sequences. We characterized STP in the mouse hippocampus CA3-CA1 network using optogenetic patterned stimuli in CA3 while recording from CA1 pyramidal neurons. STP modulates EI summation across patterns, first amplifying, then reducing responses. We parameterized a multiscale model of network projections onto hundreds of E and I boutons on a CA1 neuron, each including stochastic signaling to mediate STP. The model detected mismatches in trains of input patterns, which we experimentally confirmed. Mismatch selectivity depends on stimulus overlap, network weights, and connectivity. It is robust over a wide range of model parameters and assumptions about input spike timing jitter, postsynaptic spiking and stochasticity. Finally, we predict that optimal mismatch selectivity can be tuned over low to high gamma frequencies by modulating network parameters, and show that there is strong mismatch detection for gamma-frequency bursts between theta cycles, consistent with theta-tuned snapshots of novel input.

## Introduction

The balance between excitatory and inhibitory drives (EI balance) in neural networks has a wide range of computational functions. These include increased storage capacity (Kanamaru et al. 2023), optimal coding (Denève and Machens 2016), fast temporal encoding and gating (Kremkow et al. 2010; van Vreeswijk and Sompolinsky 1996; Tim P Vogels and Abbott 2009), and higher dynamic range (Bhatia et al. 2019; Wehr and Zador 2003). EI balance applies over tight time scales and even combinations of inputs at a single neuron level, termed precise balance (Bhatia et al. 2019).

Homeostasis plays a crucial role in maintaining EI balance over time-scales ranging from minutes to days (Wen and Turrigiano 2024). Multiple forms of plasticity converge to achieve this balance, including Hebbian plasticity, disinhibition, and homeostatic plasticity (Nelson and Turrigiano 2008). At still shorter time-scales short-term plasticity (STP) mechanisms come into play. These mediate a dual role: they contribute to rapid firing probability changes as part of ongoing computations (Anwar et al. 2017), but they must also maintain network stability even as they transmit bursts of activity.

There is an extensive body of work on fast EI dynamics. Pouille and Scanziani (F. Pouille and Scanziani 2001) used minimal stimuli to show that spikes could be elicited in the narrow interval between arrival of E and I inputs in the CA3->CA1 feedforward circuit. Feedforward inhibition expands dynamic range both in cortex (Frédéric Pouille et al. 2009) and in hippocampus (Bhatia et al. 2019). Tight EI balance has also been shown to track input signals and alter the amplitude of successive gamma cycles even in the absence of plasticity processes (Atallah and Scanziani 2009).

Active synapses undergo STP, during which neurotransmitter release varies upon receiving successive action potentials at the boutons (Fortune and Rose 2001; Regehr 2012; Tsodyks and Markram 1997; Zucker and Regehr 2002). STP emerges from the interaction and competition between pre-synaptic processes of vesicle depletion, calcium influx, calcium build-up, and neurotransmitter reuptake. Together, these processes create a temporal filter between activity in presynaptic and postsynaptic neurons and networks (Fortune and Rose 2001; Klyachko and Stevens 2006a; Galarreta and Hestrin 1998).

With the incorporation of STP, EI balance becomes even more dynamic. In cortical slices, E synapses depress more strongly than I during sustained stimulus trains, thus bounding postsynaptic activity (Galarreta and Hestrin 1998). In contrast, burst stimuli on Schaffer collaterals in rat hippocampal slices have been reported to lead to STD on inhibitory inputs and STP on excitatory ones, leading to a switch-like sharp increase in excitation during bursts (Klyachko and Stevens 2006a).

While most studies on synaptic plasticity utilize 2mM external Ca^2+^ (extCa) this is substantially higher than physiological Ca^2+^ (1.2 to 1.5 mM) and has been shown to significantly alter plasticity outcomes. Long-term plasticity is particularly sensitive to elevated extCa levels (Inglebert et al. 2020). There are significant but smaller changes in short-term plasticity in several systems, wherein lower extCa leads to a) higher paired pulse ratio (PPR) and b) lower EPSCs (Rat hippocampal culture: (Sippy et al. 2003), Calyx of Held: (Thanawala and Regehr 2013; Bernal-Correa et al. 2026; Lorteije et al. 2009), Mouse hippocampal slice: (Blatow et al. 2003)). Specifically, in hippocampal CA3->CA1 glutamatergic synapses at 1.5 mM extCa the paired-pulse ratio is 15% larger than at 2mM (Blatow et al. 2003). In primary hippocampal neuronal culture, inhibitory synapses also undergo a 50% increase in paired-pulse ratio at 0.5 mM compared to 2 mM extCa (Kravchenko et al. 2006). By linear extrapolation/interpolation both these results suggest a PPR increase of ∼20% at 1.35 mM extCa compared to the conventional 2 mM. In the current study we extend our simulations into the physiological domain of extCa to check consistency of the predicted computational outcomes.

Although most plasticity studies focus on the time-domain, network computation engages many neurons in spatially patterned activity. Pattern memories are predicted to coexist with network-scale homeostasis through EI balance that combines plasticity with spatial patterns of activity (T. P. Vogels et al. 2011). Activity in the CA3 converges onto ensembles of cells representing distinct stimuli and contexts (Howard Eichenbaum et al. 1999; Yuste et al. 2024). In abstract terms, these ensembles form instantaneous patterns, or vectors of activity, which are decoded in a heteroassociative manner by the CA1 (Rolls and Kesner 2006).

A static view of such pattern decoding implies differential responses by individual CA1 neurons to different patterns, in other words, each pattern would result in a distinct firing rate. Given that activity is not static, it is more salient for neurons to detect transitions between input patterns. A classical expression of this computation is mismatch negativity (Butler 1968). While the mechanisms for this remain debated, at least one of them involves short-term synaptic adaptation (Garrido et al. 2009).

The current study integrates several research themes of EI balance, short-term plasticity, and network computation to systematically characterize and model the properties of a network with feedforward inhibition. We complete the experiment-model-prediction-testing loop and show that differential changes on E and I synapses may provide a mechanism for single neurons to extract salient features of spatiotemporal inputs through STP (Asopa and Bhalla 2023), while keeping mean activity steady.

We used in vitro whole cell patch clamp recordings and patterned optical stimulation of the CA3-CA1 in the mouse hippocampus, to probe how EI balance evolves over spatiotemporal input trains at CA3. We incorporated this STP data into mass-action stochastic models of presynaptic release for E and I synapses, and embedded a CA1 neuron receiving hundreds of such synapses into an abstract model of the hippocampal FFEI network. This model predicted that the postsynaptic CA1 neuron could detect pattern changes (mismatches), which we confirmed experimentally. We showed that mismatch detection works for pattern transitions and oddball patterns in spiking neurons and in the presence of gamma-frequency modulation of theta bursts. Finally, we show that this computation is robust and can be modulated over a wide range of network parameters.

## Results

### Optogenetic patterned stimulation in the CA3-CA1 network

We obtained whole-cell patch-clamp recordings from mouse CA1 pyramidal cells (PCs) in 350-micron mouse hippocampal slices, while stimulating channelrhodopsin (ChR2) expressing CA3 pyramidal cells using optical patterns generated by a digital micromirror device (Figure 1A-B, see Methods). Our recordings were performed at 32-33 ℃ which has been shown in rats to yield similar short-term plasticity properties as at physiological temperatures (Klyachko and Stevens 2006b). Our bath solution had 2mM Ca^2+^, which is above the physiological 1.2 to 1.5 mM range and hence is likely to exhibit ∼20% lower STP both for inhibitory and excitatory synapses (Kravchenko et al. 2006; Blatow et al. 2003). A pattern consisted of either 1, 5, or 15 spots (spot size 13 x 7 µm, power 14.5 µW/spot) of blue light (470 nm). Due to the grid pattern of illumination and the rectangular profile of the illumination spots, we refer to them as ‘squares’ in subsequent analysis, though the exact shape is unlikely to affect the activation of CA3 neurons. We illuminated the CA3 layer in a series of eight pulses at a fixed frequency of 20, 30, 40, and 50 Hz (Figure 1D). A probe pulse preceded this train by 300 ms to provide a reference baseline response. We measured the dynamics of E-I balance in a three-dimensional stimulus space consisting of CA3 activation size (number of illumination squares), activation frequency (stimulation frequency), and short-term plasticity dependence (pulse number) in the train (Figure 1C).

**Figure 1:**
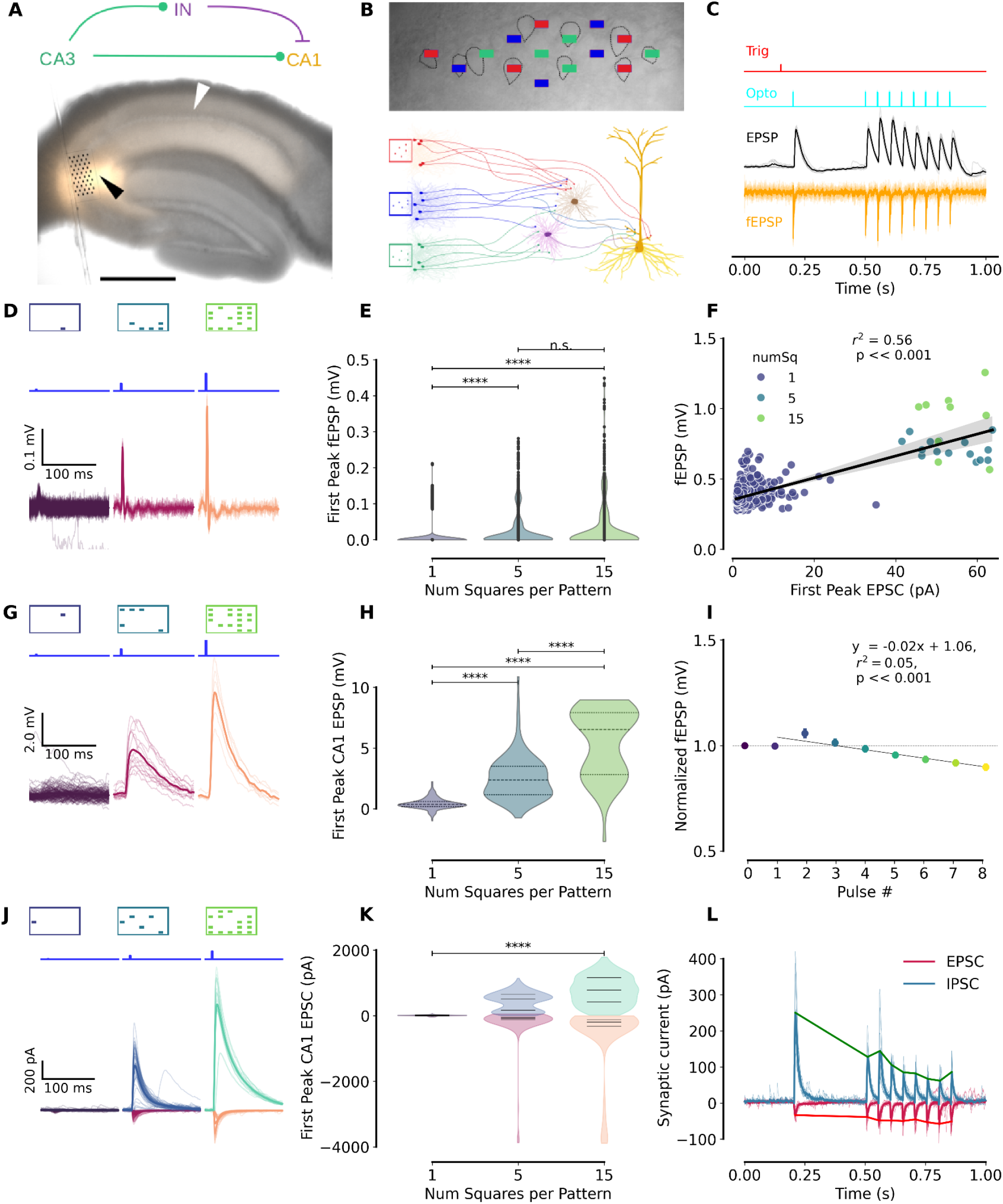
Experiment design and basic response properties. **A.** (Top) Network schematic of hippocampal CA3-IN-CA1 network. (Bottom) A transverse hippocampal section showing channelrhodopsin expression in orange (tdTomato), with the stimulation grid drawn to scale overlaid on the CA3 network. An extracellular (field) electrode was used to record the total optogenetic excitation of the CA3 layer (black arrowhead), and CA1 recordings were made from individual pyramidal cells with a whole-cell patch clamp electrode (white arrowhead). (scale bar = 500µm) **B.** (Top) A DIC image of the CA3 cell layer with a few illumination squares of a stimulation grid overlaid, drawn to scale. (Black dotted lines mark the outlines of a few CA3 cells). (Bottom) A schematic of the CA3-IN-CA1 layer showing the recruitment of CA3 pyramidal cells with three different patterns. The downstream CA1 pyramidal cell receives direct monosynaptic excitation from the activated population and disynaptic inhibition via a heterogeneous, shared population of CA1 interneurons. **C.** Burst recording protocol. Top (red): Trigger to start sequence. Cyan: Optical stimulus. Black: Postsynaptic potential of a patched CA1 cell (black). Orange: Simultaneously recorded field potential (orange). **D.** Extracellular recording of CA3 layer activation for 1,5,15 square patterns. **E.** CA3 field response to first pulse increases with number of squares per pattern (Kruskal-Wallis test). **F.** Correlation of optically stimulated current recorded from patched CA3 cells with the corresponding field response from CA3. **G.** Post synaptic potentials for first pulse (hence, no STP) recorded in a CA1 pyramidal cell for different pattern sizes. **H.** Distribution of PSPs across all recorded CA1 cells differs for number of squares (n=16, p < 0.001, Kruskal-Wallis test). **I.** Channelrhodopsin desensitization during burst is under 10% (linear regression fit with slope 2%, r^2^=0.05, bars indicate 95% CI). **J.** Postsynaptic currents (PSC) recorded for first pulse from CA1 cells in voltage clamp show the proportional relationship between excitatory and inhibitory PSCs for each pattern size. **K.** Distribution of PSC amplitude differs between 1 and 15 square stimuli. **L.** A sample recording of excitatory (pink) and inhibitory (teal) currents recorded from a CA1 cell in the burst protocol described in panel C, for a 15 square pattern.

To monitor the strength and consistency of the total resultant optogenetic activation of the CA3 layer, we used an extracellular field electrode in the CA3 stratum radiatum (Figure 1A, methods). The field response correlated well with optically-driven CA1 PC depolarization (Figure 1E-G), and scaled with the size of the pattern (Figure 1F). This was also consistent with the observation of a wide field of excitability around individual CA3 neurons (Bhatia et al. 2019) (Figure 1-figure supplement 1). From this we expect that there is some overlap in the sets of CA3 neurons activated by different patterns, and this overlap increases with more stimulus squares. Notably, the distribution of field amplitudes was tight (Figure 1E, Figure 1-figure supplement 2F), more so than the distribution of the corresponding EPSPs (Figure 1H). Together with previous work using a similar optical stimulus system and intracellular CA3 recordings (Bhatia et al. 2019) we interpret this to say that the spiking responses from CA3 neurons to optical stimuli were consistent from trial to trial. The field response showed a slight decrease over the course of the 8-pulse train of approximately 2% per pulse (regression fit slope=0.02, r^2^=0.05). We attribute this to ChR2 desensitization, however in longer pulse trains this levels out as we discuss below. We observed a small amount of ‘ringing’ of the field response which we interpret as either CA3 spiking in a burst, or recurrent activation of the CA3 neurons (Figure 1-figure supplement 2). The second peak was ∼40% of the first, within 4 ms of the first. The ringing was down to ∼5% by the third peak which occurred within ∼9 ms of the flash. This sets an upper bound to any contribution to patterns by recurrence or multiple spikes during a ChR2 flash.

On the basis of these field responses and previous reports (Bhatia et al. 2019), we infer that the CA3 neuron response to optical input was a brief event, with reliable probability of a spike on any given flash, and typically having one or occasionally two spikes within 6 ms of the light flash.

As expected, patterns with more illumination squares elicited larger responses in CA1 in current clamp (Figure 1 H-I) and voltage clamp (Figure 1 K-L). We separated E and I components by holding the cell at -70 mV (GABAR1 reversal potential) and 0 mV (gluR reversal potential) respectively. As reported previously (Bhatia et al. 2019), different activation patterns in CA3 produced proportional E and I post-synaptic currents in the recorded neuron (Figure 1K). However, over the pulse train, E and I underwent distinct STP profiles (Figure 1 M).

Thus our patterned optical stimulation provided multiple input combinations to probe the dynamics of monosynaptic excitation and disynaptic inhibition arriving at CA1 pyramidal cells.

### EI balance evolves over the pulse train

We next asked how precise EI balance at the level of CA1 PCs dynamically evolves due to short-term plasticity processes across a range of frequencies and input strengths (Figure 2A). All the PSPs of an 8-pulse train were normalised to the probe pulse. The PSP varied significantly on all the tested three axes of parameters: pattern size (p<0.001), stimulus frequency (p<0.001), and pulse index (p<0.001) (ANOVA). Notably, the PSP of the first four pulses was significantly larger than that in the last four. We tested this by comparing the pulse responses divided between the two halves of the train against a shuffled pulse order (p<0.001, Wilcoxon rank test).

**Figure 2.**
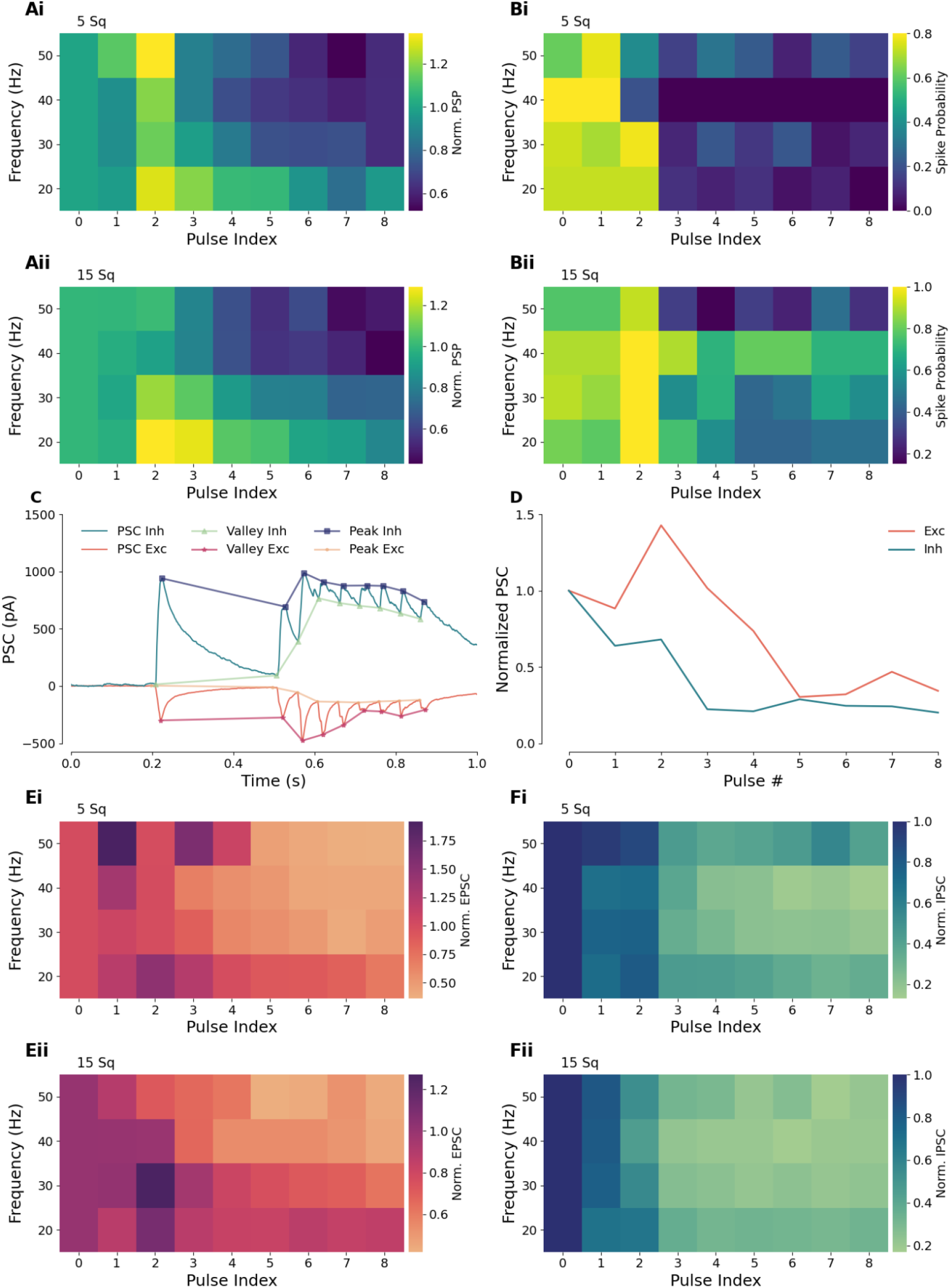
PSP, spiking, EPSC and IPSC dependence on stimulus frequency over a train of pulses. (A) Trial-normalised excitatory postsynaptic potentials across stimulation frequency and pulse index in the train for 5 square (Ai) and 15 square (Aii) patterns across all recorded cells (n=16). B. Spike probability as a function of the stimulus frequency, pulse index and number of squares in the current clamp cells showing higher likelihood in the first half of the train for five squares (top) and 15 square patterns (bottom). C. Schematic for derivation of Post-synaptic responses. Kernel fits were obtained to valley to peak height for each response for both postsynaptic currents (voltage clamp, EPSC and IPSCs) and postsynaptic voltage (current clamp, not shown). D. Normalised EPSC and IPSC responses obtained for an example cell for a sample pattern showing consistent depression in inhibition and biphasic response in excitation. E, F. Both excitatory (Ei-Eii) and inhibitory (Fi-Fii) PSCs show short-term depression along the pulse train. Traces were normalised to a reference pulse 0, delivered 300ms before the burst.

While PSPs are a measure of subthreshold EI balance dynamics, neuronal firing is the network readout of escape from EI balance. Spiking was rare for the stimulus strengths in our experiments. We observed spiking in only 7% of trials (n=154/2201). We measured the likelihood of escape from EI balance by counting the fraction of total spiking trials that elicited for each combination of parameters (Figure 2B). Similar to the PSP response amplitudes, spike likelihood was higher in the first half of the pulse train (p < 0.001, ANOVA) Similarly, the spike likelihood depended significantly on both the pulse train frequency (p<0.001, ANOVA) and on the number of squares in the stimulus (p << 0.001, ANOVA).

We next measured E and I currents under voltage clamp across pattern size, pulse frequency, and pulse number. The EPSCs showed a trend of early potentiation followed by depression (Figure 2D, 2E), while the IPSCs underwent depression from the start (Figure 2 D, F). For the EPSCs, the 15-square trials had a higher reference pulse and higher stimulus overlap (discussed below), hence their normalised peak values were smaller (Figure 2E).

Thus EI balance over a burst briefly tilts in favour of E, after which both E and I undergo depression. This may result in a brief opportunity of escape from EI balance two or three pulses into a burst.

### Multi-synapse summation and divisive normalization evolve over a pulse train

How might a neuron sum distinct input ensembles if they have distinct histories? We used the pattern-size axis of our stimulus protocol to probe history dependence of summation and subthreshold divisive normalization. We have previously shown that summed PSPs are not direct outcomes of EI ratios, because the delay between E and I onset is also a function of stimulus strength (Bhatia et al. 2019). We therefore measured each of these terms: postsynaptic potentials, postsynaptic excitatory and inhibitory currents, and their onset and peak delays (Figure 3 A-D) over the pulse series, and obtained gamma (γ) as a measure of linearity (Bhatia et al. 2019) (Figure 3 E-G). Gamma (γ) relates the observed response of a CA1 cell to the expected response obtained by summing the inputs linearly in the following manner:

**Figure 3:**
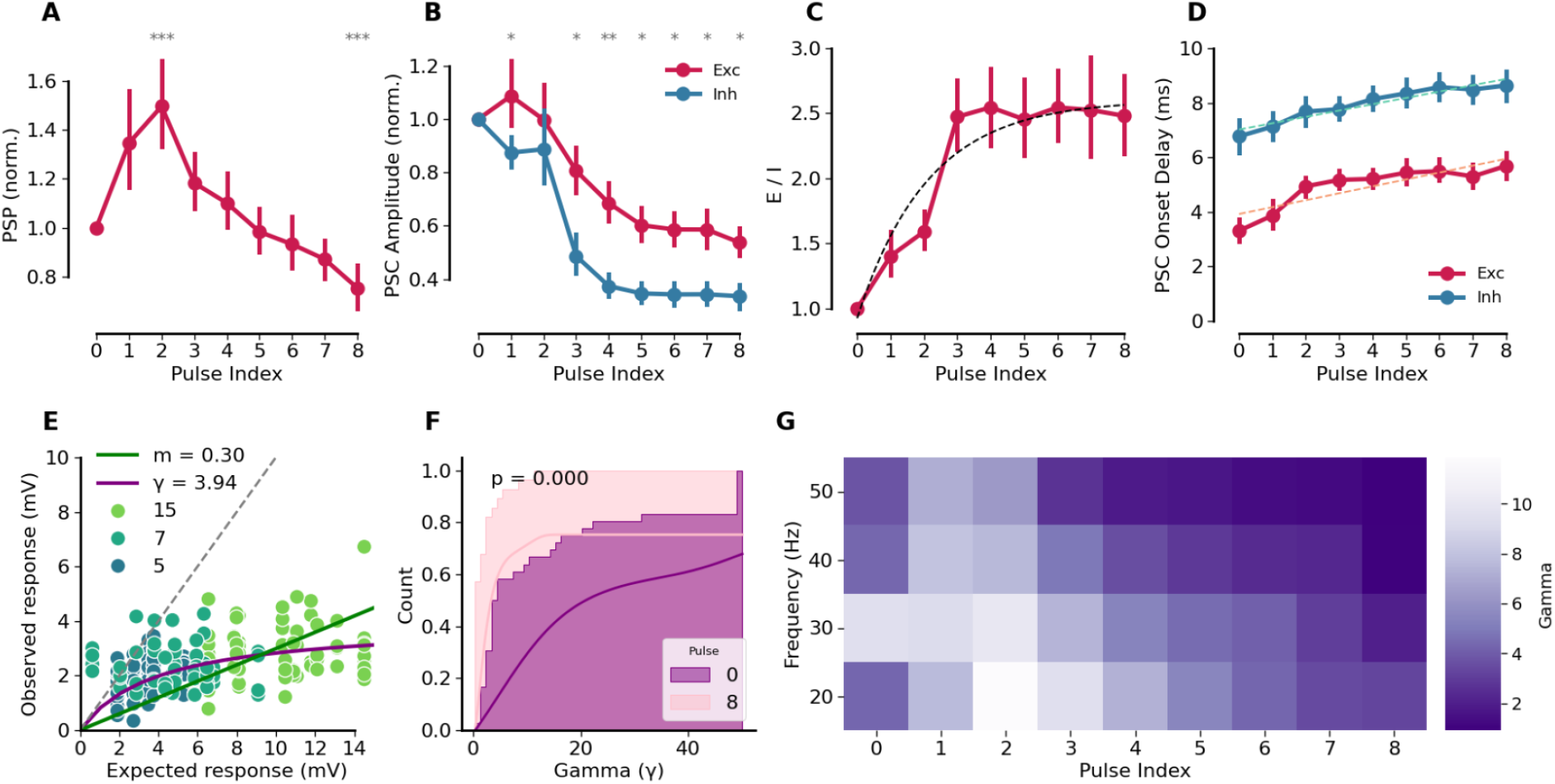
Excitatory-inhibitory balance evolves over the pulse sequence. A. Postsynaptic potentials have a biphasic profile: they first rise, then fall in a pulse train. Pulse 2 > 1.0 at p=0.0003, Pulse 8 < 1.0 at p = 0.0003, Wilcoxon signed rank test. Stimulus was 20 Hz and response is normalized to pulse 0, for all patterns of size 5 and 15. B. Same stimulus as A, for voltage clamp. Postsynaptic excitatory (red) currents first rise slightly, then fall. Inhibitory (teal) currents fall throughout the train. All points except pulse 2 differ at p < 0.05, Holm-Bonferroni corrected Wilcoxon test. C. E-I ratio rises along the pulse train. Dashed line: saturating exponential fit, τ=2.115±1.0 pulses. D. The onset delay for both excitation and inhibition rises steadily over the pulse train. Dashed lines are linear regression fit with slope = 0.253ms/pulse for E and 0.232ms/pulse for I, slope difference not significant at p=0.123). E. Scaling of observed responses in CA1 for patterns of different sizes (5, 7, 15 illuminated squares) as compared to the expected responses obtained by the sum of constituent single-square responses, shown for cell 3402, to illustrate the response normalisation. Solid purple curve is a fit to the equation with gamma = 3.94 = expected*observed/(expected - observed). Solid green line is best fit linear slope, m = 0.30. F. Gamma shows a leftward shift along the pulse train, here shown by comparing cumulative distribution of gamma for probe pulse vs pulse 8 of the train. (p<1e-3, Mann-Whitney test) G. Gamma depends both on pulse index and frequency. Gamma during the first four pulses is larger than in the later four (p<0.001, ANOVA).

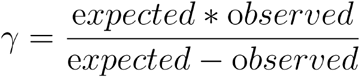

The smaller the gamma, the larger the deviation from the linear combination of inputs due to the divisive effects of inhibition.

By comparing observed vs. expected responses, we replicated earlier observations (Bhatia et al. 2019; Wehr and Zador 2003) showing sublinear summation, and obtained a median gamma of 7.16 (95% CI = 4.76 - 10.2) (Figure 3G). For comparison, we also performed a linear regression and obtained a median slope of 0.57 (95% CI = 0.45 - 0.72). We found that both gamma and slope values show a rise and then a fall in a history-dependent manner (Figure 3G). Thus the differential STP profiles for E and I synapses yield a history-dependence of pattern-specific summation, with greater linearity around pulse two or three in a train.

### Presynaptic release models are bounded by observations from pulse-train sequences

Having characterised the dynamics of plasticity of excitatory and inhibitory synaptic currents over trains of pulses, we next used this data to develop models of presynaptic signalling. Datasets with similar pulse trains have been used to obtain phenomenological models (e.g.,(Barros-Zulaica et al. 2019)). Instead, we chose to build a chemical kinetics-based multi-step neurotransmitter vesicle release model similar to previously published models (Neher 2015; Schneggenburger et al. 1999) (Figure 4A). This incorporated both calcium buffering, which contributes to short-term potentiation, and depletion of docked vesicles, which underlies short-term depression. We implemented a ball-and-stick cellular model having AMPA and GABA synapses on the dendrite, with current or voltage-clamp recordings from the soma to compare to experiments. This chemical-electrical model was implemented in MOOSE (Ray and Bhalla 2008), and parameters were fit using a multi-objective optimization pipeline to match waveforms and dynamics over all frequencies and the entire pulse train. We independently fit presynaptic signaling models for E and I to their respective voltage-clamp waveforms (Methods, Figure 4-figure supplement 1-3) (Nisha Ann Viswan et al. 2024). Each cell had somewhat distinct STP characteristics (Figure 4-figure supplement 4).

**Figure 4:**
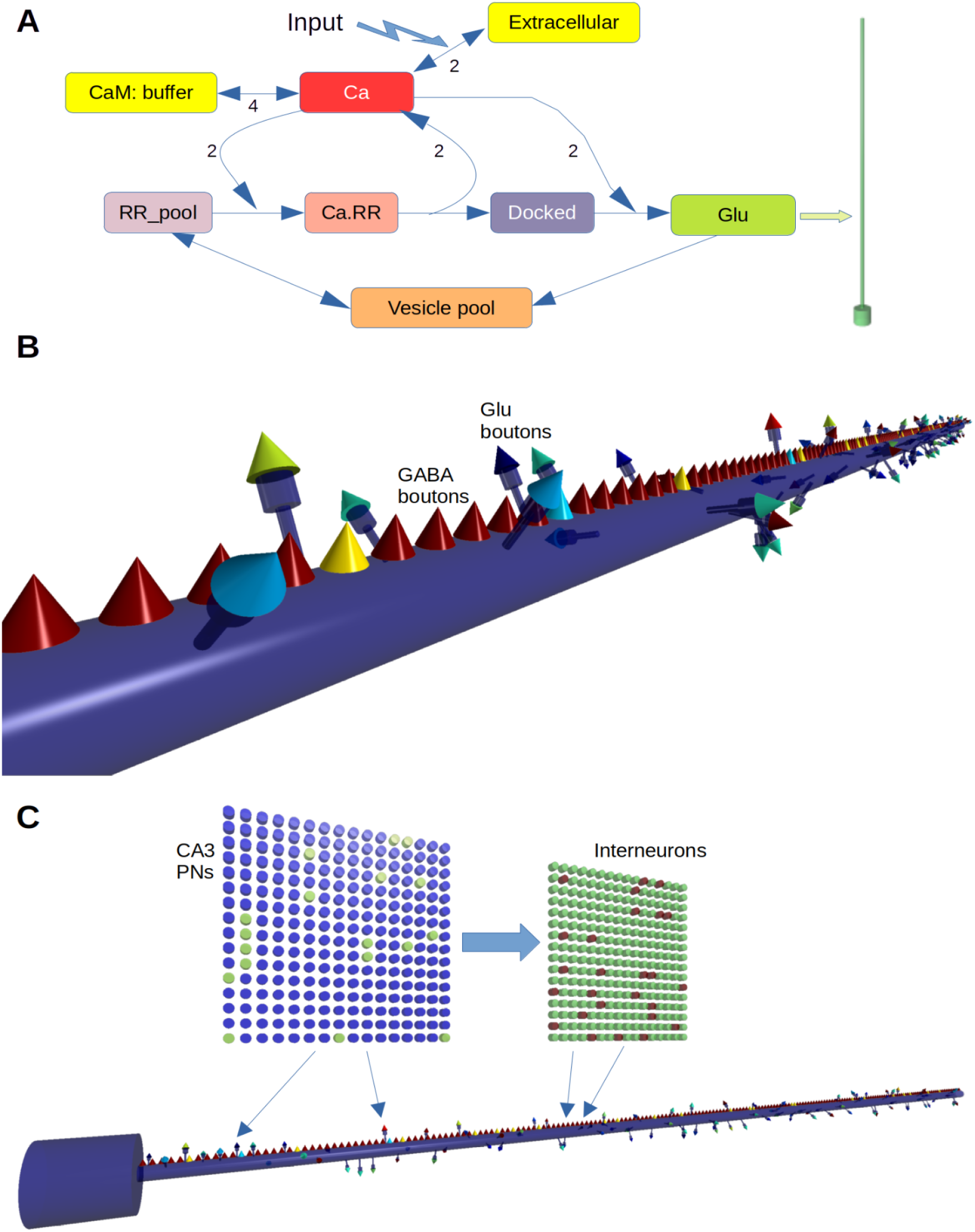
Multiscale model. A: Model of reaction system in each bouton. Synaptic input triggers entry of extracellular Ca2+ into the bouton, and a pump removes the Ca. CaM buffers the Ca2+. Ca binds/unbinds from successive stages of the readily releasable (RR) pool of vesicles till they are docked, at which point the final Ca-binding step causes synaptic release. The released neurotransmitter (Glu for excitatory synapses, GABA for inhibitory) opens a ligand-gated receptor channel on the postsynaptic CA1 neuron, which is held voltage-clamped.(green arrow). The reaction scheme was identical for Glutamatergic and GABAergic synapses, but the rates were different to fit the respective voltage-clamp recordings. B: Close-up of dendrite of CA1 pyramidal neuron model. The cell has a compartment for soma, a 200-micron compartment for the dendrite, and 100 spines. Each spine is modelled as a head compartment, a neck compartment, and a presynaptic glutamatergic bouton. There are 200 GABAergic presynaptic boutons positioned directly on the dendrite. C: Network model. The CA3 has 16x16 neurons, projecting randomly to 16x16 interneurons. CA3 also projects directly to the Glu synapses on the spines. The interneurons project to GABA synapses.

While the E fit applied directly to the CA3->CA1 pyramidal neuron glutamatergic synapse, the I fit was a composite of CA3->Interneuron and Interneuron->CA1 projections (Figure 4B, C). Based on the peak and onset times, we made the further simplifying assumption that only one interneuron class was involved, most likely PV interneurons. An example of CA3-triggered presynaptic signalling and postsynaptic currents is presented in Figure 4-figure supplement 5. Note that the synaptic chemistry was modelled using the Gillespie Stochastic Systems Algorithm (Gillespie 1977), hence the responses are noisy and synaptic release is probabilistic.

We used parameters for Cell 7492 for subsequent model-building (Figure 4-figure supplement 1).

#### Burst EPSP responses constrain single-cell parameters

We next embedded our presynaptic signalling models as synaptic boutons in a conductance-based electrophysiology model of a neuron, and used burst-EPSP responses to constrain its properties. 100 glu synapses were placed on dendritic spines, and 200 GABA synapses on the dendrite (Figure 4B). In the CA1 the E:I synapse ratio is between 12:1 to 25:1 (Bezaire and Soltesz 2013), but we implemented a subset of E synapses targeted by the CA3 pattern-responsive neurons in the interests of computational efficiency, as most of the E inputs would be silent.

We implemented an abstract hippocampal network to drive activity on the synapses. CA3 was implemented as a 16x16 array of integrate-and-fire neurons. Optical patterned input was mapped onto this array (Figure 4-figure supplement 6) and included charge buildup from successive light pulses as well as ChR2 desensitisation (methods, Figure 4-figure supplement 7). The interneuron layer was a 16x16 array of binary neurons. The CA3 array projected sparsely to the 100 glutamatergic synapses on the CA1 model and to the 256 interneurons. The interneuron array projected onto the 200 GABAergic synapses (Figure 4C). We used this full model, with optical stimulus, CA3, Interneurons, CA1 neuron, probabilistic connectivity, and presynaptic signaling chemistry, for all subsequent calculations in this study. We also made a variant of this model adjusted for more physiological conditions at 37℃ and 1.35 mM external Ca^2+^ (methods).

As a first-pass test of our model and comparison with experimental data, we modelled pulse train experiments performed using current clamp. We ran the simulations deterministically to correspond to the averaged experimental traces (Figure 5 Ai-Aiv). There was no significant difference between experiment and simulation distributions for any of the frequencies (Figure 5 E, Table 1), nor for the slope over frequencies (Figure 5Av, Table 1). Since simulations permit access to internal state variables not available to the experiments, we also looked at how E and I currents balanced out during the pulse train (Figure 5B). Notably, the simulations predict that the E current builds up, especially on the second pulse of the train. This corresponds well with the observations in Figure 5 A and in Figures 2 and 3. We additionally compared our experimental traces with the physiologically tuned model, and it too matched experiments closely (Figure 5 C, Table 1).

**Figure 5:**
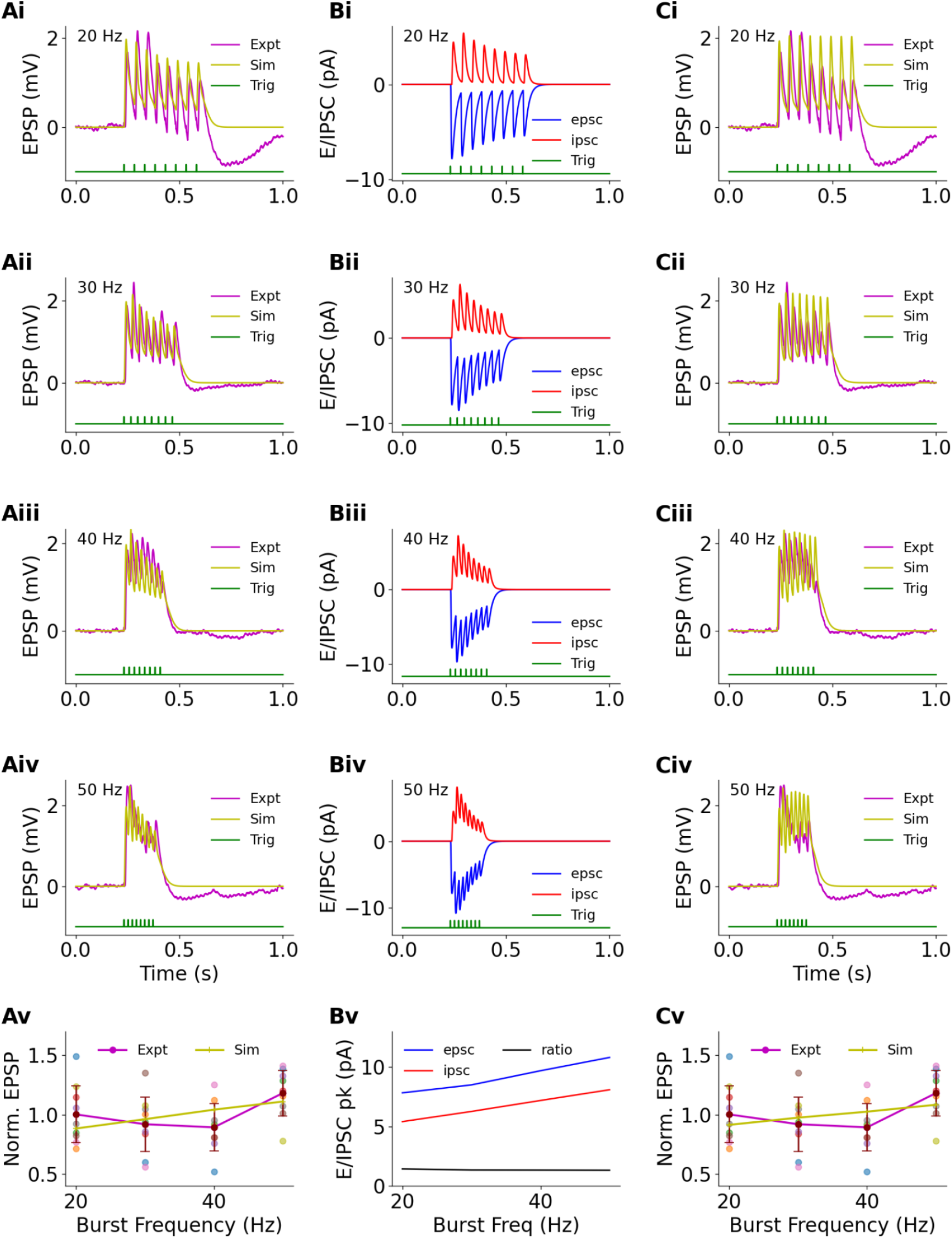
Temporal summation and EI balance in burst stimuli. Ai-Aiv: EPSP measured at soma for representative cell 3402 (maroon) and simulation (yellow) for each frequency. Trigger pulse timings are in green. Av: Distribution and mean of EPSP over all recorded cells for each frequency for the experiment (maroon) and for a simulated neuron (yellow). There is no significant dependence of EPSP on frequency for the recordings (Linear regression, slope=0.005, p=0.168) or for simulations (Linear regression, slope = 0.01, p=0.134). Simulated slope lies within the 95% range of the confidence interval [-0.002, 0.0126] of the slope for experimental data (Bootstrap, 10000 resamples, Table 1). The simulated peak EPSPs lie within the range of experimental cells at each of the four frequencies (Table 1) Bi-Biv: Simulated excitatory and inhibitory currents for each frequency. Note that the currents are measured close to resting potential, so the driving force (E_GABA_ - Vm) is small compared to the voltage-clamp experiments in Figure 1K, where Vm was held at 0 mV. Bv: EI balance (ratio, black line) is maintained for simulated excitatory and inhibitory current peaks for all frequencies. C: Similar to A except the simulation now uses physiological temperature (37℃) and external Ca^2+^ (1.35 mm). The simulated EPSP peaks again lie within the range of experimental cells for all four frequencies.

**Table 1:** Summary of statistical comparisons between simulation and experiment for figures 5 and 6. For Figure 5 panel Av, the simulation EPSP peaks do not differ significantly from experiment either for the means for each individual frequency, nor does the slope across frequencies differ from experiment. For Figure 6 we obtained the specified statistics for each panel, and asked if the simulation was within the range of values seen for individual cells. In all cases it was within the range, except for panel 6F for the 5 Square stimulus.

|  | Figure | Data description | Test | statistic | Comparison | Notes |
| --- | --- | --- | --- | --- | --- | --- |
| 1 | 5Ai-Aiv | Experiment: 9 cells, 4 frequencies.<br>Simulation: 1 run, 4 frequencies | Wilcoxon signed-rank | 20Hz: W=13<br>30Hz: W=18<br>40Hz: W=6<br>50Hz: W=14 | p-value<br>0.301<br>0.652<br>0.055<br>0.359 | No difference at 95% |
| 2 | 5Ai-Aiv | Same | Is simulation in range of experiment<br>[Experiment min, experiment max, **simulation**] | 20 Hz<br>30 Hz<br>40 Hz<br>50 Hz | [0.716, 1.492, **0.884**]<br>[0.563, 1.355, **0.962**]<br>[0.520, 1.253, **1.043**]<br>[0.782, 1.413, **1.111**] | Sim within range at all frequencies |
| 3 | 5Av | Same | Regression fit for slope. Bootstrap for slope at 95% confidence interval to compare simulated vs. experimental | Confidence Interval CI: [-0.002, 0.0126]<br>Sim slope = 0.0076 | 95% confidence interval | Sim is within CI. |
| 4 | 5Ci-Civ | Experiment: 9 cells, 4 frequencies.<br>Simulation: 1 run, 4 frequencies, model@ 37°C and 1.35mM external Ca<sup>2+</sup> | Wilcoxon signed-rank | 20Hz: W=13<br>30Hz: W=18<br>40Hz: W=6<br>50Hz: W=14 | p-value<br>0.496<br>0.496<br>0.129<br>0.164 | No difference at 95% |
| 5 | 5Ci-Civ | Same | Is simulation in range of experiment<br>[Experiment min, experiment max, **simulation**] | 20 Hz<br>30 Hz<br>40 Hz<br>50 Hz | [0.716, 1.492, **0.915**]<br>[0.563, 1.355, **0.975**]<br>[0.520, 1.253, **1.026**]<br>[0.782, 1.413, **1.084**] | Sim within range at all frequencies |
| 6 | 5Cv | Same | Regression fit for | Confidence | 95% | Sim is within |
|  |  |  | slope. Bootstrap for slope at 95% confidence interval to compare simulated vs. experimental | Interval CI: [-0.002, 0.0126]<br>Sim slope = 0.0056 | confidence interval | CI. |
| 7 | 6C | Power spectral density of EPSP due to Poisson train of optical stimuli | Cosine distance between experiment mean and sample: Is simulation within experimental range. | [min, max, **sim**]<br>5 Sq trace:<br>15 Sq trace | [0.933, 0.980, **0.962**]<br>[0.926, 0.990, **0.957**] | Sim within range. |
| 8 | 6D | Probability of spiking following stimulus after initial settling time of 2s | Probability in % | [min, max, **sim**]<br>5 Sq trace<br>15 Sq trace | [28.9, 64.5, **58.7**]<br>[61.9, 90.2, **86.9**] | Sim within range<br>Sim within range |
| 9 | 6E | EPSP vs time: Slope of initial decline for $t < 1.8s$ | Linear regression fit slope range for cells, compared with simulation | [min, max, **sim**]<br>5 Sq trace<br>15 Sq trace | [-1.78, 0.564, **-0.033**]<br>-3.07, 0.329, **-0.124**] | Sim within range<br>Sim within range |
| 10 | 6E | Mean amplitude | Mean amplitude range for cells, compared with simulation | [min, max, **Sim**]<br>5 Sq trace<br>15 Sq trace | [0.395, 1.15, **0.814**]<br>[0.994, 2.88, **1.91**] | Sim within range<br>Sim within range |
| 11 | 6F | Decay time-course of EPSP vs ISI | Time constant tau computed by fitting to data points | [min, max, **sim**]<br>5 Sq trace<br>15 Sq trace | [0.34, 17.6, **34.6**]<br>[8, 933, **49.9**] | Sim **NOT** within range<br>Sim within range |
| 12 | 6G | Fit EPSP amplitude distribution to lognormal | Lognormal mu (log-mean) | [min, max, **sim**]<br>5 Sq trace<br>15 Sq trace | [-1.09, -0.0038, **-0.536**]<br>[-0.139, 0.847, **0.392**] | Sim within range<br>Sim within range |
| 13 | 6G | Fit EPSP amplitude distribution to lognormal | Lognormal sigma (log-std-dev) | [min, max, **sim**]<br>5 Sq trace<br>15 Sq trace | [1.18, 1.39, **1.23**]<br>[0.881, 1.18, **0.99]** | Sim within range<br>Sim within range |

Overall, the synapse and single neuron components of our model worked well together to replicate burst inputs in current clamp.

#### Poisson train stimuli constrain network parameters

Following the synapse and cell-level analysis of signal summation and short-term plasticity in the CA3-CA1 system, we next tuned our model to network-level readouts. Here we explicitly included an analysis of scatter in the experimental data. We modeled the origin of this scatter as stochastic synaptic chemical kinetics and transmitter release (methods). To simulate recording noise we overlaid Gaussian noise with an amplitude of 1 mV and upper cutoff of 200Hz on the simulated EPSP traces. We delivered a ‘frozen’ Poisson-train sequence to experiment and model, to test the capacity of the model to replicate data and to further constrain network-level parameters using a panel of statistics of the responses (Table 1). Figure 6 A and B show example EPSP traces for a single trial of an experiment and simulation, respectively. We used simple parameter sweep runs to tune the model network parameters.

**Figure 6:**
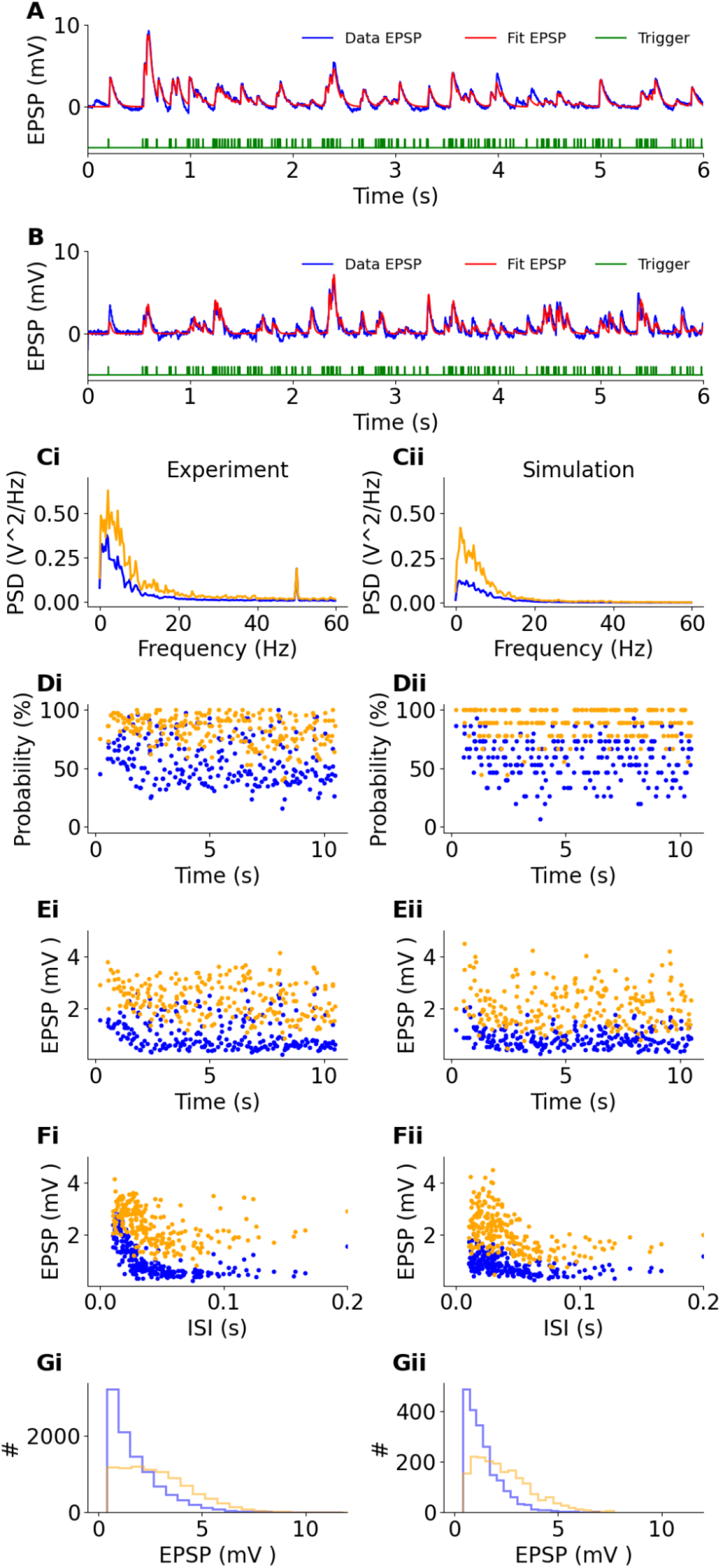
Experimental and simulated responses to Poisson spike train input, all patterns. A: Sample experimental EPSP train aligned with light trigger pulses (green). Blue trace is recorded data; red trace is the fit used to compute peaks and subsequent statistics. B: Same for simulated data. C-G: Comparisons for readouts of experiment (left column) and simulated (right column) data. Blue traces/markers: 5-square stimulation, orange traces/markers: 15-square stimulation. Outcomes are detailed in Table 1. In all cases except panel F-5Sq, the simulated response lies within the range of individual cell responses. C: Power spectral density of EPSP response over the entire dataset. Note 50Hz peak due to line noise in experimental dataset, despite this the cosine distance between experimental mean and simulated response was within the range of individual cell cosine distances to mean. D: probability of trigger to generate a peak in the EPSP trace. E: Scatter plot of EPSP peaks as a function of time over the Poisson train. Note the initial decline due to STP and ChR2 desensitisation. Comparisons were made both for this initial decline (up to 1.8 s) and for the subsequent steady-state EPSP peak distribution. F: Scatter plot of EPSP peaks vs. Inter-spike-interval. The experimental data show an elevation of EPSP at short ISI. We fit this using a simple exponential decay function y=y0.exp(-t/tau)+y1. G: Distribution of EPSP peak amplitudes.

We first estimated the power spectral density of the EPSP waveforms (Figure 6 C, Table 1). We obtained the experimental mean spectral density as a reference, and compared individual cells and simulation to this mean using the cosine distance. The simulation fell within the experimental range of cell cosine distances.

Then we compared the probability that each optical stimulus would elicit an EPSP (Figure 6 D). As expected, 15-square patterns (yellow dots) frequently gave an EPSP (77.5±11.7%), while 5-square patterns failed about half the time (51.4±16%). The simulated runs matched this (Table 1). The probability of failure reduced with increasing volume of the simulated presynaptic boutons, because larger volumes experienced smaller chemical noise (stochasticity) in synaptic release (Figure 6-figure supplement 1). We note that for the purposes of eliciting a postsynaptic response, any unreliability in optical stimulus-triggered firing of the CA3 neuron folds into the probability term for stochastic synaptic release. By matching this metric to experiment, we fine-tuned the scaling term for the presynaptic boutons to 0.2, which after the randomization of synaptic volumes during setup gave a volume of 0.0205±0.0008 femtolitres.

In Figure 6 E we compare scatter plots of EPSP amplitudes over time. Note that the initial high EPSP responses declined over the first two seconds. Hence we made two comparisons between model and experiment: First, the slope of the initial decline (t<1.8s) and then the steady-state distribution (t>2s). The simulations were within the range of experimental cells for both these metrics (Table 1). We asked if ChR2 desensitisation of the stimulus might account for the initial decline. We found that the field response does indeed show a decline over multiple pulses (Figure 6-figure supplement 2). We also checked for short-term depression as a mechanism for this decline by comparing the reference and the no-STP model (Figure 6-figure supplement 3A). Without STP there was no decline, but the absolute EPSPs were considerably elevated.

We then plotted each EPSP against the inter-stimulus interval (ISI) preceding it (Figure 6 F). We observed that the EPSP was high for very short intervals and then declined to a lower steady value, both in experiments and simulations. We initially assumed this was due to increased STF for closely succeeding input pulses. Unexpectedly, the no-STP model also showed the same profile of high EPSP at short ISIs (Figure 6-figure supplement 2B iii). Removal of NMDA receptors in the model also did not change the EPSP-vs-ISI profile (Figure 6-figure supplement 3B iv). Thus, we conclude that the high EPSP was simply due to charge accumulation during rapid synaptic input. For this comparison the simulation was within the experimental range for 15-square stimuli, but not for 5-square stimuli (Table 1).

We finally compared the distributions of EPSP amplitudes (Figure 6 G). We used a log-normal distribution, and thus obtained the log-mean and log-standard deviation terms for each histogram, for each of 5- and 15-square stimuli. In all four comparisons the simulation was within the experimental range (Table 1).

Overall, we were able to quantitatively replicate almost all features of the experimental dataset in our multiscale model incorporating presynaptic signalling, postsynaptic electrophysiology, and abstracted network connectivity and responses. Between the datasets in Figure 4-figure supplements 1 to 3, Figure 5, and Figure 6, we were able to substantially constrain the parameters in our model, from chemical to cellular physiology to network. Network connectivity parameters are presented in Table 2. All subsequent simulations used this ‘tuned’ model except for spiking models, which had 2x greater synaptic weights.

**Table 2:** Network parameters and their definitions.

| Parameter | Description | Default value |
| --- | --- | --- |
| wtGlu | Mean conductance (nS) of population of 100 AMPA synapses. Defined in model as relative value with reference = 5. Open conductance scales this by the release amount of Glu transmitter. | 0.474 ± 0.013 nS |
| wtGABA | Weight (conductance) of each of 200 GABAergic synapses. Relative reference value in model = 10. Open | 12.57 nS |
|  | conductance scales this by the release amount of GABA transmitter. |  |
| pCA3_CA1 | CA3 to CA1 connection probability, i.e., probability that a given CA3 neuron synapses onto a CA1 neuron | 0.02 |
| pCA3_Inter | CA3 to Interneuron connection probability | 0.01 |
| pInter_CA1 | Interneuron to CA1 connection probability | 0.01 |
| zeroIndices | Number of silent CA3 neurons, 'masking' optical input. | 192 |
| ChR2_ampl | Scaling factor for optical stimulus to depolarize CA3 neuron. (Not varied) | 0.1 |

### STP and sparse coding yield pattern mismatch detection

We next used our model to ask how an important neural computation, that of mismatch detection between successive pulses of network activity, could emerge from single-cell computation involving EI balance and STP. The existing heuristic argument based on synaptic adaptation(May and Tiitinen 2010) is as follows: Consider two patterns of activity, A and B, comprising different ensembles of CA3 pyramidal cells. Let these overlap to a small extent, say AՈB < 20%. Since the CA3 projects directly to CA1 spines, the overlap onto excitatory synapses will also be around AՈB. Now, we consider STP. Repeated inputs of pattern A result in the initial strengthening of excitatory inputs, then depression. At the transition from pattern A to B, most of the excitatory inputs have not been stimulated before, so the fresh synapses are stronger. This gives a larger excitatory drive. From this heuristic argument, we predict two things: First, at transitions between patterns (mismatches), we expect a transient elevated response in the postsynaptic cell. Second, the size of the transient should be larger if the input patterns are sparse in E, because this lessens the overlap between them. We note that our model also includes plasticity in inhibitory synapses, which has implications for transient changes in EI balance as analyzed below.

To test the prediction of mismatch detection in the model, we simulated a sequence of patterns of the form “AAAAAAAABBBBBBBBCCCCCCCCDDDDDDDD”, that is, eight repeats of each of four spatial patterns shaped like the letters A to D (Figure 4-figure supplement 6). As expected from the above argument, EPSCs experienced an elevated response just after the pattern change (mismatch detection), and this was dependent on the repeat frequency of the patterns (Figure 7A). IPSCs, on the other hand, started with a large response and then underwent a nearly smooth decline over the repeated pulses, with very little alteration at the pattern changes (Figure 7A). This smooth decline arises from two factors: The observed short-term depression of the inhibitory inputs (Figure 3B), and the high degree of overlap on inhibitory inputs because of the two-stage inhibitory feedforward connections (Figure 7-figure supplement 1). Put together, these E and I components give both strong mismatch detection (due to sparse E) and delayed EI balance due to compensation for the slow STD of E synapses by corresponding STD of I synapses. In current-clamp mode, this gives us consistent mismatch detection (Figure 7B). When all synaptic and channel kinetics were scaled to physiological temperature (37℃) the mismatch transients were larger than the reference simulations (Figure 7C). Conversely, when external Ca^2+^ was reduced from the experimental 2 mM to a more physiological 1.35 mM, the mismatch transients were smaller but still distinct (Figure 7D).

**Figure 7:**
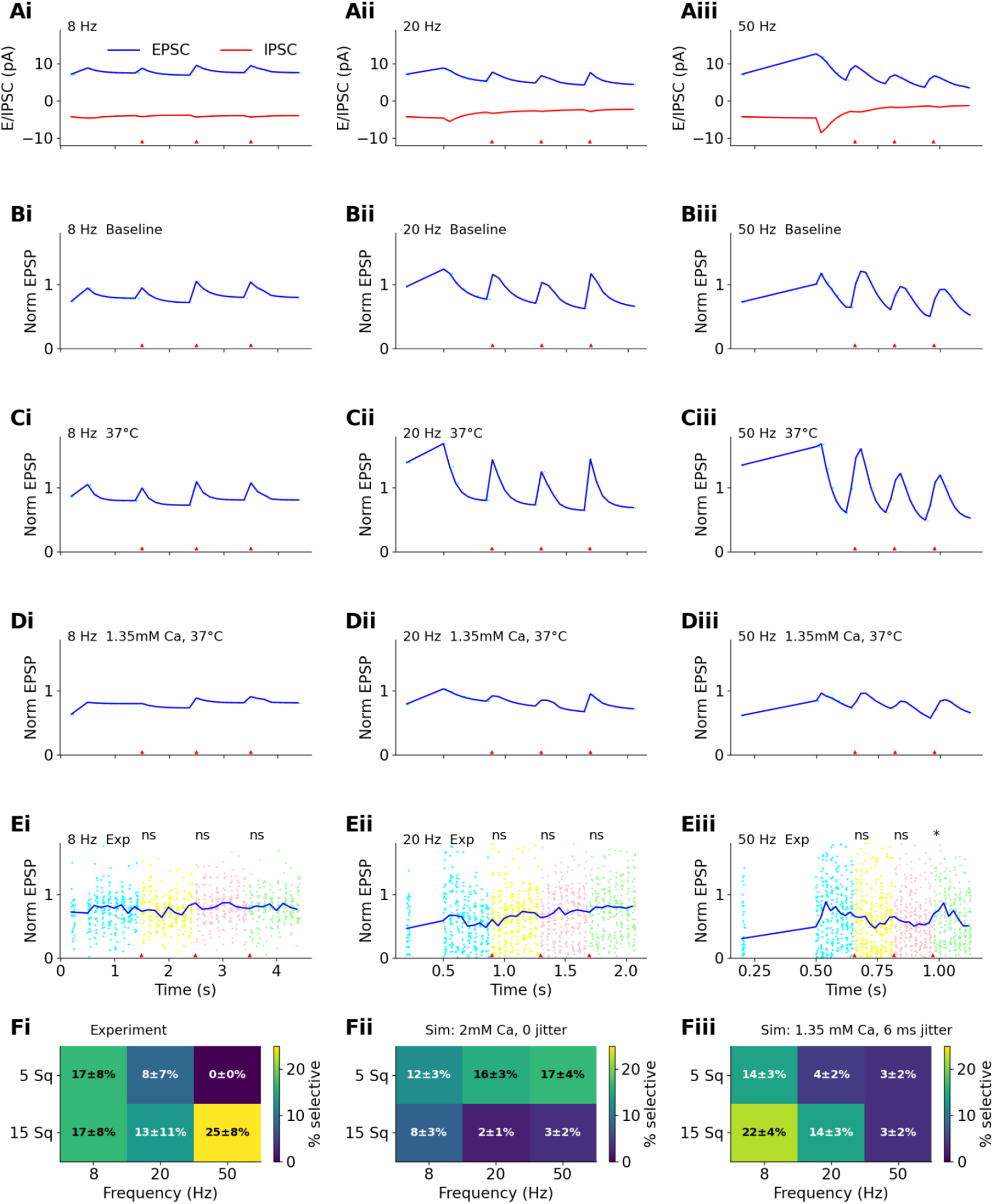
Mismatch detection in simulations and experiment. In all cases, the stimulus was 8 repeats each of 4 patterns, at the specified frequency. In panels A-E the red triangles indicate time of pattern change. A: Deterministic simulations. Excitatory and inhibitory currents contribute different terms to the mismatch signal. Excitatory currents detect the transition, whereas inhibitory currents undergo synaptic depression to balance depression in excitatory currents. B: Same simulations. At each mismatch there is an elevation of EPSP, shown normalized around the mean signal. C: Same as B except using the model configured for 37° C. D: Simulations with 1.35mM external Ca at 37° C show weaker but still distinct mismatch detection. E: Experimental data. Colored scatter points indicate individual trials, the different colors represent different patterns. The solid blue line is the mean. A significant (p=0.013, Wilcoxon signed-rank left-tailed test) mismatch signal is seen for the third transition on Eiii. F: Summary heatmaps showing percent of significant transitions (p < 0.05, Wilcoxon signed-rank left-tailed test) for experiments (Fi) and simulations (Fii and Fiii). Error estimates are from bootstrap resampling over all cells with replacement 1000 times. In the simulations we assume 1mV of 500Hz lowpass filtered noise. Fii: Selectivity for zero jitter at experimental (2mM) Ca^2+^. Most of the matches to experiment are poor, especially all the 15-square cases are well below experimental selectivity. Fiii: 6ms half-Gaussian distributed jitter at physiological Ca^2+^ and 37°C. Most of the experimental estimates of selectivity are within the error bars, with the exception of 15-sq, 50Hz. The introduction of jitter substantially changes the probability of selective transitions.

Based on these predictions of mismatch selectivity, we asked if we could experimentally detect pattern transitions in our slice preparation. Rather than projecting letter-shaped pixel patterns in the optical array (which would have required high pixel coverage), we generated randomized non-overlapping sets of 5 distinct patterns of 5 squares, and 3 of 15 squares (Figure 1, Methods). We delivered the same eight repeats each of four of these patterns at repeat rates of 8, 20 and 50 Hz to span a range from theta to gamma frequencies (Figure 7 E). We used the Wilcoxon test to compare experimental EPSPs for 3 pulses immediately before and after pattern transitions. We observed that mismatch detection did indeed occur, but it was selective. In the example cell we observe a single significant mismatch response in Figure 7Eiii at 50 Hz for the transition from pattern 3 to pattern 4. Over the population of cells we found that mismatch detection was dependent on a) frequency, b) specific pattern transitions, and c) number of illuminated squares. Mismatch detection occurred most often for 15 square patterns at 50 Hz (Figure 7Fi).

We observed this mismatch selectivity despite the reduction in inputs due to ChR2 desensitization over the long 32-pulse train. To estimate this reduction, we measured the field EPSP in the CA3 during the optical stimulation, and fitted an exponential decay function to the peaks (Figure 7-figure supplement 2). We find that the largest decay was ∼30% of the initial fEPSP. The fEPSP decayed over a wide range of time-courses with median around 800ms. We estimate that the recovery time from ChR2 desensitization was around 150ms, again highly different between slices (Figure 7-figure supplement 2). Thus the mismatch detection was largely robust to ChR2 desensitization effects over the pulse train.

We then asked if our model with full stochastic synaptic kinetics could replicate these results. We ran our model over a population of 30 simulated cells in which the specific connectivity differed due to different random number seeds, and furthermore the stochastic chemistry also used different seeds (Methods). In addition we applied a range of spike jitter times (0 to 10ms, Figure 7-figure supplement 3). There was considerable variability between simulations, but overall we obtained a higher likelihood of mismatch responses than experiment at the default model settings with 6ms jitter. (Figure 7Fii). The percentage of significant mismatch responses both for 5 and 15 squares became closer to experiment when we incorporated more physiological conditions of 37℃ and 1.35 mM external Ca^2+^, and added 500Hz bandpass filtered 2mV Gaussian noise (Figure 7Fiii).

We next asked how sensitive the mismatch detection was to spike jitter, since in-vivo neural activity has a distribution of spike timings. We reran our model with 30 distinct simulated networks projecting to a CA1 neuron, each for a range of jitter values from 0 to 10ms. We performed a cell-wise bootstrap analysis to obtain error estimates for mean selectivity for each condition. We find that for 5-square stimuli selectivity is high at 0 jitter for all frequencies, but surprisingly for 15-square, selectivity is higher when jitter is nonzero (Figure 7-figure supplement 3). We suggest that when synaptic inputs are synchronous their effects add sublinearly, so some degree of jitter causes larger EPSPs when there are many inputs, as in the 15-square case. Consistent with this, the centroid for selectivity was always at a higher jitter level for 15-square than for 5-square (Figure 7-figure supplement 3, bootstrap test, p < 0.001).

For the 50-Hz stimulus, a 10ms jitter is half the stimulus interval, so the onset of patterns are quite smeared. Thus, as expected, selectivity fell both in 5 (bootstrap p<0.001) and 15-square (bootstrap p=0.005) cases for 50 Hz when jitter was larger than 4 ms. (Figure 7-figure supplement 3).

As discussed later, further network details such as cell-to-cell heterogeneity and different classes of inhibitory interneurons may be needed to account for the remaining differences between model and experiment.

Thus, at this stage we had predicted STP-dependent single-neuron detection of mismatches in sequences of inputs, confirmed it experimentally. We then showed that our model performed mismatch detection in the presence of stochastic synapses, stimulus timing jitter, recording noise, and in slice conditions as well as more physiological ones of 37℃ and 1.35 mM external Ca^2+^.

#### Mismatch detection is effective in spiking neurons

Our previous analysis of mismatch detection was in the sub-threshold regime. We next introduced spiking, as a closer approach to relevant network responses. To do this we increased the conductance of voltage-dependent sodium and potassium delayed rectifier channels in the soma of the CA1 neuronal model from placeholder values to firing values (125 nS and 141 nS) and doubled the synaptic weights of AMPAR and GABAR inputs to 10 and 20 respectively. No other changes were made to the baseline model. We obtained spiking responses for 50 stochastic runs with patterned input at 50 Hz (Figure 8 A,B), and averaged the resulting firing rates with a 10ms window, as a proxy for population activity (Figure 8 C-K). As expected, the resultant spiking was locked to the period of the simulated patterned input series.

**Figure 8.**
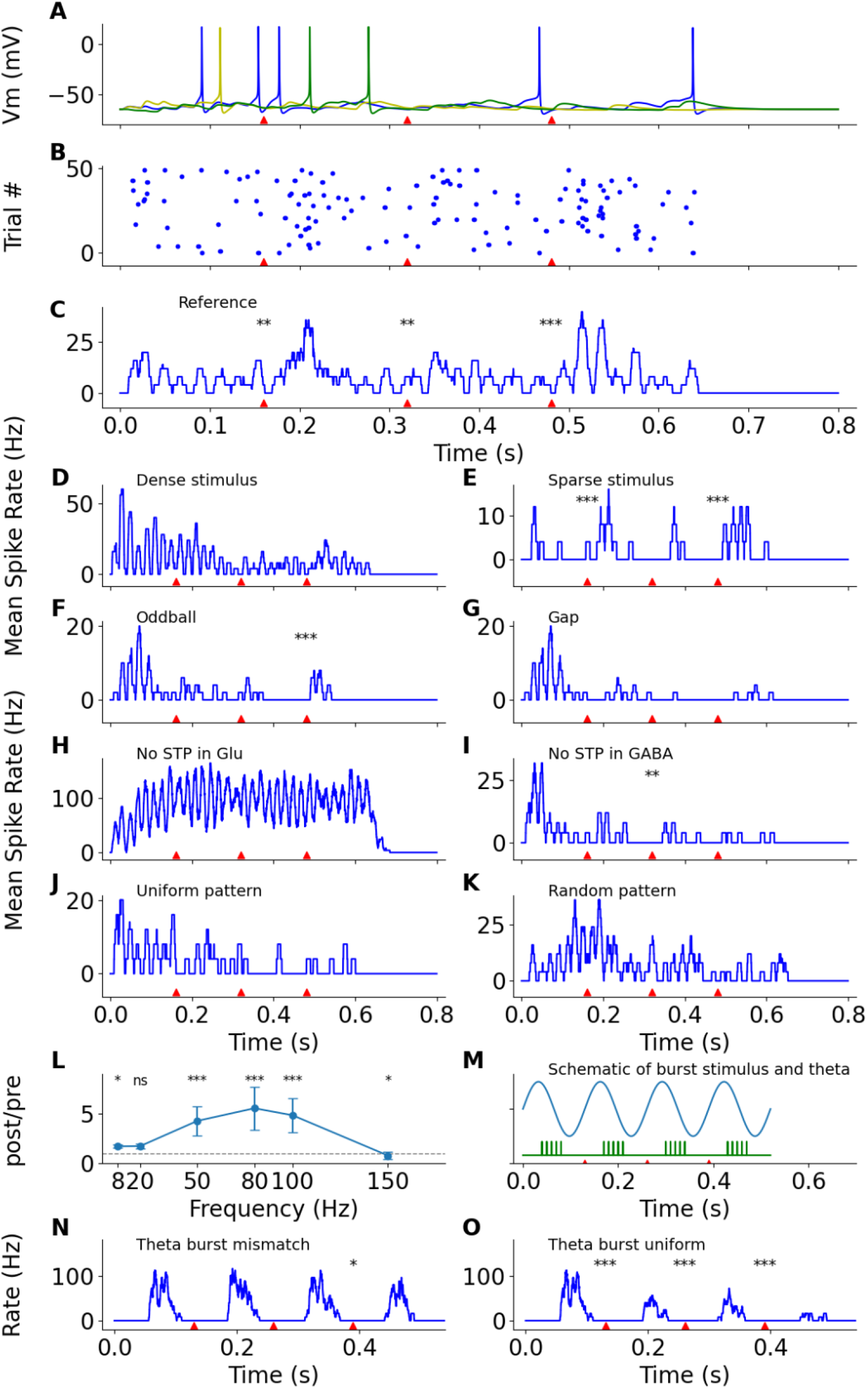
Spiking responses are selective for mismatch in patterned sequences. All runs were performed using a spiking neuron model with stochastic synaptic chemical kinetics and each run consisted of 50 trials. Red triangles indicate transition times between patterns. Asterisks over the red triangles indicate significance of transitions. For panels A to E and H to K, the stimulus consists of eight repeats of each four patterns as in AAAAAAAABBBBBBBBCCCCCCCCDDDDDDDD. Traces C-K and N-O are spike rates with a moving window of 10 ms and 5 ms respectively. In C-K, Mismatch statistics use the binomial test to compare the counts of the three pulses immediately before vs. the three immediately after a transition. A: Spiking responses on three illustrative trials. B. Raster plot of spiking for reference stimulus. C: Reference model exhibits mismatch responses for the first two pattern transitions (p=0.002, 0.032, 0.434). D: Dense stimuli (12.5% squares nulled) do not give any mismatch signals. E: Sparse patterns (94% of squares nulled) elicit a mismatch response in two transitions (p=1.5e-5, 6.1e-5). F: Oddball response. There was a strong and pattern selective mismatch response (p=9.5e-7) to the third oddball (deviant) stimulus. Oddballs were presented every 8th pulse, against a uniform background pattern: AAAAAAAABAAAAAAACAAAAAAADAAAAAAA. G: Gap stimulus. Similar to oddballs, gaps were presented every 8th pulse, but instead of a deviant stimulus, no stimulus pattern was delivered. There was no significant response, and activity declines over the course of the trial. H. Lack of STP in Glu removes mismatch detection, and spike rates are high. I. Lack of STP in GABA results in sustained strong inhibition and sparse firing, but some mismatch selectivity remains for the second transition (p=0.004). J: Control with uniform pattern (A repeated 32 times). The spiking response rapidly drops to close to zero. K. Control with randomized pattern (one of pattern A to pattern E) on each pulse. There is a slow decay over the 32 pulses. L: Mismatch response is tuned to frequency of pattern repeats. The strongest responses are in the gamma range between 50 and 100 Hz. Response compares three post-transition vs three pre-transition pulses. Y axis is ratio, p-values are Binomial test for spike counts comparing post vs pre.). O: Schematic of gamma burst stimulus (green) repeated at theta frequency of 7.69 Hz (blue). Each burst consisted of a pattern repeated 5 times at 100 Hz. Panels N and O use the Binomial test to compare spike counts in the first theta cycle with those in the next three cycles. N: Theta modulated gamma burst responses showed strong spiking responses when each burst had a different pattern. O: Spiking in second, third and fourth theta cycles was lower than the first when each burst had the same pattern (p=1.9e-07, 3.1e-07, 6.68e-21).

The reference spiking model replicated both mismatch detection and pattern selectivity, with significant elevation of spiking at each of the transitions (red triangles Figure 8C, p = 0.0036, 0.0040, and 2.1e-5 for the respective transitions, Binomial test). The spiking model supported the heuristic prediction that networks with high overlap of excitatory inputs to CA1 would be less sensitive to mismatches (Figure 8D, E). None of the transitions in Figure 8D (dense stimuli, 34% overlap) were significant, but two transitions in Figure 8E were significant (sparse stimuli with 2.5% overlap, p = 1.53e-5 and 6.1e-5).

We next explored a range of stimulus conditions for functional implications. First, we considered the frequently used behavioural paradigm of ‘oddball’ detection, which produces a rapid ∼100-250ms evoked potential in several modalities (Butler 1968; Garrido et al. 2009). We delivered a constant, repeating pattern interspersed every 8 stimuli with the oddball (deviant) stimulus. We obtained a selective response to the third oddball presentation (Figure 8 F). The repeated stimuli resulted in a strong depression of firing rates, hence for this and the next stimulus, we doubled the number of trials to 100 in order to increase the statistical strength of the binomial test.

Gap stimuli, where the deviant stimulus is an absence of stimulus, are also known to be strong behavioural triggers for oddball responses (Garrido et al. 2009). We implemented this as above, simply replacing the oddball stimuli with an absent stimulus. In our model the gap stimuli did not give any significant responses (Figure 8 G). In the discussion we consider how gap and other complex deviant stimuli may be detected without high-level network mechanisms.

We then asked how STP in E and I contribute to mismatch detection. Selectivity was completely lost when we removed STP from glutamatergic synapses (Figure 8 H). In the absence of GABAR STD, there was a steep decline of spiking activity over the time course of the trial (Kendall’s tau=-0.456, p=0.0003, Figure 8 I), consistent with the requirement for EI balance over multiple pulses in order to retain excitability. Selectivity, albeit with very few spikes, was still present for the second transition (Binomial test, p=0.004, Figure 8 I).

We next ran two controls to confirm the role of transitions in eliciting mismatch responses. In the first control we delivered the same spatial pattern for all 32 pulses (Figure 8J). As expected, this led to a rapid decline in spiking (Kendall’s tau=-0.53, p=3.6e-5). Next, we randomly selected one pattern among our set of five for each pulse (Figure 8 K). Surprisingly, this also gave a decline in spiking, though it was not as steep as the uniform pattern (Kendall’s tau=-0.314, p=0.008). Upon further analysis, it turned out that random sequences had sufficient overlap to trigger a subset of the same synapses (Figure 7-figure supplement 1). The net effect was that excitatory synapses were stimulated at ∼21 Hz for random patterns, as compared to the 50 Hz for uniform patterns. This led to slower synaptic depression.

We next asked if there was a frequency-dependence of mismatch detection in the spiking model (Methods). Briefly, we ran a frequency sweep from 8 to 150 Hz, with 10 simulated cells with distinct random seeds for connectivity. We found that there was a broad peak of selectivity in the mid-gamma range from 50 to 100 Hz (Figure 8 L).

To place the gamma-frequency tuning in a more physiological stimulus context, we considered the well-known phenomenon of gamma bursts modulated by a theta background(Lisman and Jensen 2013). We modeled theta-modulated gamma bursts, where each theta cycle had five pulses of input at 100 Hz, with the start of each burst separated by 130 ms to give a 7.69 Hz theta rhythm (Schematic in Figure 8 M). When each burst was a different pattern, their amplitude was nearly consistent (Figure 8N), but the final burst was smaller than the first (p=0.0424, Binomial test). However, when the same pattern was repeated over successive bursts there was a steep decline in burst amplitude (Figure 8O, p=1.9e-7, 3.1e-7, 6.7e-21, Binomial test compared to first burst). Thus mismatch detection also works when the ensemble of neurons comprising a gamma burst changes between theta cycles.

In summary, our final set of model predictions showed that mismatch detection remains sharp in spiking neurons. The mismatch response also occurs for oddball (deviant) patterns, but not gaps. We confirmed that STP on both E and I synapses are required, and showed that mismatch detection also works for 100 Hz gamma-frequency bursts riding on the theta rhythm. Together, these support possible functional relevance of these mechanisms for spatio-temporal pattern selectivity in in-vivo-like network conditions.

#### Parameter sensitivity analysis

To conclude the characterization of our model, we asked: How robust is mismatch detection, and how do different network parameters affect it? We varied the five key network parameters: weights for Glutamate and GABA receptors, and the connection probabilities from CA3 to CA1, CA3 to interneurons, and interneurons to CA1 (Figure 9A). We also varied the stimulus pattern density, expressed as the percentage overlap of activity converging onto Glu synapses on CA1 due to any given pattern (Figure 9A, Figure 7-figure supplement 1, Figure 9-figure supplement 1).

**Figure 9.**
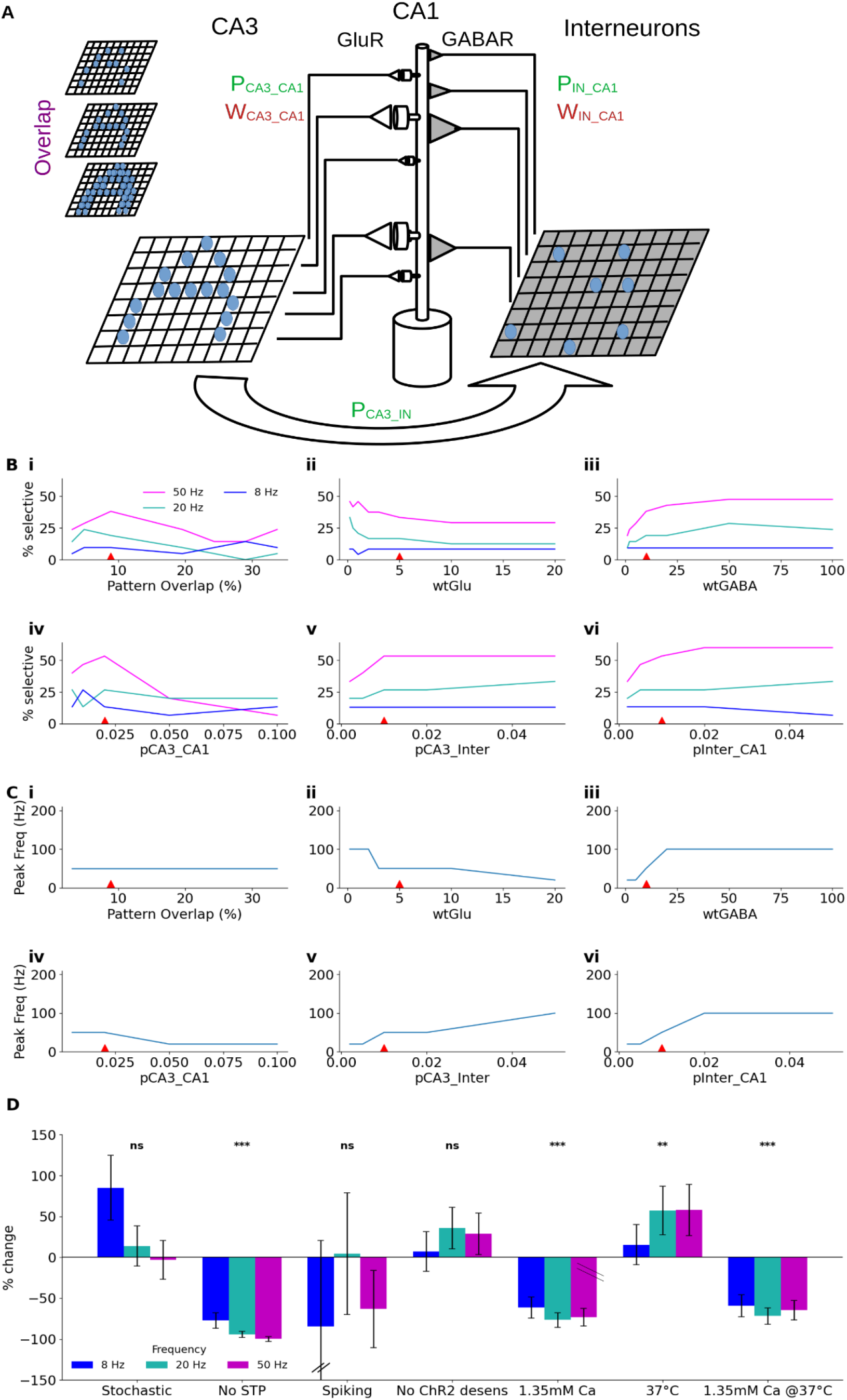
Parameter sensitivity of network model. A: Schematic of model, indicating the six network parameters. Overlap specifies the fraction of postsynaptic axons which are activated by more than one illuminated square as pattern density increases. W_Glu_ and W_GABA_ specify synaptic weights for gluR and GABAR synapses, measured as 1/ohm.m^2^. P_CA3_CA1_ is the probability that a given CA3 neuron connects to the CA1 neuron. Similarly for P_CA3_inter_ and P_Inter_CA1_. B: Pattern selectivity as a function of the six parameters. Red triangles indicate parameter values used for the reference model. Note the distinct selectivity profile for 8, 20 and 50 Hz, where 50Hz is almost always more selective over the entire range of parameters. C: Frequency of greatest mismatch detection as a function of the network parameters. The optimal mismatch detection frequency can be tuned over a wide range by network parameters, but for pattern overlap, in which case it remains at 50Hz. D: Effect of global network changes on mismatch detection, compared to baseline deterministic model. Stochasticity does not affect mismatch detection (T=1.53 p=0.13), but lack of STP completely eliminates it (T=-8.77, p=2.2e-12). Introduction of spiking greatly increases trial-to-trial variability but there is no difference of the mean compared to the reference (T=-1.16, p=0.25). The absence of ChR2 desensitization also does not change it (T=1.81, p=0.073). Lowering external Ca to 1.35mm reduces selectivity (T=-6.1, p=3.2e-8). Raising the temperature of the network to 37 degrees improves mismatch detection (T=2.99, p=0.0034). Finally, the simulated in-vivo case with 1.35mM ext Ca and 37 C has a net reduction (T=-5.53, p=3.4e-7) amounting to about 65% smaller responses than baseline. For each of the runs and each frequency, the ratio (post-pre)/(post+pre) was taken for the 3 pulses before (pre) and after (post) the transition. % change = 100*(run_mean - baseline_mean)/baseline_mean. The three frequency terms were merged and Welch’s T-Test was used to compare each run against the baseline run.

We first conducted a simple parameter sweep, measuring mismatch selectivity as a function of different values of the six parameters (Figure 9B). To estimate selectivity, we repeated the design from Figure 6 panel D in which we simulated 6 distinct instances of the network and obtained the probability of mismatch detection for any given transition. The general outcome was that different pattern repetition frequencies exhibited distinct profiles of parameter dependence, with the 50Hz pattern having the greatest selectivity for most of the range.

We observed that sparse patterns (with low overlap) had stronger mismatch responses, particularly at 20 and 50 Hz (Figure 9Bi). There was little dependence on the weight of the Glutamate receptor. Mismatch was small for low values of wtGABA, supporting the role of inhibition in mismatch detection (Figure 9Biii). Greater values of excitatory synaptic projection probability, pCA3_CA1 led to lower selectivity (Figure 9Biv). This is complementary to the trend for pattern overlap (Figure 9Bi), which is expected since they similarly affect the number of activated synapses for each stimulus pattern. Both the inhibitory projection terms pCA3_Inter and pInter_CA1 showed a modest increase in selectivity with increasing connectivity (Figure 9Bv, 9Bvi). We interpret this as due to increasing overlap between input patterns, so that depression of inhibitory inputs carries over between patterns.

Since these runs showed considerable difference between the three reference pattern repetition frequencies, we performed a systematic analysis to find the frequency at which selectivity peaked. We plotted peak frequency as a function of the same set of six network parameters (Figure 9C). There was a small increase in peak frequency for intermediate levels of pattern overlap (Figure 9Ci). In general, peak selectivity was seen in the gamma frequency range. but at low values of GABA input the peak went down to 20 Hz (Figure 9 Ciii, Cv, and Cvi). High connectivity between CA3 and CA1 also resulted in a low (8Hz) peak frequency (Figure 9 Civ). Conversely, the highest tuning frequency was obtained for very low values of GluR weights (Figure 9Cii).

We finally tested seven qualitative manipulations to the network (Figure 9D). In all cases we asked how selectivity changed compared to values obtained for a deterministic, subthreshold reference model. Reassuringly, stochasticity caused only a small decrement in selectivity. This supports the use of deterministic models as efficient proxies for the full stochastic version. Lack of STP, not surprisingly, completely abolished selectivity. The introduction of spiking led to a considerable increase in variability between runs and simulated cells, but there was no significant change in overall selectivity. Removal of ChR2 desensitization also did not change selectivity.

The remaining three manipulations explored different axes along which our slice preparation differed from physiology. At reduced (1.35 mM) Ca2+ we find lower STP both for E and I, and as expected, the mismatch detection was weaker. At elevated temperature (37℃), mismatch detection became stronger. Finally, when these were combined, mismatch detection was about 65% weaker than the default model.

In summary, mismatch detection in our model is robustly present and can be tuned over a wide range of network parameters and model assumptions, with the notable exception that it is absolutely dependent on the presence of STP.

## Discussion

We probed CA3-CA1 input-output properties for functional outcomes of pattern-specific short-term plasticity using spatially and temporally patterned optical stimuli. EI balance and summation properties tilted briefly to excitation on the second or third pulse of a burst due to differential STP profiles for excitation and inhibition. We parameterized a molecule-to-network multiscale model of CA3-CA1 network and plasticity using a series of burst, summation, and Poisson train inputs of optically defined input patterns. The model predicted that STP dynamics of E and I inputs provide a mechanism for single cells to detect transitions in input pattern sequences, which we then confirmed experimentally. We used the model to explore network configurations which could detect such transitions, and showed that spiking is strongly mismatch-tuned in continuous as well as theta-modulated gamma bursts. Overall, EI+STP based mismatch detection is robust over a wide range of network conditions.

### Relevance to in-vivo computation

Slice physiology is, by definition, a reductionist way of investigating in-vivo circuit function. Here we have taken several steps to support our claims of relevance to brain computation. 1. We use optical patterned stimuli to stimulate a cross-section of CA3 neurons with a variety of distributed patterns, theta, and other frequency rhythms. These stimuli are sparser and more dispersed than Schaffer collateral electrical stimuli which tend to stimulate adjacent fibres and in most cases are very strong. 2. Prior work in rat slices has shown that short-term plasticity properties are consistent in the 33-38 ℃ range (Klyachko and Stevens 2006b). Further, we show that the small 3.5 degree temperature increase to physiological levels strengthens, not weakens, our main finding of pattern mismatch detection. Additionally, we made a ‘physiological’ model variant with 1.35 mM external Ca2+ as well as 37℃ synapse kinetics, and show that it too exhibits the key computation of mismatch selectivity (Figure 7, figure 9). 3. Our model is a network + cell + synaptic signaling model, thus spanning multiple scales of function relevant to physiology. We factor in CA3 to CA1 connectivity distributions in addition to cell physiology and presynaptic signalling dynamics. We have, of course, simplified the network, most notably in the use of only one inhibitory interneuron class which maps to parvalbumin-positive fast-spiking interneurons with perisomatic connectivity. This level of detail was chosen as it was able to quantitatively fit a large number of observations with minimal circuit complexity. 4. We have extensively matched experiment to simulation at each of the specified levels: synaptic (Figure 4), cell (Figure 5, Table 1), and network (Figure 6, Table 1), in which we explicitly used Poisson input patterns and synaptic stochasticity to stress-test the model across scales. In almost all cases the simulation values fall within the range of measured cellular properties. 5. The prediction of mismatch detection is highly robust to network parameters (Figure 9), stimulus frequency (Figure 9), and stimulus timing jitter (Figure 7), and we confirmed it in slice experiments (Figure 7). This strengthens the case for it working in vivo.

As expected for a reduced model, our predictions differed quantitatively from experiments in a few respects. The strongest of these is the difference in mismatch selectivity at 50 Hz in Figure 7Fi vs 7Fiii. We speculate that this is due to one of the major simplifications in our model, the use of a single class of inhibitory interneurons. An additional point of difference is the decay time-course of EPSP vs ISI from Figure 6F. Here again we implicate the simplification with respect to interneurons. An additional likely cause is our limited use of the diversity of STP profiles in the model compared to experiment. Specifically, we used STP profiles from Figure 4-figure supplement 1, but the population is quite diverse (Figure 4-figure supplements 1 to 4). Finally, despite our efforts to model the CA3 cell population with ChR2 desensitization constrained by experiment, we have clearly simplified the recurrence and optical illumination profiles of the CA3. Thus, at the spiking level, our stimulus patterns can only approximate physiological activity.

In-vivo input activity to this circuit is complex, and on successive stimuli there will be a distribution of exact CA3 pyramidal cell AP timings, number of APs per cell, and number of activated axons. To the extent that model robustness (Figure 9, Figure 7-figure supplement 3) gives insights into the real system we suggest that our small model of network + single neuron model + presynaptic signaling chemistry captures some of the attributes of the biological system, which lead to such robustness. This robustness is almost a necessity in a system where the probability of release for individual synapses is in the range of 20% (Murthy et al. 1997): any signal transmission or plasticity-type computation has to accommodate huge stochastic uncertainty on any given trial.

### A resource to model the CA3-CA1 feedforward circuit

A major outcome of our study is the development and parameterisation of a multiscale, chemical+electrical+circuit level model of the CA3-CA1 feedforward circuit, which runs rapidly even on laptop hardware (∼20x slower than real time on an AMD 6800HS). Feature-wise, several studies combine signalling (typically postsynaptic spine and dendrite signalling) with electrical signalling (Bhalla 2017; Dainauskas et al. 2023; Dorman and Blackwell 2022; Mattioni and Le Novère 2013; Somashekar and Bhalla 2024; Bhalla 2011). Some of these combine compartmentalized signalling with standard branched neuronal electrophysiology calculations, as we do. Others include detailed 3-dimensional reaction-diffusion kinetics on spines(W. Chen et al. 2022) or boutons(Nadkarni et al. 2012, 2010) and these run much more slowly. Our current model is distinct in that it is truly multiscale, closely constrained by experiment, yet runs on modest hardware. It incorporates the network, a conductance based model of a CA1 pyramidal neuron, and chemical kinetic models of a population of stochastic synapses on its dendrite.

Our network model is much reduced compared to models with exhaustive cellular and network-level detail(Markram et al. 2015). Its simplicity enables extensive exploration of the network parameters and comparison with recorded activity under a series of well-controlled stimulus patterns (Figures 4-9). For almost all tests, including molecular, single-cell, and network, the model was within the range of experimental readouts, but it may be necessary to extend the model with more interneuron subclasses to match experiment with respect to mismatch detection probability. (Figure 7 E).

Our model is modular. One can swap out different cellular geometries, provide distinct synaptic signaling models using SBML or compatible standards, and alter the detail of the network components. In the current study, we employed four different spine kinetic models, with different combinations of STP in E and I, simply by loading in different kinetic definitions. Thus the model is consistent with the ‘Interoperable’ and ‘Reusable’ aspects of FAIR principles (Eriksson et al. 2022).

### EI balance is dynamically tuned through differential STP of E and I inputs

EI balance is a ubiquitous mechanism for controlling neural excitability. Numerous processes, including inhibitory plasticity, homeostatic plasticity, and developmental co-tuning have been proposed to underlie the origin and maintenance of EI balance (Agnes et al. 2020; Agnes and Vogels 2024; L. Chen et al. 2022; Jia et al. 2022; Lagzi and Fairhall 2024; Mackwood et al. 2021; Miehl and Gjorgjieva 2022; Trapp et al. 2018; Wu et al. 2022). EI balance is dynamic, due in part to differential effects of STP on excitatory and inhibitory inputs to the postsynaptic neuron (Bartley and Dobrunz 2015; Sun et al. 2018; Bartley et al. 2015; Wu and Zenke 2021; Klyachko and Stevens 2006a). However, the mode of delivery of these stimuli may lead to narrow timing and spatial patterning of activation of the inputs to CA1. For example, studies using field electrode stimulation of the Shaffer collaterals report a sustained shift to excitation during burst input (Klyachko and Stevens 2006a). In contrast, our sparse optical patterned stimuli results in a small window of escape from EI balance around pulse 2 or 3 in a burst (Figure 3), following which both E and I undergo depression to restore balance (Figure 3, 8). Thus, spatial patterning intersects with short-term plasticity to add another layer of timing control through gating of E-I balance (Rotman et al. 2011).

### STP leads to single-neuron mismatch detection in pattern sequences

Where can temporally precise gating be useful? At the single-synapse level, synaptic depression performs the operation of decorrelation, leading to more efficient coding (Goldman et al. 1999; Rotman et al. 2011). Our observations can be framed as decorrelation of the spatial pattern (alternatively, activity vector) of multi-synaptic input converging onto a neuron(Kremkow et al. 2010). A putative example of this is place field formation in novel environments. Dombeck and co-workers (M. E. Sheffield and Dombeck 2019; M. E. J. Sheffield et al. 2017), found that dendritic inhibition is reduced when a mouse traverses a new environment. Consistent with this, our simulations predict elevated spiking when novel patterns occur, in part due to presynaptic depression of inhibition (Figure 7,8).

Mismatch detection has been extensively studied at the whole-organism level, particularly in the context of a large EEG transient following an unexpected stimulus in a regular stream of events. This is known as mismatch negativity. There is a vigorous debate on the mechanisms of auditory mismatch negativity (e.g., (Garrido et al. 2009; May and Tiitinen 2010; Näätänen et al. 2005)) of which the fresh-afferent model maps to part of our proposed mechanism (Figure 10A). Leaving aside the obvious differences between auditory cortex and hippocampus, we frame our model as a transient differential tilt in EI balance (Figure 3, Figure 8A,B, Figure 10B), in distinction to the fresh-afferent model. This makes our model robust over a wide range of stimulus and network conditions (Figure 9), and has the functional implication that transient responses remain at about the same amplitude over a prolonged stimulus sequence (Figure 8B, Figure 10B), rather than declining. We examined only three of several possible stimulus changes for mismatch detection: Change in pattern, a single oddball stimulus, and a gap. Of these, our model detected the first two. We speculate that two circuit elaborations may extend the applicability of our model to more categories of mismatch. First, feedback inhibition in CA1 through other interneuron classes might introduce a time-delay element capable of resolving gap stimuli. Second, upstream areas may encode higher order stimulus features such as gaps, duration, intensity, localization, and frequency steps into distinct input patterns. Our proposed EI-balance shift mechanism could be a common end-point for all of these. This would transform quite complex mismatch detection tasks into a uniform computation of pattern change, generalizing the mechanism to stimuli which were previously considered to require a more complex network-level implementation (Garrido et al. 2009; May and Tiitinen 2010; Näätänen et al. 2005).

**Figure 10:**
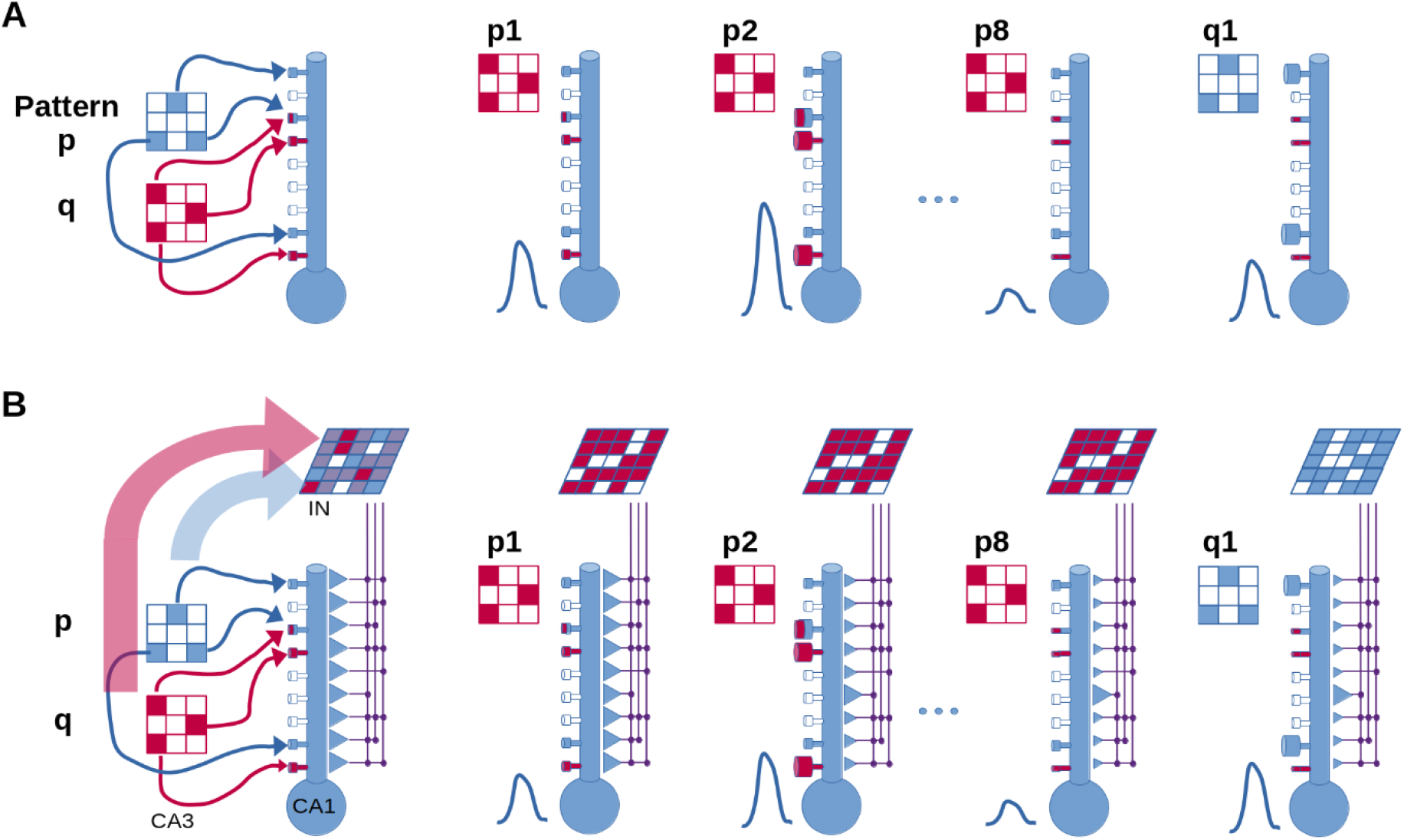
Proposed mechanisms for mismatch detection. A: Fresh-afferent model. Left: Circuit involving only excitatory synapses from CA3 to CA1. The two input patterns p and q (in the CA3) have only a little overlap. Right: on successive stimulus p repeats, the STP in the synapses (indicated by spine size) leads first to an elevated response, and then to depression. When the stimulus changes to pattern q, most of the afferents are fresh so we get an elevated response. B: EI-balance shift model. The excitatory component of the circuit is as in A, but in addition there is an intermediate interneuron layer receiving input from the CA3. During a train of pulses of pattern p, E transiently exceeds I, then both undergo STD. When a different pattern q arrives, there is again a transient advantage for the E input because the two-stage connectivity leads to substantially overlapping input on the I inputs, so most of them are still depressed. Overall, the transient response at p2 and q1 are closer in amplitude, or equivalently, the pattern selectivity is less dependent on the averaged activity history.

We note that the time-scale of our mismatch response (∼125 ms) is similar to the 100 to 200 ms of the classic auditory mismatch negativity (Butler 1968). A similar computation of stimulus-specific adaptation in the auditory cortex has been modeled using a recurrent network with short-term depression(Yarden and Nelken 2017). Our findings of pattern mismatch detection may also provide a mechanistic interpretation of published results showing behavioural and physiological selective responses to changes in visually delivered patterns(M. E. Sheffield and Dombeck 2019). The authors found that a rate model with short-term depression was able to replicate the experiments. In a recent study, signatures of prediction error (implemented by absence of conditioned response) were observed in recurrent cortical networks in organotypic preparation(Liu and Buonomano 2024). These prediction errors were visible after a 24-hour long training period that achieved the associative conditioning. Thus the study showed the role of inhibition stabilized attractor networks of cortical circuits to produce prediction-error like responses in single neurons. Our model achieves equivalent computations with simple feedforward connectivity at a single-neuron level, without training, and with direct linkages to the underlying physiological mechanisms.

Both our experiments and models show selectivity for specific transitions. This suggests a further computation of sequence disambiguation. Consider sequence S1 AAAABBBBCCCC and sequence S2 FFFFBBBBGGGG, overlapping in ensemble B. We stipulate the presence of neuron S1_AB_ which detects the A->B transition, and so on. Then S1 is uniquely identified by the activity of S1_AB_,S1_BC_, and S2 by S2_FB_,S2_BG_. This is a rapid and parsimonious mechanism for distinguishing two intersecting sequences such as time-cell sequences (Modi et al. 2014), on sub-second time-scales.

### Mismatch detection is pronounced in the gamma band

Gamma oscillations have been implicated in memory formation and communication between brain regions (Van Vugt et al. 2010; Misselhorn et al. 2019). Gamma bursts are typically modulated by the theta rhythm (Lisman and Jensen 2013), which is itself closely coupled to sensory intake such as sniffing (H Eichenbaum et al. 1987; Tsanov et al. 2014) and whisking (Grion et al. 2016), and also to movement (Joshi et al. 2023). Our proposed mechanism for mismatch detection is enhanced precisely in the gamma range (Figure 9). Further, we find that theta-modulated gamma bursts perform mismatch detection between theta cycles (Figure 8 N,O). We propose that sensory snapshots obtained through sniffs or whisks provide the CA1 with theta-modulated gamma bursts of stimulus-specific activity vectors. When inputs change from one theta cycle to the next, individual cells in CA1 light up to detect specific changes between stimuli, such as odor 1 changing to odor 2, or texture 1 to texture 2 (Figure 7, 8). This implements a dimensionality reduction of a sequence of sensory inputs into a trajectory through a space of stimulus transitions.

## Methods

### Animals

Experiments were conducted on a mouse line with the expression of cre-recombinase specific to hippocampal CA3 (Grik4Cre) obtained from Jackson laboratory (ID: 006474). The animals were housed in a cage with up to 6 cage mates and under a 14/10 light/dark cycle. Food and water were provided ad libitum. The housing and health were managed as per the National Centre for Biological Sciences Institflexutional Animal Ethics Committee (IAEC) guidelines (Project: NCBS-IAE-2022/8 (R2ME). Only transgenic F1 mice, identified with genotyping, of both sexes were used for experiments.

### Virus Injection

Transgenic mice between the ages 27-35 days were selected for virus injections. These animals were transferred to a separate holding facility where they were kept till the terminal experiment.

Virus: We used an adeno-associated virus AAV-FLEX-rev-ChR2-tdtomato (Addgene 18917) to express channelrhodopsin2 in CA3 pyramidal cells in a cre-lox-dependent manner. The virus was diluted using sterilised PBS to a titre of 1-5 x 10^12^ GC/ml and stored at -80°C in aliquots. The animals were injected with a dose of 300-500 nl of virus suspension in 10-20ms long pulses of pressurised nitrogen using a Picospritzer (Parker-Hannifin, USA). The animals were then allowed to recover under supervision and veterinary care. Terminal experiments were performed after an incubation period of 35 days or more, resulting in cre-lox-dependent expression of channelrhodopsin2 (ChR2) specific to the CA3 region of the hippocampus on the injected side. The virus AAV-FLEX-rev-ChR2-tdtomato was a gift from Scott Sternson (Addgene plasmid #18917; http://n2t.net/addgene:18917 ; RRID: Addgene_18917)(Atasoy et al. 2008).

### Electrophysiology

Slice preparation: Animals aged between 56-120 days and at least 28 days after the virus injection were used for in vitro experiments. The animals were humanely euthanised using isoflurane (Forane) and brain was harvested into ice-cold slush of cutting solution (87mM NaCl, 2.5mM KCl, 7mM MgCl_2_.6H_2_O, 0.5mM CaCl_2_.2H_2_O, 25mM NaHCO_3_, 1.25mM NaH_2_PO_4_, 75mM Sucrose) maintained over ice and aerated with carbogen (95% O_2_ + 5% CO_2_). After a rest period of 2 minutes, the hippocampus from the injected side was dissected out, and 350 µm thick sections were made using a vibrating microtome (Leica VT1200) submerged in the above-mentioned solution. The hippocampal sections were collected in a beaker filled with the recording solution (124mM NaCl, 2.7mM KCl, 1.3mM MgCl2.6H2O, 2mM CaCl2.2H2O, 26mM NaHCO3, 1.25mM NaH2PO4, 10mM D-(+)-glucose, pH 7.3-7.4 and osmolarity of 305-315 mOsm) at room temperature and allowed to recover for one hour in the dark before starting the experiments.

Recording Setup: Electrophysiological recording and optical stimulation was done on an upright DIC Olympus 53WI microscope. A slice hold-down (Warner Instruments WI 64-0246) was used to keep the slice from moving. The recording bath was perfused with the recording solution at a rate of 2 ml/min with a peristaltic pump. The recording solution was kept aerated with carbogen and the influx of the solution into the recording chamber was heated using an inline heater (Warner Instruments TC-324B) to 32-33°C.

Electrical activity was recorded using a headstage amplifier controlled by a patch clamp amplifier (Multiclamp 700B, Axon Instruments, Molecular Devices) and a data acquisition system (Digidata 1550B, Axon Instruments, Molecular Devices) using Multiclamp commander (Molecular Devices) and custom written protocols in Clampex (Molecular Devices). The electrophysiology data was acquired at 20kHz and filtered in the 0-10kHz band.

In some experiments, another recording electrode was used to record extracellular responses in the CA3 cell layer. Whole cell patch clamp recordings from CA3 and CA1 cells were made using borosilicate glass micropipettes pulled using tungsten filament puller (P1000, Sutter Instruments) and filled with an internal solution containing 130mM K-gluconate, 5mM NaCl, 10mM HEPES, 1mM Na_4_-EGTA, 2mM MgCl_2_, 2mM Mg-ATP, 0.5mM Na-GTP and 10mM Phosphocreatine, pH ∼7.3, and osmolarity ∼290 mOsm. For voltage-clamp recordings, K-gluconate was replaced with Cs-Gluconate. Field response was often biphasic with multiple cycles, hence, a peak-to-peak value was used.

The cells were either recorded in the current clamp or the voltage clamp. In current clamp recordings, the cells were maintained at a membrane potential of -70 mV. Cells were rejected if the membrane potential changed by 5mV over the course of the recording, holding current either was above 100 pA or changed by more than 25 pA to maintain the mentioned membrane potential. In voltage-clamp recordings, the cells were rejected if the series resistance crossed 25 MΩ or changed by 30% of the initial value. Liquid junction potential was calculated to be 16.926 mV according to the stationary Nernst–Planck equation (Marino et al. 2014) using LJPcalc (RRID: SCR_025044) and was not compensated.

In optogenetic preparations, the additional stimulation that comes in due to the excitation of axons and dendrites can not be ruled out. To assess that, we patched a CA3 cell and recorded its optical depolarization caused by a single spot of light (illuminated square) in a grid using a whole-cell voltage clamp. The CA3 cells have a large receptive field that arises due to the expression of ChR2 on their dendritic arbour. This receptive field is stronger closer to the soma but spans on both sides of the dendritic tree (Figure 1-figure supplement 1A). A patched CA3 cell fired action potential with 100% success (Figure 1-figure supplement 1B) when the cell layer was stimulated with patterns with multiple squares. A single square, on the other hand, was only sufficient to induce action potentials when directly incident on soma (Figure 1-figure supplement 1B).

### Optical Stimulation

Pyramidal cells in the CA3 cell layer expressing channelrhodopsin were stimulated using a DMD-based patterned projector system (Polygon 400G, Mightex Systems, Canada) using a square size of approximately 14µm x 7.8µm under a 40x water immersion objective. The spots (or squares) made a hexagonal grid with an inter-square spacing equal to 32 µm. The grid occupied the middle 2/3rd of the projector frame (336µm x 187.2µm), and the microscope was translated on a stage to make the grid part overlay on the CA3 cell layer under a 40x objective. In 1-square stimulation experiments, the CA3 cell layer was illuminated one square at a time with an interstimulus time of 3 seconds to avoid desensitisation of channelrhodopsin. In other experimental protocols, 5-square and 15-square patterns were used at various stimulation frequencies. There were five different non-overlapping patterns of 5-squares, and three non-overlapping patterns of 15-squares. Each presentation of a pattern was either for 2 ms (Poisson spike train experiments, Figure 6 and frequency sweeps, Figure 1-3) or 5 ms (Mismatch experiments, Figure 7 and frequency sweeps, Figure 1-3). We chose short light pulses for two reasons. First, to decrease jitter in the light-triggered CA3 neuron spike timings. Second, if one is below saturating light intensities, short pulses at higher power give more consistent spiking over multi-pulse trains ((Mattis et al. 2012), Figure 2A, using a 2ms pulse width). Note that the Mattis study was done at 20-23℃ as compared to our ∼33℃, and the authors state that ChR2 kinetics are strongly temperature dependent. Our light source had a total estimated maximum intensity of 6.7 mW, and for the grid size used, each square had an absolute power of approximately 14.47 µW and a power density of 100 mW/mm^2^. For all experiments, unless mentioned, the intensity of light was kept at 100%. Optical stimulation patterns were made and loaded onto Polygon using Polyscan2 (Mightex, Canada) and triggered using time series generated from custom protocols written in Clampex (Molecular Devices) and communicated to Polygon using Digidata 1440 DAC. A phototransistor (OPT101) was placed behind the dichroic to acquire the light stimulus waveform using the DAq and recorded along with the patch-clamp time series data.

### Code and Model accessibility

All scripts and models for performing simulations for figures are presented in the GitHub site https://github.com/BhallaLab/STP_EI_paper_figs. Simulations were run using MOOSE 4.0 compiled using gcc version 11.4.0 and Python version 3.10. Short calculations were run on Ubuntu 22.04 on an 8-core laptop, and large runs were done on a 128-core server also running Ubuntu 22.04. All software used is open-source and hosted on github. MOOSE is available at https://moose.ncbs.res.in and https://github.com/BhallaLab/moose. Optimization of presynaptic models was performed using HOSS 1.0(Nisha Ann Viswan et al. 2024) (https://github.com/BhallaLab/HOSS) and FindSim(Nisha A. Viswan et al. 2018) (https://github.com/BhallaLab/FindSim).

### Presynaptic Plasticity Model

Voltage clamp data from the pulse-train experiments (Figure 3) was used to train a model of presynaptic short-term plasticity (Figure 4A, Figure 4-figure supplements 1, 2, 3) implemented in MOOSE (Ray and Bhalla 2008). The model included presynaptic signalling, activation of postsynaptic glutamate and GABA receptors, and electrophysiology of a ball-and-stick model of the postsynaptic neuron held in the voltage-clamp. The glutamate receptors were placed on dendritic spines implemented as separate electrical compartments, and the GABA receptors were on the dendritic shaft. Model fitting was performed in two stages. First, the raw voltage-clamp experimental data files in hdf5 format were analysed using a Python script and the peaks and valleys for each trial were extracted and stored in an intermediate hdf5 file. Second, the HOSS pipeline (Hierarchical Optimization of Systems Simulations) (Nisha Ann Viswan et al. 2024) was used to fit the presynaptic signalling model to this data, uniquely for each cell and separately for excitatory and inhibitory synapses. There were ten to eighteen ‘experiments’ run on the model to perform the fitting for each cell: Two experiments for each frequency, two to four pulse frequencies (depending on how many were feasible in each patch recording), and an additional experiment for time-course of neurotransmitter release. All this was done for 5-square and 15-square stimulation, respectively. For each frequency (typically 20, 30, 40, and 50 Hz pulse rate), we performed peak-to-valley comparisons with experiments for eight pulses in a trial and paired-pulse response experiments focusing on the first couple of pulses. The simulations for the experiments were performed using MOOSE to compute the somatic voltage-clamp response to synaptic input on the dendrite of a ball-and-stick cellular model.

The synaptic input was obtained using mass-action simulations of presynaptic signalling for the model in Figure 4A. This, too, was part of the multiscale MOOSE model. Excitatory synapses were implemented as boutons connecting to a dendritic spine with a compartment for the spine shaft and spine head, respectively. The simulated neurotransmitter release triggered the opening of ligand-gated receptor channels for AMPA and NMDA receptors placed on the spine head. Inhibitory synapses were implemented as presynaptic boutons with the same chemical topology but different kinetics (Figure 5A) and coupled now to GABA receptors placed on the dendritic shaft. This entire process was repeated for excitatory and inhibitory synapses, respectively. All calculations for the optimisation were run deterministically since, otherwise, massive averaging would be needed for each estimate of voltage-clamp current. Equations, kinetic parameters, and concentrations for the STP and no-STP models are in Supplementary Tables 1 and 2.

#### Model tuning for physiological conditions

We implemented two sets of changes to build a distinct presynaptic model which more closely matched physiological conditions. First, we increased the rates of all the reaction steps in the E and I presynaptic model by 40%, with the assumption of a 3.5℃ rise in temperature and a doubling of rates every 10℃ (Q10 of 2). Second, we lowered the calcium influx during stimulus by a factor of extCa/defaultCa = 1.35mM/2.0mM, again both for E and I. This effectively halves synaptic release (Figure 4-figure supplement 8, compare panels A and B), which we compensated for by a simple doubling of synaptic weight, both for E and I. It also causes an increase in STP levels compared to the reference 2.0 mM extCa model. We confirmed that these changes were qualitatively similar to those expected from experiments over a range of extCa levels (Kravchenko et al. 2006; Blatow et al. 2003). However, several features in these earlier experiments differed from ours: the inter-pulse intervals, the preparation, and the temperature. Accordingly, rather than paste in their numbers of ∼20% increase in STP in E and I, we used the 37℃ and 1.35mM extCa variant of our model for additional simulations.

### Postsynaptic neuron model

The postsynaptic neuron model (Figure 4B) was a ball-and-stick model with a soma compartment of 10 μm length and 10 μm diameter and a single dendritic compartment of 200 μm length and 2 μm diameter. The dendritic compartment was ornamented with 100 evenly spaced spines, each containing a spine shaft and a spine head compartment. The spine head had a model of the AMPA receptor, the NMDA receptor, and calcium dynamics resulting from influx through the NMDA receptor. However, calcium dynamics were not utilised for any further calculations. A separate presynaptic bouton with the glutamate release kinetic model was apposed to each spine head.

The dendritic compartment also had 200 evenly spaced inhibitory synapses, likewise apposed to presynaptic boutons with the GABA release kinetic model.

Under the passive conditions of our model (RM =1.0 Ω.m^2^, RA = 1.0 Ω.m) the length constant of this model was ∼700 μm, which is much longer than the geometrical length. Hence space clamp problems are unlikely. Furthermore, the short electrotonic length means it does not matter if the Glu and GABA receptors are interspersed as opposed to GABA being more proximal.

The soma had voltage-gated Na and delayed rectifier K_DR channels in the Hodkin-Huxley formalism with kinetics drawn from previous models(Traub et al. 1991). For most models, the Na and K_DR conductances were held to small values (6 and 3.5 Siemens/m^2, respectively) as we were interested in linear somatic integration, but for spiking simulations, we increased Na and K to 400 and 450 Siemens/m^2. Equations and kinetics for the voltage-gated and ligand-gated channels in the model are presented in Supplementary Table 3.

### Network model

We implemented a simple two-layer network model converging onto a single CA1 neuron. As input, the network had a 16x16 array of integrate-and-fire neurons for the CA3. There was another 16x16 array of inhibitory interneurons implemented as sum-threshold units (Figure 5C). Excitatory projections from CA3 to the 100 synapses on the CA1 neuron were implemented as a 256x100 matrix. Default entries for the connections were zeros, and ones were filled in at random with a probability pCA3_CA1 as indicated in the table. Similarly, projections from CA3 to interneurons were defined as a 256x256 matrix with connection probability pCA3_Inter. Projections from the interneurons to the 200 GABA synapses on the CA1 neuron were a 256x200 matrix with default connection probability pInter_CA1. All told, there were just seven parameters in this model (Table 2), of which only six were varied.

Each optically stimulated spatial pattern of activity in CA3 was converted into a vector of ones and zeros by thresholding the CA3 membrane potential, calculated as per the next section. Activity on the vector of glutamatergic synapses was:

Glu synapse vector:

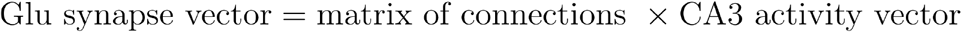

Synaptic activity was thresholded at 1 to drive the presynaptic signalling. Similarly, Interneuron vector :

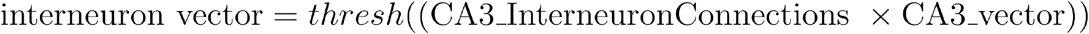

GABA synapse vector:

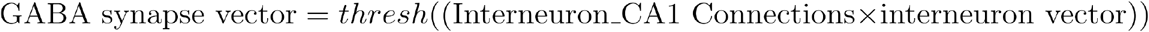

#### Stimulus model

To deliver the stimulus, we defined a set of eight input patterns. These were hand-coded as 8x8 grids of ones and zeros, set up to resemble the letters A through H. We expanded the 8x8 patterns into 16x16 patterns by replicating the pattern twice over the array (Figure 4-figure supplement 6). For the first five patterns we zeroed out a specified number of entries on the 16x16 pattern using the parameter ‘zeroIndices’ to adjust sparseness. This zeroing out was done irrespective of the original state of the entry. For the default level of sparseness (75%) the first five patterns (A through E) had 18 to 26 nonzero squares. The last three patterns (F through H) were dense, with 144, 152 and 152 squares.

The product term:

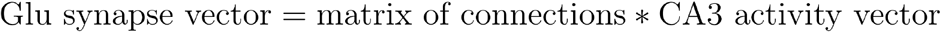

sometimes led to very different numbers of active glutamate synapses, differing by as much as 50% between patterns. Hence we carried out a preliminary calculation to fill in entries in the connection matrix to bring the final synaptic activation for all the 5-square patterns to roughly the same level (within 15%). For the default 75% sparseness this came to between 35 and 38 active glutamatergic synapses (Figure 7-figure supplement 1).

During the simulation runs we provided a time-series of patterns according to frozen Poisson timing series with a mean of 20 Hz (Figure 6) or a time-series of four patterns each repeated 8 times, at 8, 20 and 50 Hz (Figure 7). The time-series was generated with a resolution of 0.5 ms, and the stimulus duration (activation of Ca influx into the presynaptic boutons) was 2 ms. We used a fixed delay of 5 ms between the Glu and GABA synapse activation.

We implemented ChR2 desensitisation and membrane potential buildup over multiple pulses on the CA3 neurons using the following equations:

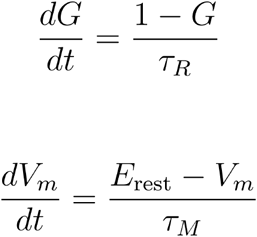

For each light pulse, we updated G and Vm as follows:

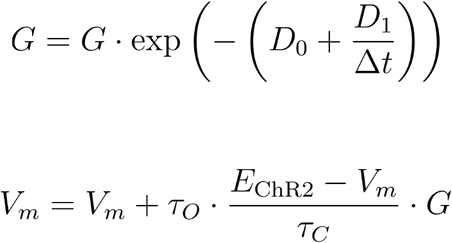

Where

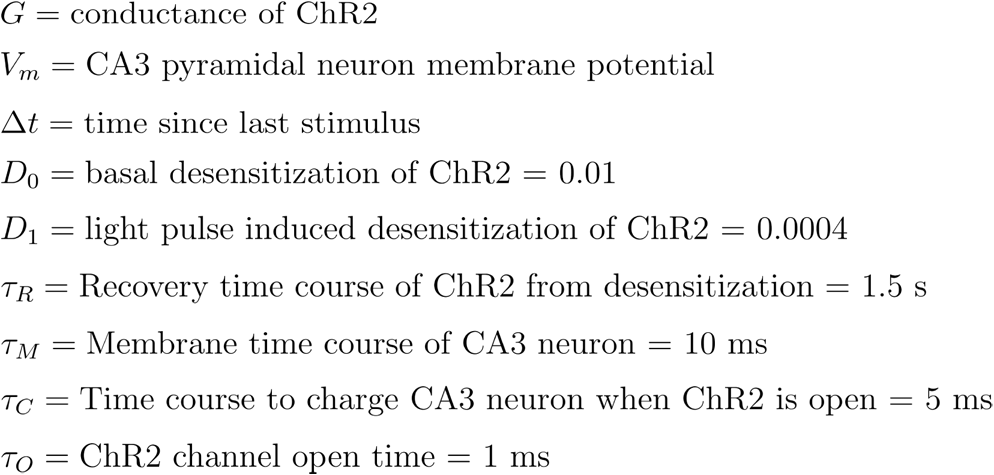

Example pulse trains with these parameters are illustrated in Figure 4-figure supplement 7.

### Simulation parallelization and Visualization

Presynaptic kinetic model fitting (Figure 4) used E/IPSC peak and valley data from experiments as a reference, and used the hierarchical optimization method HOSS (Nisha Ann Viswan et al. 2024) to adjust parameters for the Glu and GABA presynaptic release models respectively. Data generation for multi-trial stochastic runs (Figures 6, 7, 8, and 9) involved parallel calculations. The stochastic runs, in particular, were embarrassingly parallel in that they required multiple repeats of the same model with different random seeds. This was orchestrated through Python scripts using the multiprocessing library. The calculations were carried out on a 128-core AMD Epyc 7763 server. Data from these runs was organised into pandas data frames and dumped into hdf5 files in the same format as the experimental data. Figure 5 involved deterministic calculations and ran in serial to generate its hdf5 files. Figures 5 to 9 were generated from these data files using pandas, Matplotlib and Python scripts.

3-D model visualisation (Figure 4 and Supplementary movie) was performed using the built-in interface within MOOSE, to the 3-D graphics library *vpython*.

## Supporting information

Supplementary Tables for Parameters.

## Author Contribution

Aditya Asopa: Conceptualization, Data curation, Analysis, Investigation, Visualization, Methodology, Writing original draft, Writing review and editing;

Upinder Singh Bhalla: Conceptualization, Resources, Simulation, Visualization, Analysis, Software, Supervision, Funding acquisition, Writing original draft and revision, Project administration, writing review and editing

## Supplementary Figures

**Figure 1-figure supplement 1.**
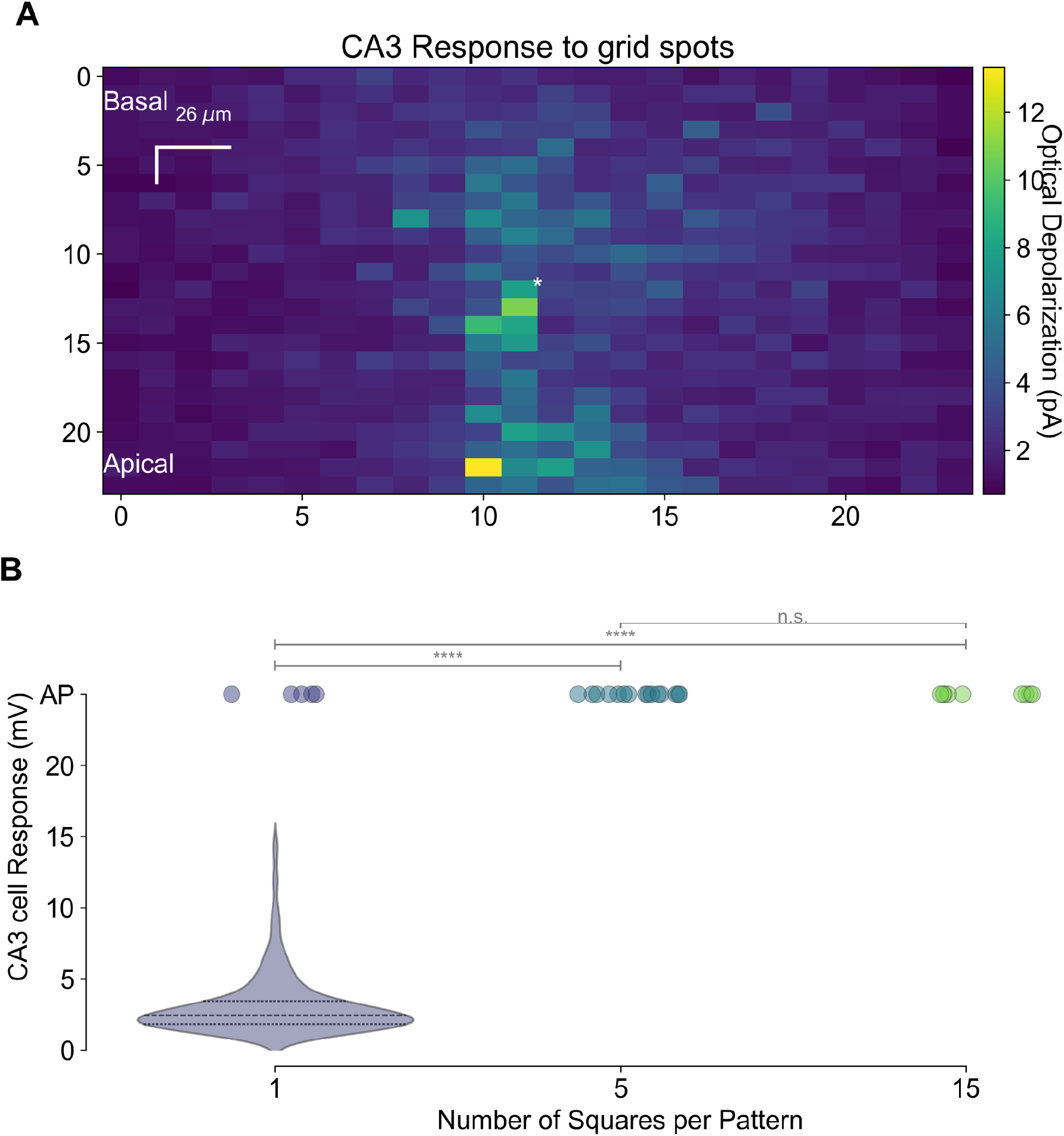
A) Heatmap of optogenetic depolarisation of a single CA3 pyramidal cell obtained by single squares of light in a 24x24 grid while keeping the patched PC in the centre of the frame (asterisk). B) Distribution of optical depolarization in CA3 PC against patterns of size 1 square (single illumination spots), 5 squares, or 15 squares. Filled circles denote the instances when the patched cell fired an action potential. For patterns of size 5 and 15, all the trials resulted in spikes, hence there is no subthreshold distribution of EPSPs. For 1-square patterns, four out of 576 squares induced a spike.

**Figure 1-figure supplement 2:**
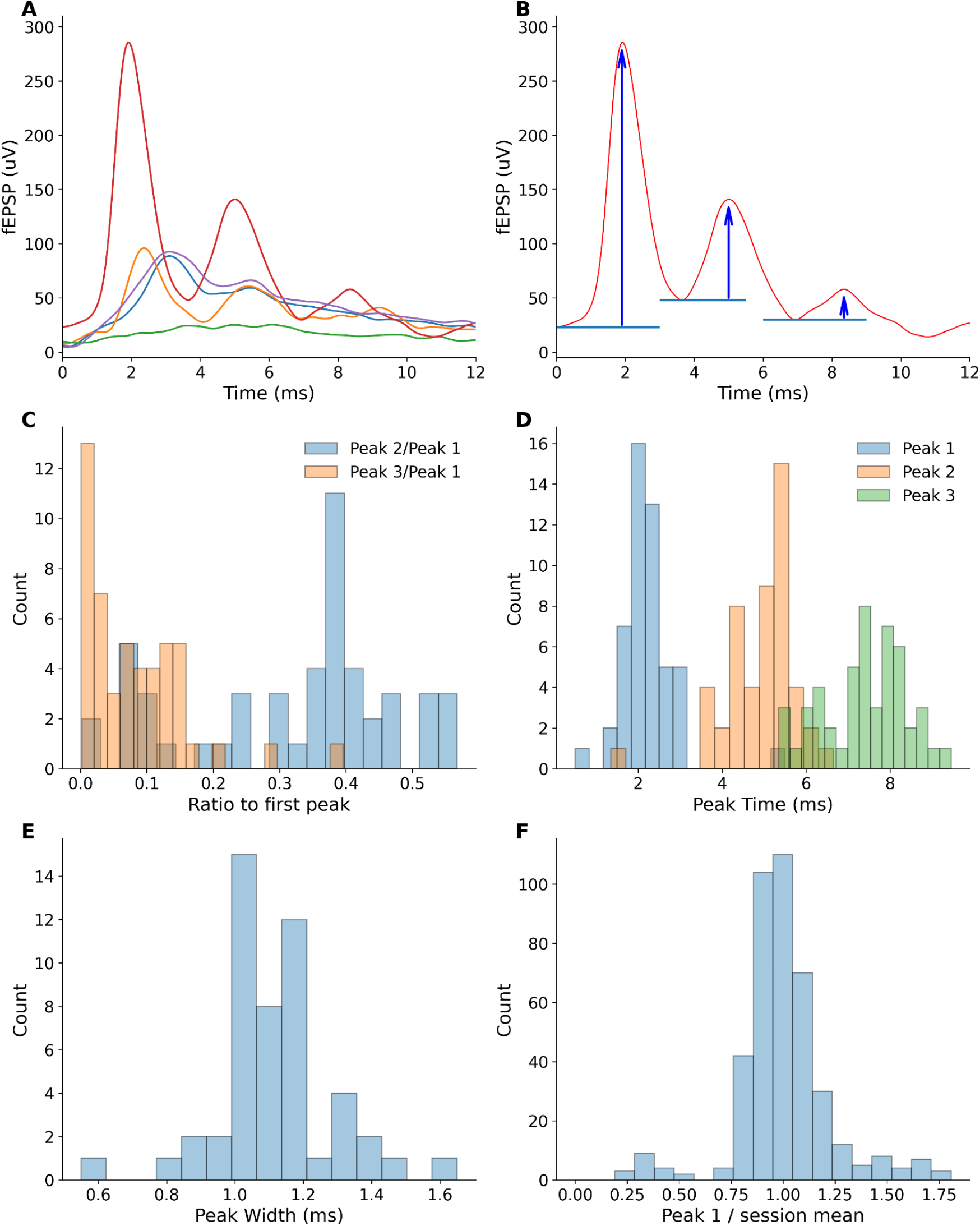
Peak distributions in CA3 field recordings to estimate occurrence of multiple spikes. A: Five sample traces, exhibiting different amounts of ‘ringing’, which indicates multiple spikes. B. Methodology for computing peak heights, illustrated on the trace with the largest ringing response. For each peak we computed the fEPSP elevation with respect to the immediately preceding valley. C. Peaks decline rapidly. The median of the second peak was ∼40% of the first peak, but the third peak was < 5%. D. Peak times following optical pulse (of 2 ms). The third peak occurred within <10ms. E: fEPSP Peak Width distribution centred around 1.2 ms, but no peak was wider than 1.6 ms, suggesting tight synchrony in case multiple CA3 neurons were spiking. Each count is for the mean width per session, including 5 and 15 square cases. F: fEPSP per-trial normalized peak height distribution is clustered around 1.0. In each case the fEPSP height of the first peak in each trial was normalized to the mean of the first fEPSP peaks in that session. Plot includes 5 and 15 square data.

**Figure 4-figure supplement 1:**
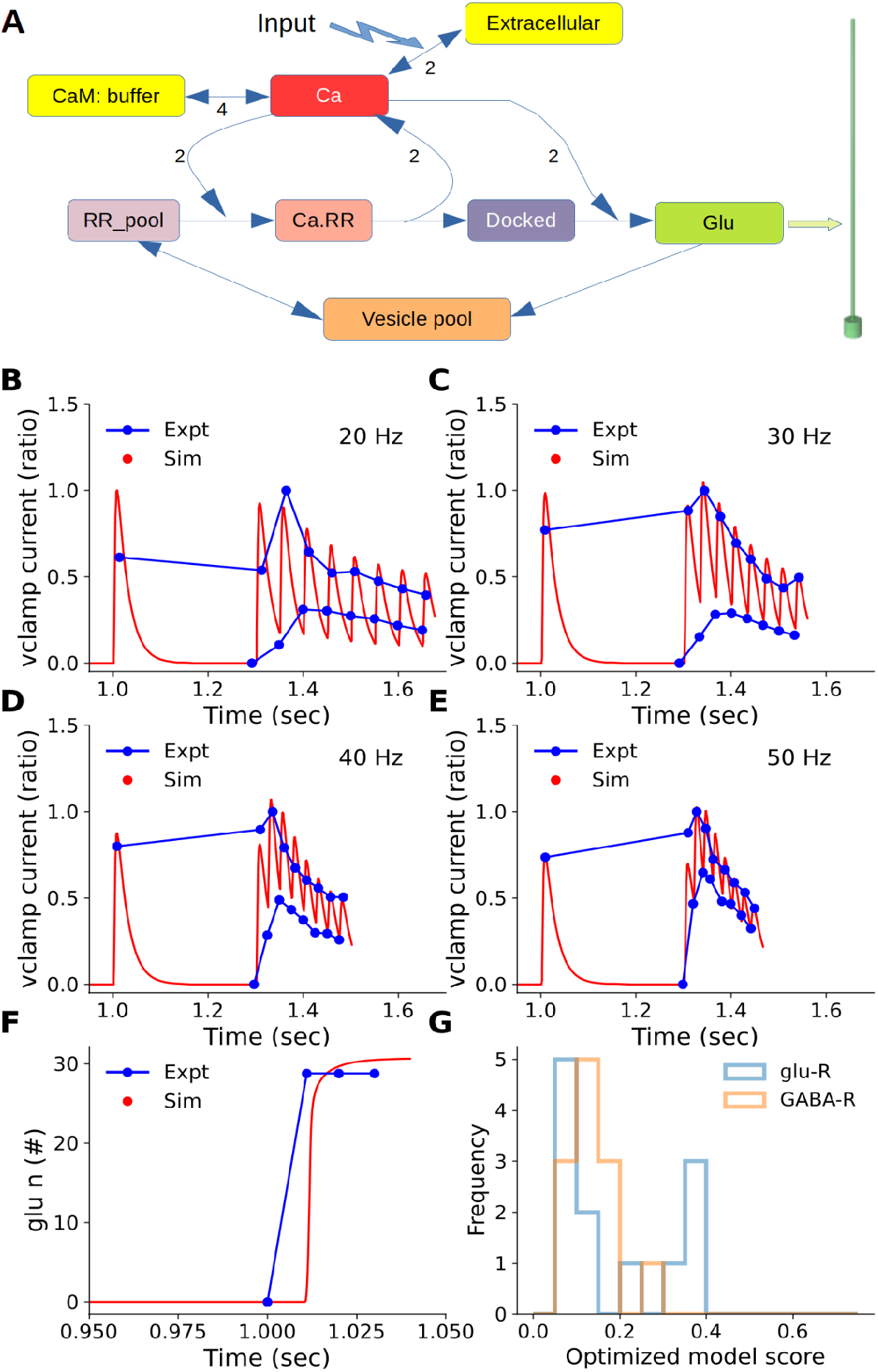
Fitting the presynaptic kinetic model to EPSC and IPSC data, for cell 7492. A: Model of reaction system in each bouton. Numbers next to reaction arrows indicate stoichiometry. Synaptic input triggers a reaction causing extracellular Ca2+ to enter the bouton, and a pump removes the Ca. CaM buffers the Ca2+. Ca binds/unbinds from successive stages of the readily releasable (RR) pool of vesicles till they are docked, at which point the final Ca-binding step causes synaptic release. The released neurotransmitter (Glu for excitatory synapses, GABA for inhibitory) opens a ligand-gated receptor channel on the postsynaptic CA1 neuron, which is held voltage-clamped.(green arrow). The reaction scheme was identical for Glutamatergic and GABAergic synapses, but the rates were different to fit to the voltage-clamp recordings. B-E: EPSC fits for 20, 30, 40 and 50 Hz, respectively. Blue traces are experimental peaks and valleys averaged over all repeats and all 5-square patterns. Red traces are simulated EPSCs. F: Cumulative release curve for glutamate vesicles. Blue: expected transmitter release completion within 10 ms based on our observations of rapid postsynaptic responses from Figure 1 H,K. Red: simulated release. G: Distribution of model fitting scores for all cells and for glu and GABA receptors. Scores are normalised RMS differences between experimental and simulated points, as in panels B-F. Any score below 0.3 is a good fit.

**Figure 4-figure supplement 2.**
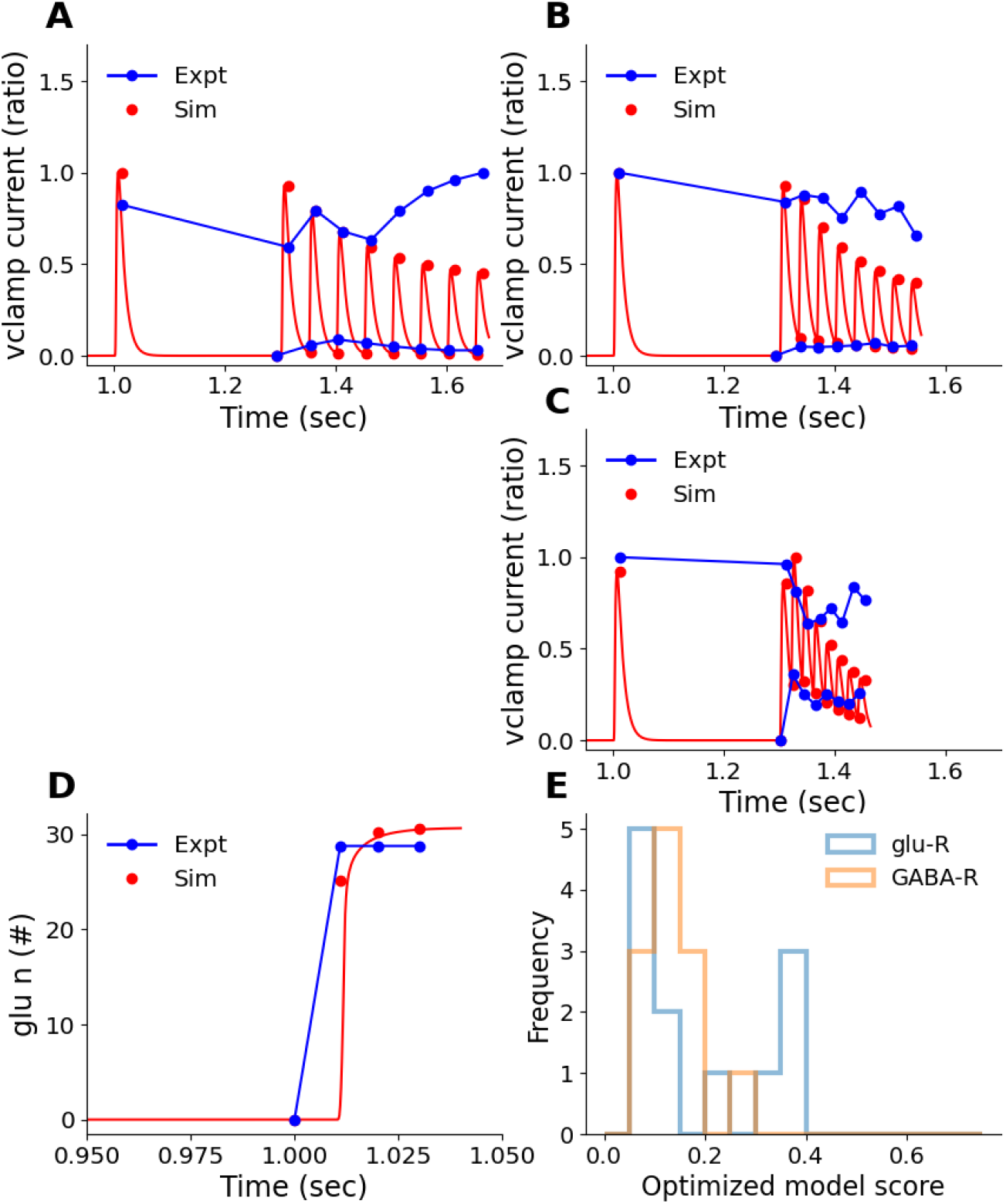
Example model fits for cell 6201. Here, the 40 Hz data was not recorded. A, B, C: EPSC fits for 20, 30, and 50 Hz, respectively. Blue traces are experimental peaks and valleys averaged over all repeats and all 15-square patterns for neuron 6201. Red traces are simulated rectified EPSCs. Note that the experimental traces are noisy hence the fits miss some of the peaks. D: Cumulative release curve for glutamate vesicles. Blue: target release complete within 10 ms. Red: simulated release. E: Distribution of model fitting scores for all cells and for glu and GABA receptors. Scores are normalized RMS differences between experimental and simulated points, as in panels A-D. Any score below 0.3 is a good fit.

**Figure 4-figure supplement 3.**
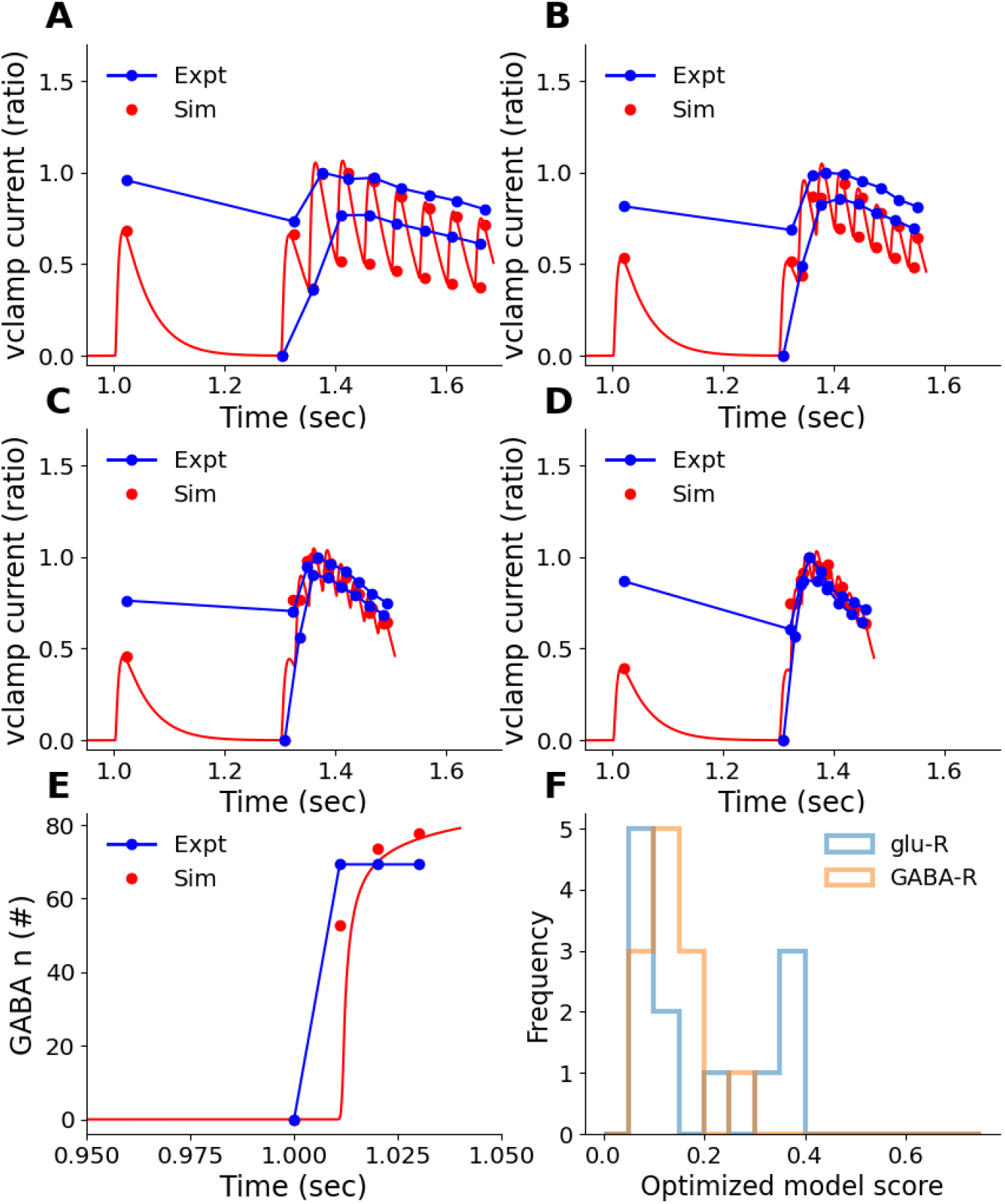
Example model fits for Inhibitory synapses on cell 7492. A, B, C, D: IPSC fits for 20, 30, 40, and 50 Hz, respectively. Blue traces are experimental peaks and valleys averaged over all repeats and all 15-square patterns for neuron 7492. Red traces are simulated IPSCs. Note that the experimental traces are noisy hence the fits miss some of the peaks. E: Cumulative release curve for GABA vesicles. Blue: target release complete within 10 ms. Red: simulated release. F: Distribution of model fitting scores for all cells and for glu and GABA receptors. Scores are normalized RMS differences between experimental and simulated points, as in panels A-D. Any score below 0.3 is a good fit.

**Figure 4-figure supplement 4.**
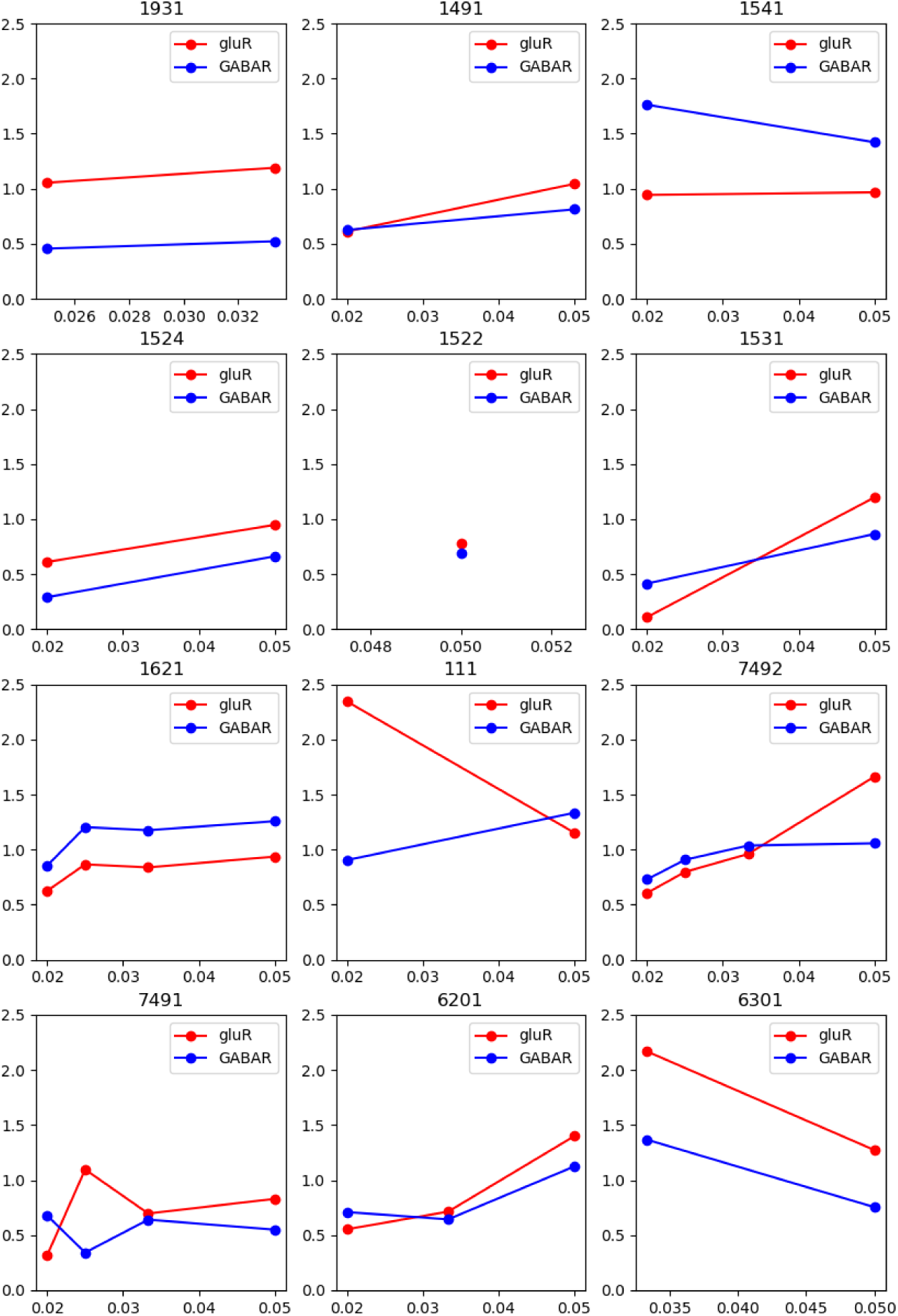
STP profiles for each of the 12 voltage-clamp recordings with the pulse-train stimulus. Some cells were not held long enough to complete all four frequencies.

**Figure 4-figure supplement 5.**
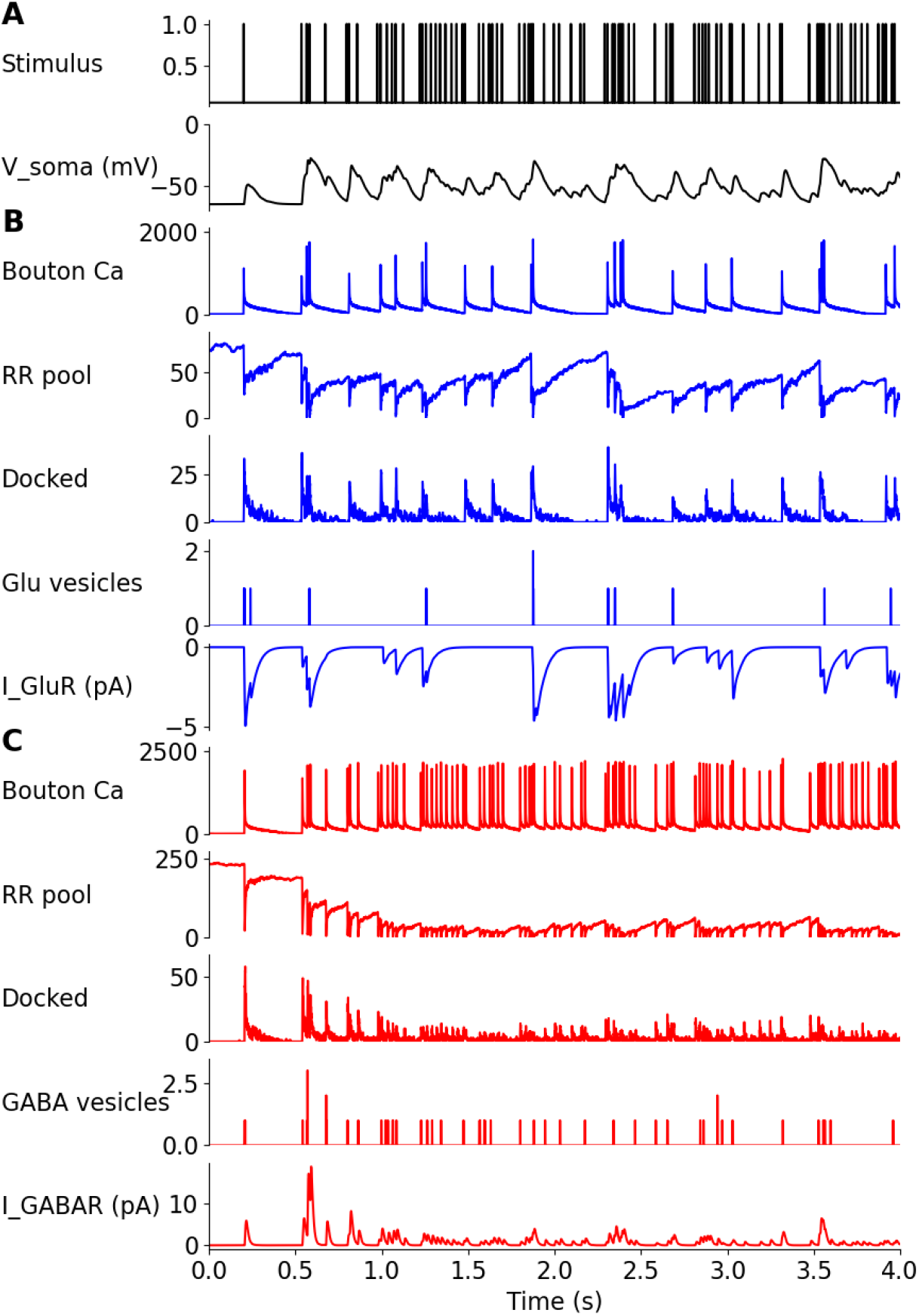
Presynaptic signalling model dynamics. All quantities are units of numbers of molecules per bouton unless otherwise stated. Signalling calculations are stochastic. Note that these are only two representative active synapses from among the 100 excitatory and 200 inhibitory synapses on the cell, and there are other synapses with distinct activity. Currents are reported as total currents for all synapses of the specified type. A: Optical stimulus train and corresponding somatic potential. B: Glutamatergic bouton signalling. C. GABAergic bouton signalling.

**Figure 4-figure supplement 6.**
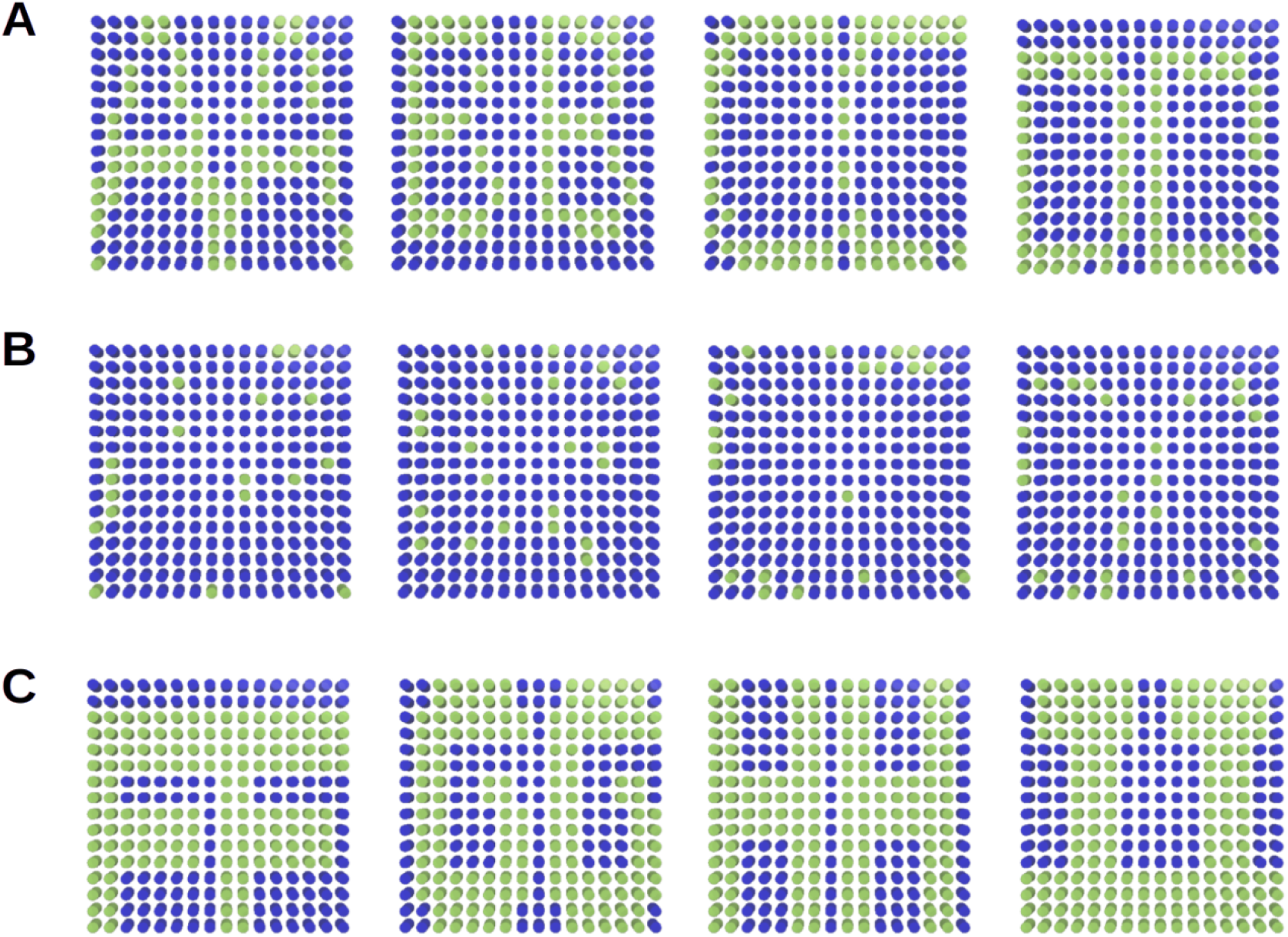
Stimulus patterns used in simulations to activate the CA3 layer. Green spots are on, blue are off. For illustration each illuminated spot is represented here as a circle, though in the experiment the shape is rectangular. A: Dense patterns used as equivalents to 5-square experimental stimuli, with 12.5% of spots set to zero, resulting in 78±2 spots per pattern. Overlap between successive patterns is 31, 23, and 37 spots. B. Reference sparse patterns to map to 5 square stimuli, with 75% of spots set to zero, resulting in 21±3 spots per pattern. Overlap between successive patterns is 1,1, and 5 spots. C. Simulation implementation of 15-square patterns, with 144, 152, 152, and 160 spots per pattern. Overlap is 68, 80 and 88 spots..

**Figure 4-figure supplement 7.**
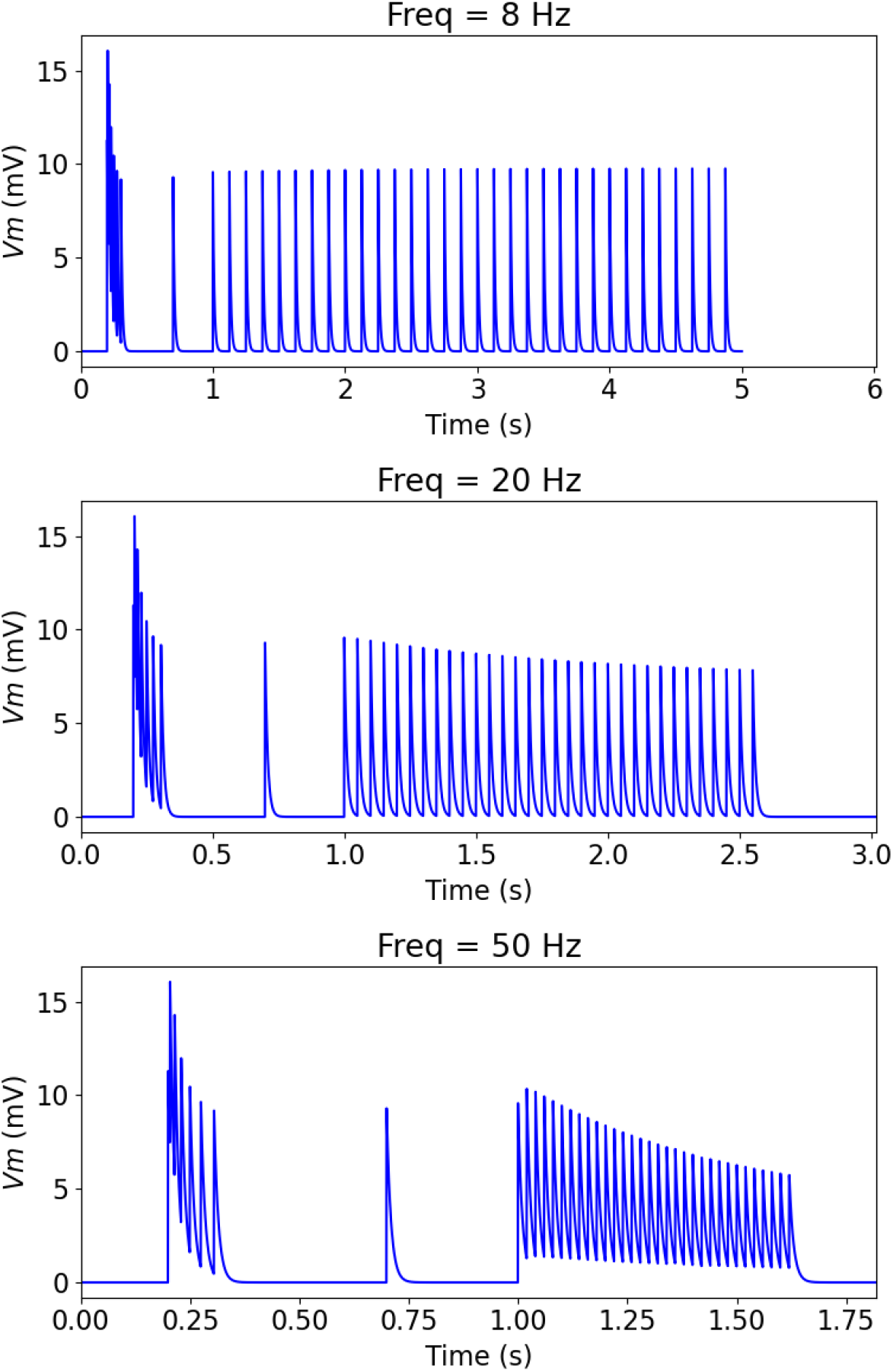
Time course of charging and desensitisation of optically stimulated CA3 neuron model. At 0.2 seconds we deliver a burst with inter-pulse intervals of 5, 10, 15, 20, 25, 30 ms. At t=0.7 seconds we deliver a single pulse. At t=1 seconds we deliver a train of 32 pulses at the frequencies of 8, 20 and 50 Hz as indicated above each panel.

**Figure 4-figure supplement 8.**
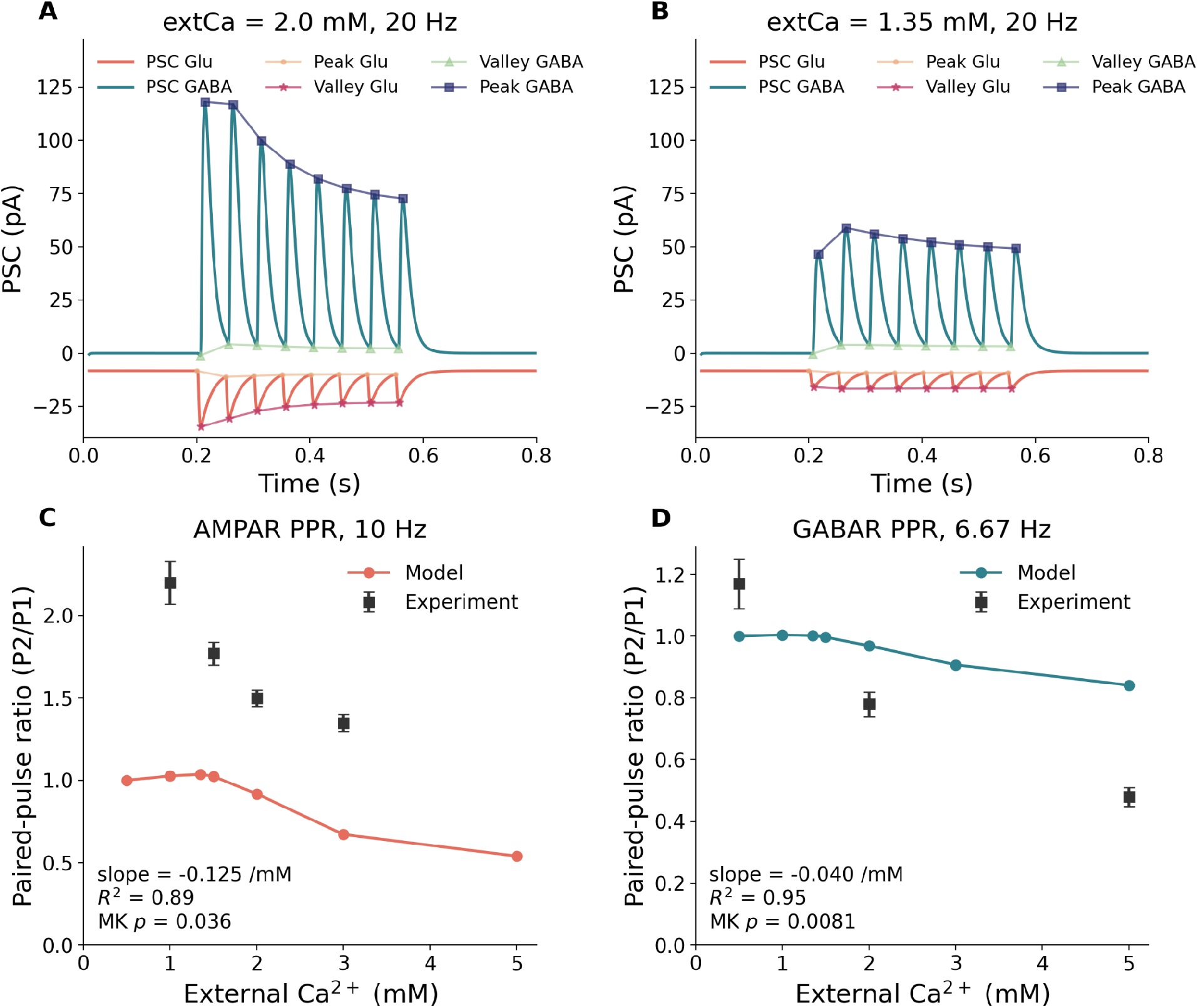
Synaptic STP responses at different levels of external calcium. A: Postsynaptic currents for Glu and GABA synapses for the reference model (2.0 mM extCa) for an 8-pulse burst at 20 Hz. B: Same as A for reduced extCa (1.35mM). Both for GluR and GABAR currents the amplitude decreases but the second peak decline converts into a small increase. C: PPR is larger at Glu synapses at low external Ca2+ for pulse interval of 100ms. Experimental data points from (Blatow et al. 2003), who used 2 week mice, 22-24 degrees C. Simulated for reference 33 degrees C model. The simulated PPR significantly declines with increasing external Ca^2+^ by the Mann-Kendall test (p=0.036). D: PPR is again larger for GABA synapses at low external Ca2+ for a pulse interval of 150ms (Mann-Kendall p=0.008). Experimental data points from (Kravchenko et al. 2006), who used 3 week primary hippocampal culture from mice at 20-23 degrees C.

**Figure 6-figure supplement 1.**
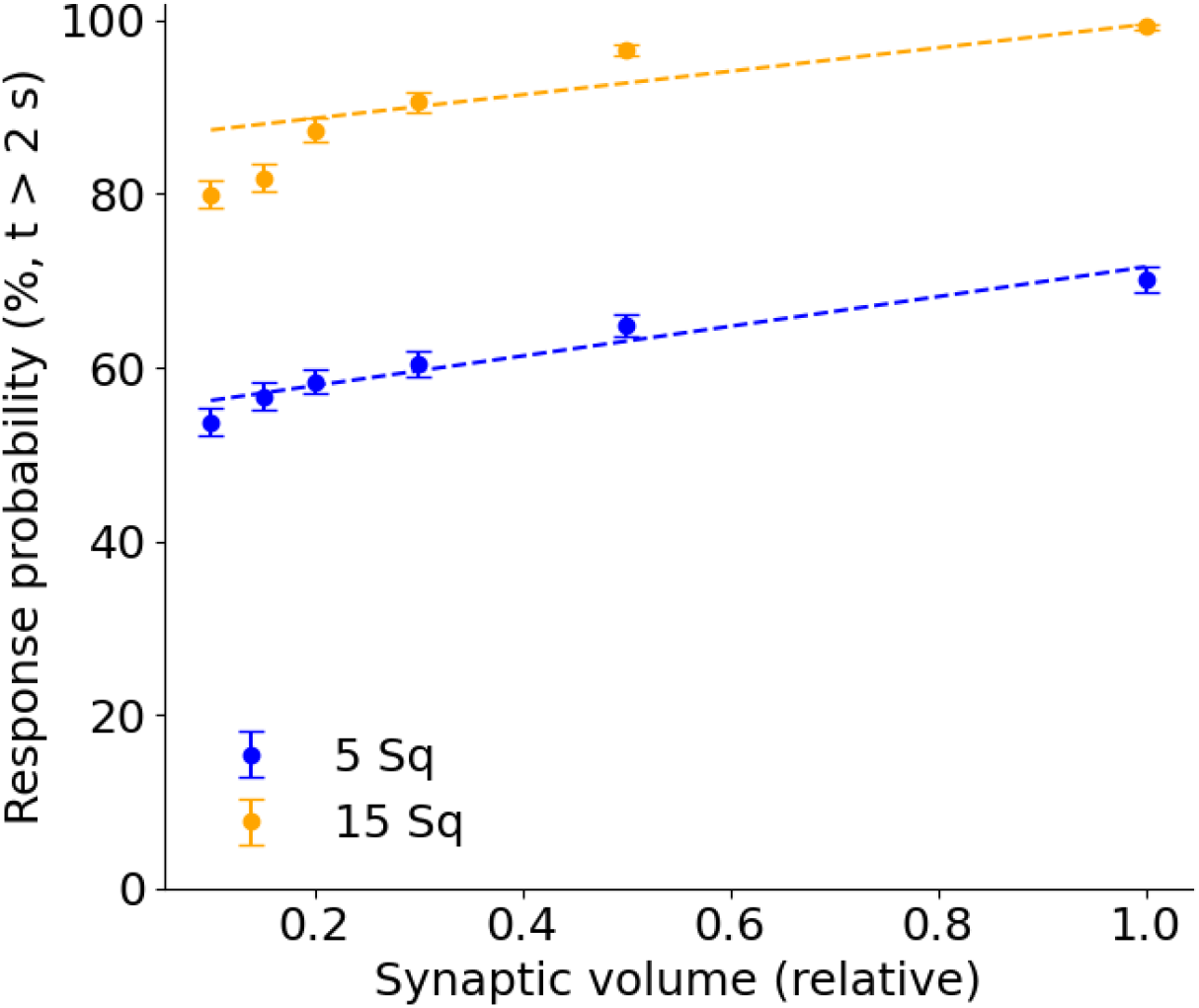
EPSP response probability rises with larger presynaptic volume. Reference volume used in simulations is scaled by 0.2, which corresponds to a synaptic volume of 0.0205±0.0008 femtolitres.

**Figure 6-figure supplement 2.**
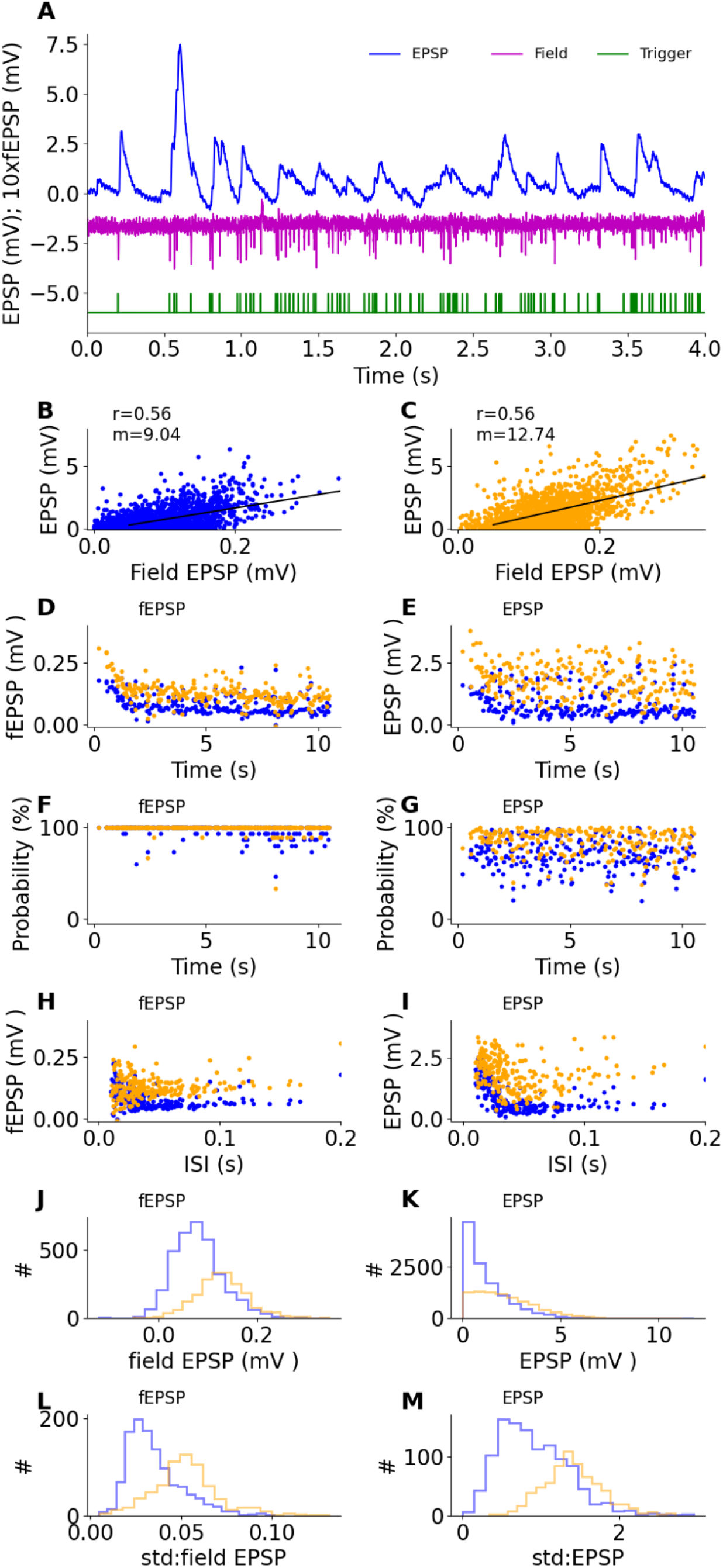
Response distributions for Poisson pulse train: field and EPSP data. A: Sample data. Blue: Raw EPSP data, maroon: field potentials, green: stimulus pulse trigger. B-M: 5 square pattern responses in blue, 15-square pattern responses in orange. B, C: Scatter plots of EPSP vs fEPSP for 5 and 15 square input, respectively. Linear regression fit is plotted in black. P-value < 0.001 in both cases. D: Field EPSP as a function of time. E: EPSP as a function of time. F. Probability that a pulse will elicit an fEPSP peak. This is almost 100% reliable. G. Probability that a pulse will elicit an EPSP. Note that the initial ∼2 seconds of the train have fewer events of low probability. H: Field EPSP as a function of inter-stimulus interval (ISI) for the pair of stimuli immediately preceding the fEPSP. I: EPSP as a function of ISI. Note that, unlike panel H, short ISIs elicit larger EPSPs. J: Distribution of field EPSP responses. K: Distribution of EPSP responses. This differs qualitatively from panel J, except that in both cases, the 15-square distributions are shifted over to the right. L: distribution of standard deviation in field EPSP for successive trials of each given pattern. M: Distribution of standard deviation in EPSP for successive trials for each pattern.

**Figure 6-figure supplement 3.**
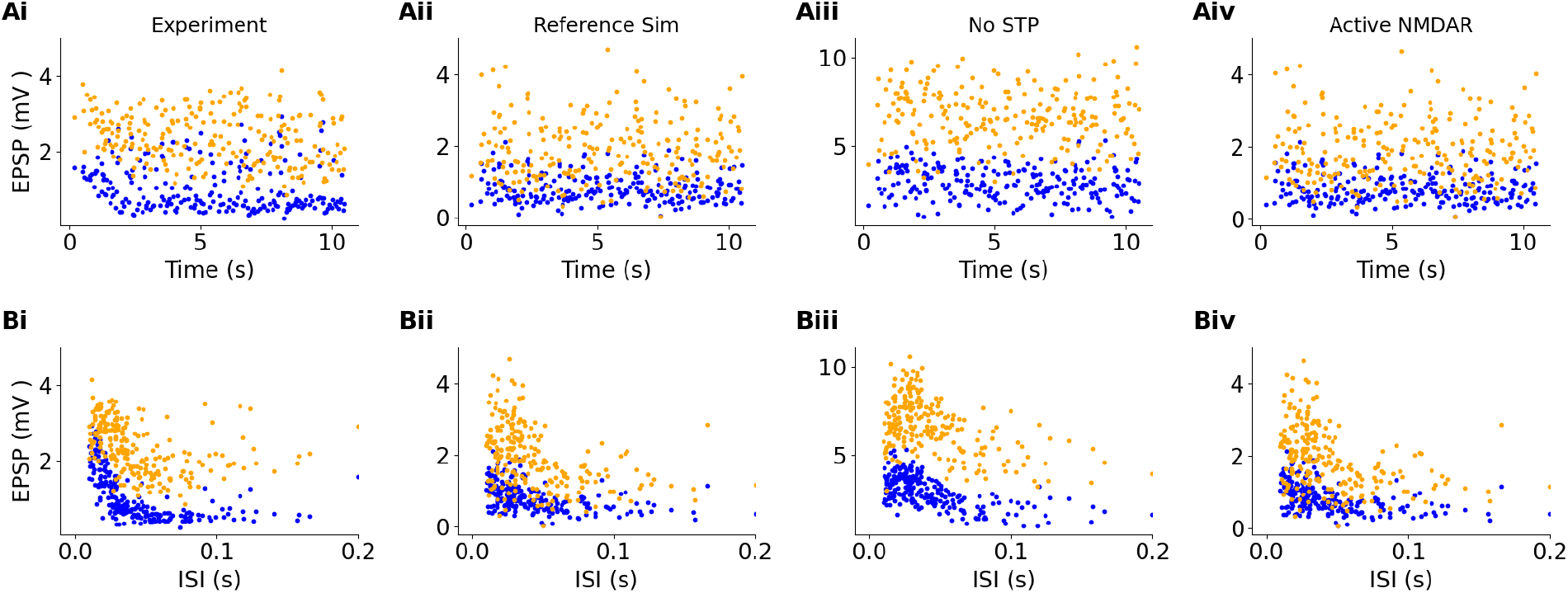
Contributions to EPSP decline with time and ISI. Yellow dots: 15 square stimuli, blue dots: 5 square stimuli. A: Contribution of STP to the initial dip in EPSPs. Ai: Experimental data. Note the decline in EPSP from t = 0 to t ∼2 seconds. Aii. Reference simulated model has a similar dip in EPSP. Aiii. Simulated model with STP disabled. In this case the 5 square case has a dip in EPSP, but not the 15 square. Aiv: Incorporation of NMDAR activity has negligible effect because the potential is well below that for release of magnesium block. B: Contributions to the dependence of EPSP on ISI. Yellow dots: 15 square stimuli, blue dots: 5 square stimuli. Bi: Experiment. There is a sharp decline, tau_5Sq = 0.3 to 18ms, tau_15Sq=8ms to 1s. Bii: Reference model. Tau_5sq=34.3±6.94ms, Tau_15sq=49.7±12.5ms. Biii: Model without STP in either GluR or GABAR synapses. Tau_5sq=58.7±9.8ms, Tau_15sq=117±35ms. The ISI dependence remains, though the time-course is slower in the absence of STP. Note that the response amplitudes are larger in the absence of STP, suggesting that presynaptic depression reduces EPSP amplitudes in the reference but not non-STP model. Biv: Model without NMDA receptors. The response is very similar to the reference response, suggesting that most of the current is carried by the Glu receptors, and the NMDA receptor opening does not lead to amplification of EPSP at short ISIs.

**Figure 7-figure supplement 1.**
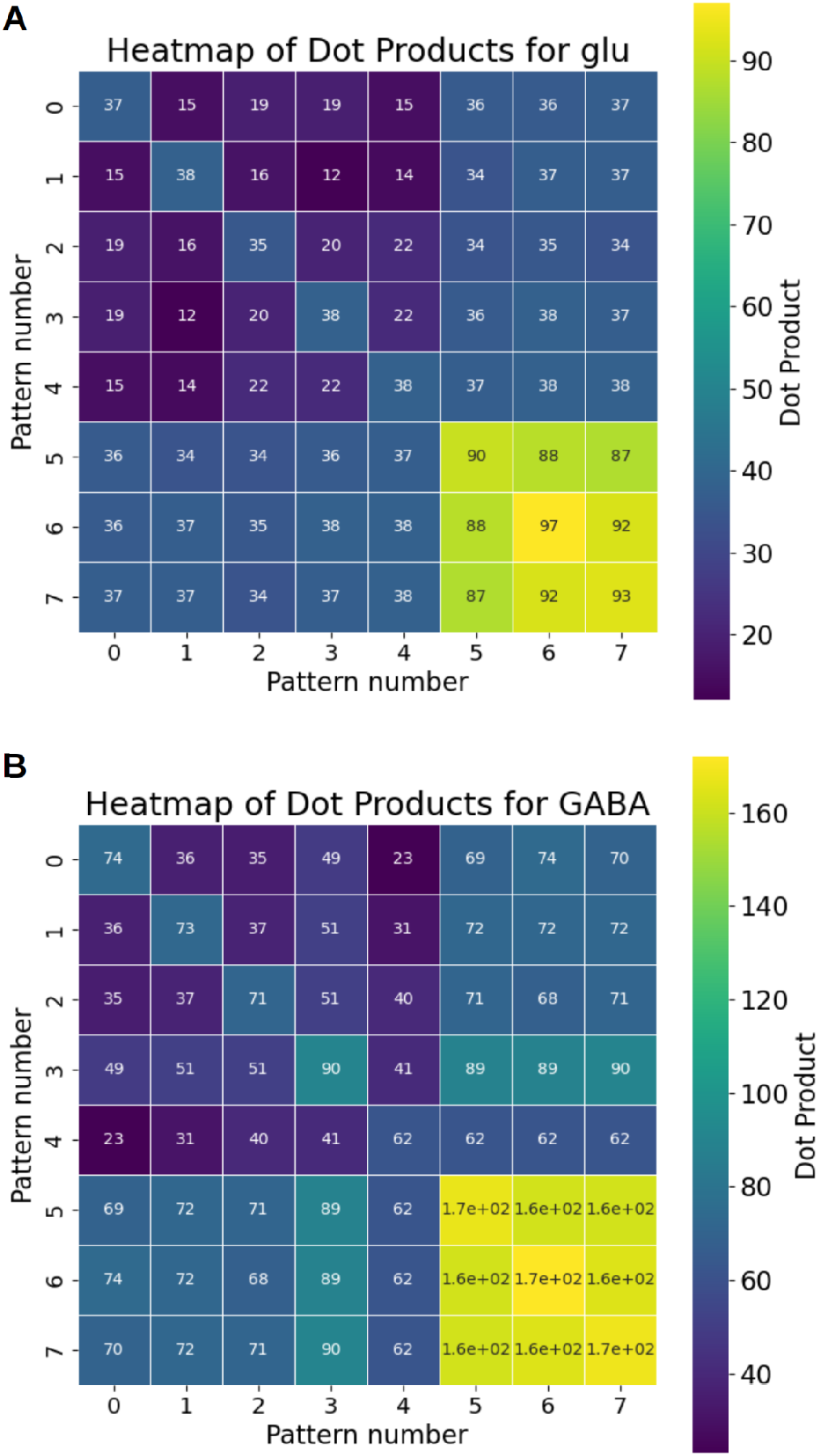
Overlap among synapses stimulated by each of 8 stimulus patterns. Patterns 0 through 4 are 5-square patterns, and patterns 5,6,7 are 15-square patterns. Diagonals indicate self-overlap, and are therefore the count of the number of stimulated synapses. A: Glutamatergic synapses. B: GABAergic synapses.

**Figure 7-figure supplement 2.**
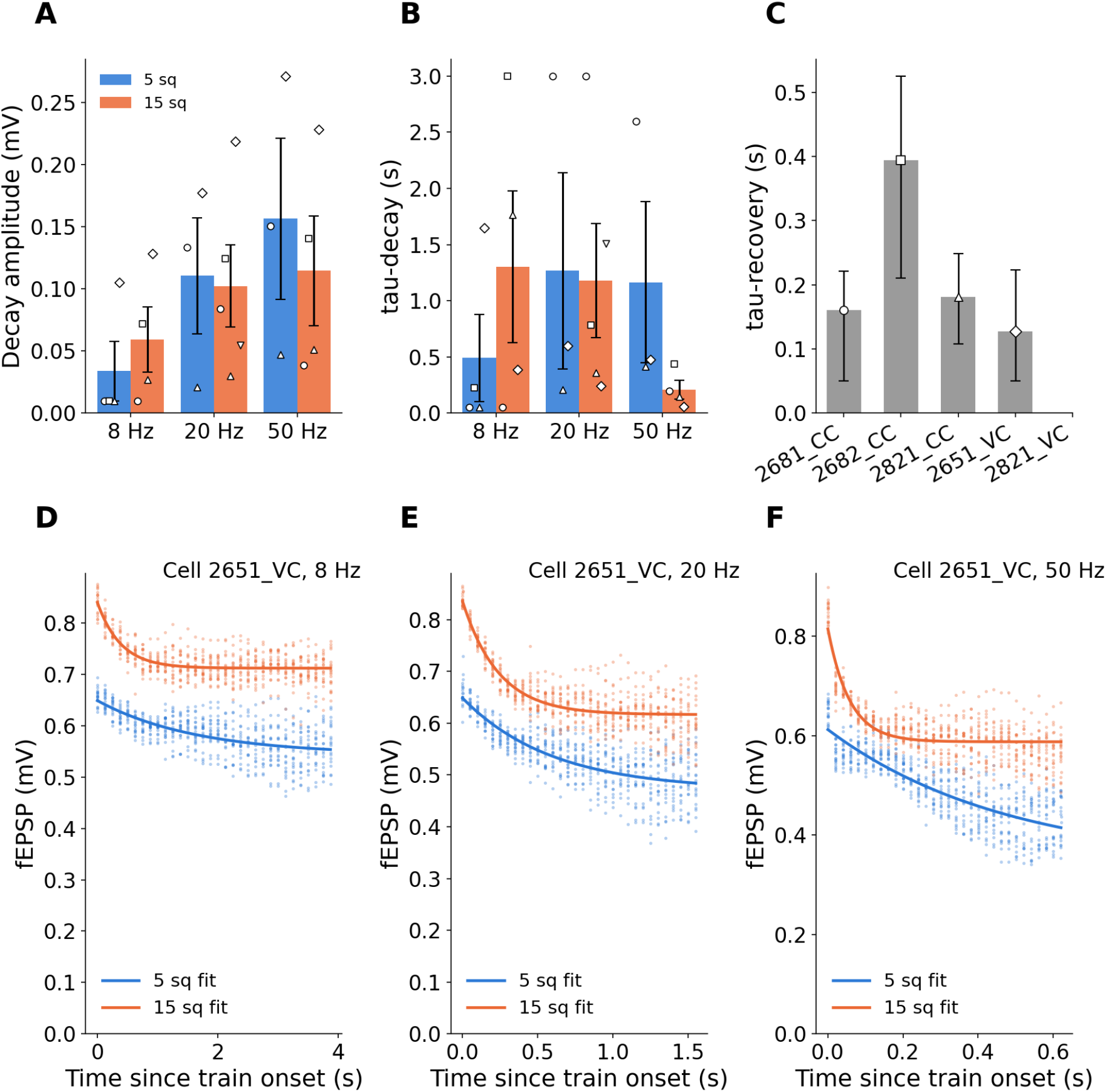
Field EPSP decay over multiple pulses in the 32-pulse train used in mismatch detection experiments. A: Decay amplitudes at different stimulus frequencies. The decay amplitude did not depend on the number of squares (Wilcoxon W=10, p = 0.58) but varied with frequency (Friedman chi^2^ = 8.27, p=0.016). The frequency dependence is expected for channelRhodopsin desensitization. B. Decay time-course means at different stimulus frequencies. Upper decay time was clipped to 3 seconds as this was nearly the total duration of the pulse train. There was no dependence on either frequency (Friedman chi^2^ = 2.0, p = 0.37) or on numSq (Wilcoxon W=18, p=0.375). C: Estimates of ChR2 recovery time, using ChR2 desensitization model in Methods and Figure 4-figure supplement 7. The estimates require model-fitting to decay amplitude, decay tau, over multiple frequencies and numSq, so only those cells which met these criteria are included in this analysis. D, E, F: Scatter plots and fits for field recordings in experiment 2651_VC, which had the largest decay amplitude. D: 8Hz pattern repeat rate: 5 sq: Decay amplitude A=0.105 mV, tau=1.649 s. 15 sq: A=0.128 mV, tau=0.390 s. E: 20 Hz repeats. 5 sq: A=0.178 mV, tau=0.598 s. 15 sq: A=0.219 mV, tau=0.241 s. F: 50 Hz repeats. 5 sq: A=0.271 mV, tau=0.477 s. 15 sq: A=0.228 mV, tau=0.058 s. Note that the fitted decay parameter A is for steady-state, which may be for a train duration longer than 32 pulses, see panel F, 5sq case where the actual decline over 32 pulses is 0.18 mV.

**Figure 7-figure supplement 3.**
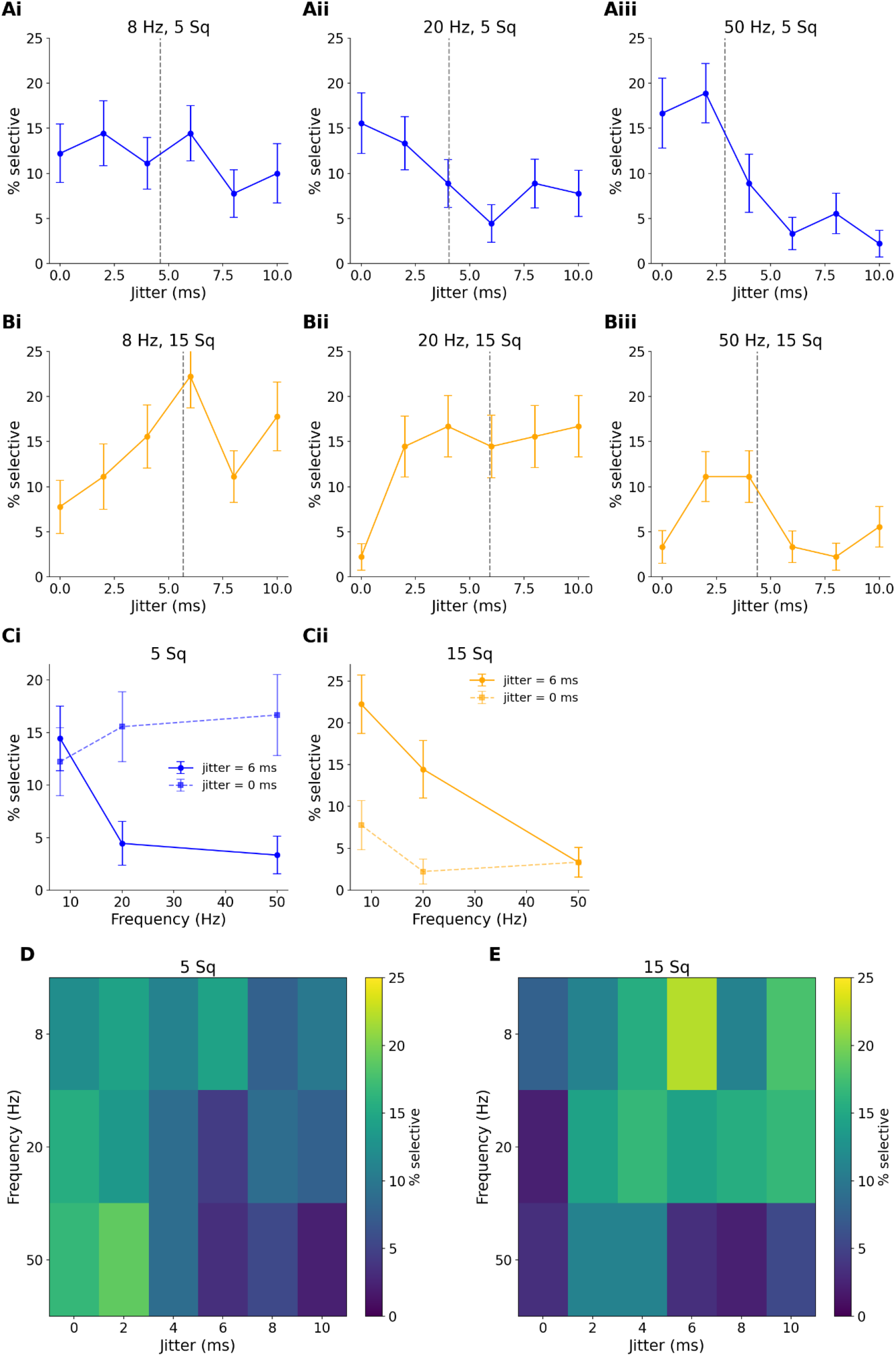
Dependence of mismatch selectivity on width of distribution of jitter in stimulus timings. Jitter was sampled from a gaussian of the specified jitter range, truncated between 0ms and 20ms. Jitter was sampled independently for each stimulating CA3 neuron spike on each trial and each repeat. A, B: Selectivity depends on jitter range and stimulus repeat frequency. Vertical dashed lines are centroids of distributions. A: 5-square stimulus. B: 15 square stimulus. Note that at all frequencies the centroid for 15 Sq stimuli is always to the right of the 5 Sq stimulus. Note also that the centroid shifts to the left for the 50Hz stimulus, both for 5 and 15 Sq. C: Dependence of selectivity on frequency, for a uniform jitter of 6 ms, as used in the main Figure 6. In both cases the selectivity drops at 50Hz, suggesting that when the jitter time becomes a large fraction of the repeat cycle time, selectivity is impaired. D, E: Heatmaps to show the profile of selectivity as a function of frequency and jitter.

**Figure 9-figure supplement 1.**
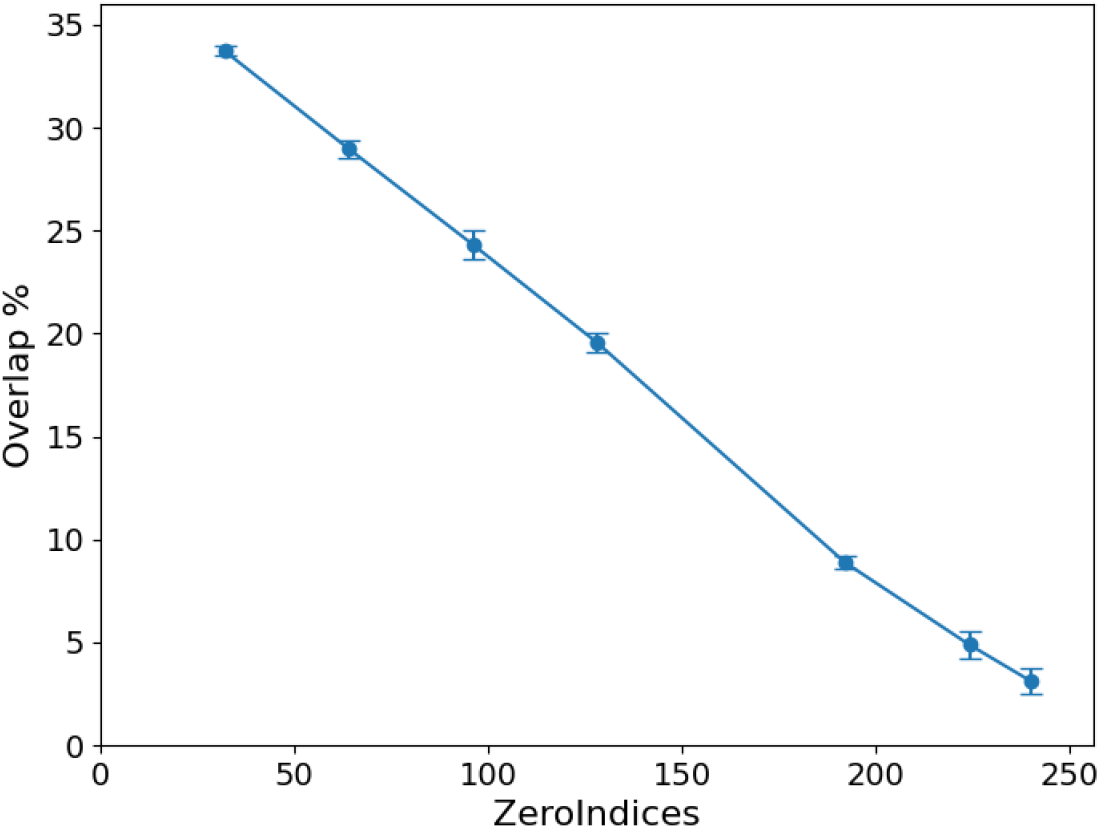
Overlap between CA1 synapses following 15 square pattern declines with the model construction parameter ZeroIndices. ZeroIndices is the number of CA3 inputs set to zero, out of a maximum of 256.

