## Supplementary Tables for Parameters. for "Single neurons detect spatiotemporal activity transitions through STP and EI imbalance"

Supplementary Methods: Model Parameters

Aditya Asopa, Upinder Singh Bhalla

### 1 Chemical model for STP synapses

#### 1.1 GABA Boutons

**Table 1.1A: Reactions for GABA boutons**

| **Reaction** | $\mathbf{K_f}$ | $\mathbf{K_b}$ |
|---|---|---|
| $2\ Ca \rightleftharpoons 2\ Ca_{ext}$ | $21.238\ \mu M^{-1} \cdot s^{-1}$ | $6.9463\ \mu M^{-1} \cdot s^{-1}$ |
| GABA $\rightleftharpoons$ 2 Ca + vesicle_pool | $38544\ s^{-1}$ | $0\ \mu M^{-2} \cdot s^{-1}$ |
| vesicle_pool $\rightleftharpoons$ RR_pool | $2.6639\ s^{-1}$ | $2.6639\ s^{-1}$ |
| 2 Ca + Docked $\rightleftharpoons$ GABA | $0.52216\ \mu M^{-2} \cdot s^{-1}$ | $0\ s^{-1}$ |
| 2 Ca + RR_pool $\rightleftharpoons$ $Ca_{RR}$ | $16.017\ \mu M^{-2} \cdot s^{-1}$ | $3.349\ s^{-1}$ |
| $Ca_{RR}$ $\rightleftharpoons$ 2 Ca + Docked | $341.59\ s^{-1}$ | $0\ \mu M^{-2} \cdot s^{-1}$ |
| Docked $\rightleftharpoons$ RR_pool | $335.88\ s^{-1}$ | $0\ s^{-1}$ |
| 4 Ca + Buffer $\rightleftharpoons$ $Ca_4$.Buffer | $335.88\ \mu M^{-4} \cdot s^{-1}$ | $1000\ s^{-1}$ |

**Table 1.1B: Pools for GABA boutons**

| **Name** | **Initial Concentration** | **Buffered** |
|---|---|---|
| Ca | $0.08\ \mu M$ | 0 |
| $Ca_{ext}$ | $0.08\ \mu M$ | 1 |
| RR_pool | $3\ \mu M$ | 0 |
| vesicle_pool | $3.1699\ \mu M$ | 1 |
| Docked | $0\ \mu M$ | 0 |
| $Ca_{RR}$ | $0\ \mu M$ | 0 |
| GABA | $0\ \mu M$ | 0 |
| Buffer | $3.4727\ \mu M$ | 0 |
| $Ca_4$.Buffer | $0\ \mu M$ | 0 |

#### 1.2 Glutamate Boutons

**Table 1.2A: Reactions for Glu boutons**

| **Reaction** | $\mathbf{K_f}$ | $\mathbf{K_b}$ |
|---|---|---|
| $2\ \mathrm{Ca} \rightleftharpoons 2\ \mathrm{Ca_{ext}}$ | $47.118\ \mu\mathrm{M}^{-1} \cdot \mathrm{s}^{-1}$ | $15.889\ \mu\mathrm{M}^{-1} \cdot \mathrm{s}^{-1}$ |
| glu ⇌ 2 Ca + vesicle_pool | $74957\ \mathrm{s}^{-1}$ | $0\ \mu\mathrm{M}^{-2} \cdot \mathrm{s}^{-1}$ |
| vesicle_pool ⇌ RR_pool | $2.6147\ \mathrm{s}^{-1}$ | $2.6147\ \mathrm{s}^{-1}$ |
| 2 Ca + Docked ⇌ glu | $0.62741\ \mu\mathrm{M}^{-2} \cdot \mathrm{s}^{-1}$ | $0\ \mathrm{s}^{-1}$ |
| 2 Ca + RR_pool ⇌ $\mathrm{Ca_{RR}}$ | $0.99392\ \mu\mathrm{M}^{-2} \cdot \mathrm{s}^{-1}$ | $0.17877\ \mathrm{s}^{-1}$ |
| $\mathrm{Ca_{RR}}$ ⇌ 2 Ca + Docked | $1986.1\ \mathrm{s}^{-1}$ | $0\ \mu\mathrm{M}^{-2} \cdot \mathrm{s}^{-1}$ |
| Docked ⇌ RR_pool | $127.73\ \mathrm{s}^{-1}$ | $0\ \mathrm{s}^{-1}$ |
| 4 Ca + Buffer ⇌ $\mathrm{Ca_4}$.Buffer | $127.73\ \mu\mathrm{M}^{-4} \cdot \mathrm{s}^{-1}$ | $1000\ \mathrm{s}^{-1}$ |

**Table 1.2B: Pools for Glutamate boutons**

| **Name** | **Initial Concentration** | **Buffered** |
|---|---|---|
| Ca | $0.08\ \mu\mathrm{M}$ | 0 |
| $\mathrm{Ca_{ext}}$ | $0.08\ \mu\mathrm{M}$ | 1 |
| RR_pool | $0.3\ \mu\mathrm{M}$ | 0 |
| vesicle_pool | $0.872\ \mu\mathrm{M}$ | 1 |
| glu | $0\ \mu\mathrm{M}$ | 0 |
| Docked | $0\ \mu\mathrm{M}$ | 0 |
| $\mathrm{Ca_{RR}}$ | $0\ \mu\mathrm{M}$ | 0 |
| Buffer | $4.0953\ \mu\mathrm{M}$ | 0 |
| $\mathrm{Ca_4}$.Buffer | $0\ \mu\mathrm{M}$ | 0 |

### 2 Chemical model for non-STP synapses

#### 2.1 GABA boutons

**Table 2.1A: Reactions for GABA boutons**

| Reaction | $K_f$ | $K_b$ |
|---|---|---|
| $2\ Ca \rightleftharpoons 2\ Ca_{ext}$ | $21.238\ \mu M^{-1} \cdot s^{-1}$ | $6.9463\ \mu M^{-1} \cdot s^{-1}$ |
| $GABA \rightleftharpoons 2\ Ca + vesicle\_pool$ | $38544\ s^{-1}$ | $0\ \mu M^{-2} \cdot s^{-1}$ |
| $2\ Ca + Docked \rightleftharpoons GABA$ | $0.52216\ \mu M^{-2} \cdot s^{-1}$ | $0\ s^{-1}$ |

**Table 2.1B: Pools for GABA boutons**

| Name | Initial Concentration | Buffered |
|---|---|---|
| Ca | 0.08 $\mu$M | 0 |
| $Ca_{ext}$ | 0.08 $\mu$M | 1 |
| vesicle_pool | 3.1699 $\mu$M | 1 |
| Docked | 0.5 $\mu$M | 1 |
| GABA | 0 $\mu$M | 0 |

#### 2.2 Glutamate boutons

**Table 2.2A: Reactions for glu boutons**

| Reaction | $K_f$ | $K_b$ |
|---|---|---|
| $2\ Ca \rightleftharpoons 2\ Ca_{ext}$ | $47.118\ \mu M^{-1} \cdot s^{-1}$ | $15.889\ \mu M^{-1} \cdot s^{-1}$ |
| $glu \rightleftharpoons 2\ Ca + vesicle\_pool$ | $74957\ s^{-1}$ | $0\ \mu M^{-2} \cdot s^{-1}$ |
| $2\ Ca + Docked \rightleftharpoons glu$ | $0.62741\ \mu M^{-2} \cdot s^{-1}$ | $0\ s^{-1}$ |

**Table 2.2B: Pools for glu boutons**

| Name | Initial Concentration | Buffered |
|---|---|---|
| Ca | 0.08 $\mu$M | 0 |
| $Ca_{ext}$ | 0.08 $\mu$M | 1 |
| vesicle_pool | 0.872 $\mu$M | 1 |
| glu | 0 $\mu$M | 0 |
| Docked | 0.225 $\mu$M | 1 |

### 3 Chemical model at 37°C for STP synapses

#### 3.1 GABA Boutons

**Table 3.1A: Reactions for GABA boutons**

| **Reaction** | $\mathbf{K_f}$ | $\mathbf{K_b}$ |
|---|---|---|
| $2\ Ca \rightleftharpoons 2\ Ca_{ext}$ | $29.7\ \mu M^{-1} \cdot s^{-1}$ | $9.72\ \mu M^{-1} \cdot s^{-1}$ |
| $GABA \rightleftharpoons 2\ Ca + vesicle\_pool$ | $53962\ s^{-1}$ | $0\ \mu M^{-2} \cdot s^{-1}$ |
| $vesicle\_pool \rightleftharpoons RR\_pool$ | $3.73\ s^{-1}$ | $3.73\ s^{-1}$ |
| $2\ Ca + Docked \rightleftharpoons GABA$ | $0.731\ \mu M^{-2} \cdot s^{-1}$ | $0\ s^{-1}$ |
| $2\ Ca + RR\_pool \rightleftharpoons Ca_{RR}$ | $22.4\ \mu M^{-2} \cdot s^{-1}$ | $4.69\ s^{-1}$ |
| $Ca_{RR} \rightleftharpoons 2\ Ca + Docked$ | $478.2\ s^{-1}$ | $0\ \mu M^{-2} \cdot s^{-1}$ |
| $Docked \rightleftharpoons RR\_pool$ | $470.2\ s^{-1}$ | $0\ s^{-1}$ |
| $4\ Ca + Buffer \rightleftharpoons Ca_4.Buffer$ | $470.2\ \mu M^{-4} \cdot s^{-1}$ | $1400\ s^{-1}$ |

**Table 3.1B: Pools for GABA boutons**

| **Name** | **Initial Concentration** | **Buffered** |
|---|---|---|
| Ca | 0.08 $\mu$M | 0 |
| $Ca_{ext}$ | 0.08 $\mu$M | 1 |
| RR_pool | 3 $\mu$M | 0 |
| vesicle_pool | 3.1699 $\mu$M | 1 |
| Docked | 0 $\mu$M | 0 |
| $Ca_{RR}$ | 0 $\mu$M | 0 |
| GABA | 0 $\mu$M | 0 |
| Buffer | 3.4727 $\mu$M | 0 |
| $Ca_4$.Buffer | 0 $\mu$M | 0 |

#### 3.2 Glutamate Boutons

**Table 3.2A: Reactions for Glu boutons**

| **Reaction** | $\mathbf{K_f}$ | $\mathbf{K_b}$ |
|---|---|---|
| $2\ Ca \rightleftharpoons 2\ Ca_{ext}$ | $66\ \mu M^{-1} \cdot s^{-1}$ | $22.2\ \mu M^{-1} \cdot s^{-1}$ |
| $glu \rightleftharpoons 2\ Ca + vesicle\_pool$ | $104940\ s^{-1}$ | $0\ \mu M^{-2} \cdot s^{-1}$ |
| $vesicle\_pool \rightleftharpoons RR\_pool$ | $3.73\ s^{-1}$ | $3.73\ s^{-1}$ |
| $2\ Ca + Docked \rightleftharpoons glu$ | $0.878\ \mu M^{-2} \cdot s^{-1}$ | $0\ s^{-1}$ |
| $2\ Ca + RR\_pool \rightleftharpoons Ca_{RR}$ | $1.4\ \mu M^{-2} \cdot s^{-1}$ | $0.25\ s^{-1}$ |
| $Ca_{RR} \rightleftharpoons 2\ Ca + Docked$ | $2789.5\ s^{-1}$ | $0\ \mu M^{-2} \cdot s^{-1}$ |
| $Docked \rightleftharpoons RR\_pool$ | $178.8\ s^{-1}$ | $0\ s^{-1}$ |
| $4\ Ca + Buffer \rightleftharpoons Ca_4.Buffer$ | $178.8\ \mu M^{-4} \cdot s^{-1}$ | $1400\ s^{-1}$ |

**Table 3.2B: Pools for Glutamate boutons**

| **Name** | **Initial Concentration** | **Buffered** |
|---|---|---|
| Ca | 0.08 $\mu$M | 0 |
| $Ca_{ext}$ | 0.08 $\mu$M | 1 |
| RR_pool | 0.3 $\mu$M | 0 |
| vesicle_pool | 0.872 $\mu$M | 1 |
| glu | 0 $\mu$M | 0 |
| Docked | 0 $\mu$M | 0 |
| $Ca_{RR}$ | 0 $\mu$M | 0 |
| Buffer | 4.0953 $\mu$M | 0 |
| $Ca_4$.Buffer | 0 $\mu$M | 0 |

### 4 Electrical Model

The voltage $V$ is in mV and referenced to the resting potential. Time is in milliseconds (ms).

#### 4.1 Ion Channel Definitions

These definitions are mostly from Traub et al, 1991.

| Ion | Conductance | Reversal | Gate | $\alpha$ | $\beta$ |
|---|---|---|---|---|---|
| Na | $g_{\mathrm{Na}} = g_{\mathrm{max\_Na}} \cdot m^2 h$ | $E_{\mathrm{Na}} = 115$ mV | m-gate | $\frac{0.32(13.1-V)}{\exp\left(\frac{13.1-V}{4}\right)-1}$ | $\frac{0.28(V-40.1)}{\exp\left(\frac{V-40.1}{5}\right)-1}$ |
| | | | h-gate | $0.128\exp\left(\frac{17-V}{18}\right)$ | $\frac{4}{1+\exp\left(\frac{40-V}{5}\right)}$ |
| K | $g_{\mathrm{KDR}} = g_{\mathrm{max\_KDR}} \cdot n$ | $E_{\mathrm{KDR}} = -15$ mV | n-gate | $\frac{0.016(35.1-V)}{\exp\left(\frac{35.1-V}{5}\right)-1}$ | $0.25\exp\left(\frac{20-V}{40}\right)$ |

#### 4.2 Receptor-gated Ion Channel Conductances

**Glutamate Receptor: AMPA**

$$g_{\mathrm{GluR}} = \frac{A \cdot g_{\mathrm{maxGluR}}}{\tau_1 - \tau_2}\left(\exp\left(\frac{-t}{\tau_1}\right) - \exp\left(\frac{-t}{\tau_2}\right)\right)$$

where: A = normalization constant such that $g_{\mathrm{GluR}} = g_{\mathrm{maxGluR}}$ at peak. $\tau_1 = 2, \tau_2 = 9$

**GABA Receptor**

$$g_{\mathrm{GABAR}} = \frac{A \cdot g_{\mathrm{maxGABAR}}}{\tau_1 - \tau_2}\left(\exp\left(\frac{-t}{\tau_1}\right) - \exp\left(\frac{-t}{\tau_2}\right)\right)$$

where: A = normalization constant such that $g_{\mathrm{GABAR}} = g_{\mathrm{maxGABAR}}$ at peak. $\tau_1 = 4, \tau_2 = 9$

**Glutamate Receptor: NMDA**

$$g_{\mathrm{NMDAR}} = \frac{g_{\mathrm{maxNMDAR}}}{\tau}\exp\left(\frac{-t}{\tau}\right)\frac{K_{\mathrm{Mg}}}{K_{\mathrm{Mg}} + [\mathrm{Mg}]}$$

where: $K_{Mg} = \frac{\gamma \cdot \exp(V - E_{rest})}{\eta}$, $\quad \tau = 20, \quad \gamma = 0.28, \quad \eta = 62$

$$I_{\mathrm{NMDAR_{Ca}}} = g_{\mathrm{NMDAR}} \cdot \mathrm{Ca_{frac}} \cdot \ln\left(\frac{[\mathrm{Ca_{out}}]}{[\mathrm{Ca_{in}}]}\right) \cdot V \cdot \frac{[\mathrm{Ca_{in}}] - \varphi[\mathrm{Ca_{out}}]}{(1-\varphi)([\mathrm{Ca_{in}}] - [\mathrm{Ca_{out}}])}$$

Where: $F = 96485$ sA/mol, $\quad z = 2, \quad R = 8.314\ J/(K \cdot mol), \quad T = 300$ K, $\quad \varphi = \exp(-VFz/RT)$, $\mathrm{Ca_{frac}}$ = fraction of current carried at 0 mV by Ca = 0.02, $\quad [\mathrm{Ca_{out}}] = 1.5$ mM, $\quad [\mathrm{Ca_{in}}] = 0.08\ \mu$M

#### 4.3 Calcium Pools

$$\frac{d[\mathrm{Ca}]}{dt} = \phi\left(I_{\mathrm{Ca}} + \mathrm{Ca_{NMDA}}\right) - \frac{[\mathrm{Ca}]}{13.33}$$

where: $\phi =$

##### 4.4 Passive Properties

$R_M = 0.2\ \Omega \cdot \text{m}^2, \quad R_A = 1.0\ \Omega \cdot \text{m}, \quad C_M = 0.01\ \text{F/m}^2, \quad E_{\text{rest}} = -70\ \text{mV}$

##### 4.5 Channel Distributions

| **Channel** | **Zone** | **Distribution (Gmax, S/m$^2$)** |
|---|---|---|
| Na | Soma | 600 |
| K_DR | Soma | 360 |
| GABAR | Dendrite | 50 |
| GluR | Spine Heads | 200 |
| NMDAR | Spine Heads | 80 |
